# Nintedanib Targets KIT D816V Neoplastic Cells Derived from Induced Pluripotent Stem cells of Systemic Mastocytosis

**DOI:** 10.1101/2020.05.06.080150

**Authors:** Marcelo A. S. Toledo, Malrun Gatz, Stephanie Sontag, Karoline V. Gleixner, Gregor Eisenwort, Kristina Feldberg, Frederick Kluge, Riccardo Guareschi, Giulia Rossetti, Antonio S. Sechi, Olli M. J. Dufva, Satu M. Mustjoki, Angela Maurer, Herdit M. Schüler, Roman Goetzke, Till Braunschweig, Anne Simonowski, Jens Panse, Mohamad Jawhar, Andreas Reiter, Frank Hilberg, Peter Ettmayer, Wolfgang Wagner, Steffen Koschmieder, Tim H. Brümmendorf, Peter Valent, Nicolas Chatain, Martin Zenke

**Author notes:** To whom correspondence should be addressed: Martin Zenke, PhD, Institute for Biomedical Engineering, Department of Cell Biology, RWTH Aachen University Medical School, Pauwelsstrasse 30, 52074 Aachen, Germany, Voice (secretary). **Key Points** - Patient-derived *KIT* D816V iPSCs and CRISPR-engineered *KIT* D816V ESCs model SM disease heterogeneity and serve as drug screening platform - Nintedanib selectively targets *KIT* D816V iPSC- and ESC-derived cells and primary samples of SM patients. **Primary Scientific Category:** Myeloid neoplasia.

## Abstract

The *KIT* D816V mutation is found in more than 80% of patients with systemic mastocytosis (SM) and is key to neoplastic mast cell (MC) expansion and accumulation in affected organs. *KIT* D816V therefore represents a prime therapeutic target for SM. Here we generated a panel of patient-specific *KIT* D816V induced pluripotent stem cells (iPSCs) from patients with aggressive SM (ASM) and mast cell leukemia (MCL) to develop a patient-specific SM disease model for mechanistic and drug discovery studies. *KIT* D816V iPSCs differentiated into neoplastic hematopoietic progenitor cells and MCs with patient-specific phenotypic features, thereby reflecting the heterogeneity of the disease. CRISPR/Cas9n-engineered *KIT* D816V human embryonic stem cells (ESCs), when differentiated into hematopoietic cells, recapitulated the phenotype observed for *KIT* D816V iPSC hematopoiesis. *KIT* D816V causes constitutive activation of the KIT tyrosine kinase receptor and we exploited our iPSCs and ESCs to investigate new tyrosine kinase inhibitors targeting KIT D816V. Our study identified nintedanib as a novel KIT D816V inhibitor. Nintedanib selectively reduced the viability of iPSC-derived *KIT* D816V hematopoietic progenitor cells and MCs in the nanomolar range. Nintedanib was also active on primary samples of KIT D816V SM patients. Molecular docking studies show that nintedanib binds to the ATP binding pocket of inactive KIT D816V. Our results suggest nintedanib as a new drug candidate for KIT D816V targeted therapy of advanced SM.

## Introduction

Mastocytosis comprises a group of hematopoietic malignancies characterized by abnormal proliferation and accumulation of neoplastic mast cells (MCs) in one or multiple tissues and organs, including bone marrow, spleen, liver and skin.^1,2^ The degree of MC infiltration, number of tissues/organs involved and mutational load contribute to a heterogeneous pathology and distinct disease categories in systemic mastocytosis (SM): indolent SM (ISM), smoldering SM (SSM), aggressive SM (ASM), MC leukemia (MCL) and SM with an associated hematological disease (SM-AHD).^1,3,4^ ASM, MCL and SM-AHD represent advanced forms of SM, which are characterized by pronounced MC infiltration compromising organ function, rapid disease progression and poor prognosis. Only a few therapeutic options are available for these patients.^4–6^

Mutations in the gene encoding the KIT receptor are central to the evolution of SM and the *KIT* D816V mutation is most prevalent and identified in all SM categories.^3,6–8^ This mutation leads to constitutive activation of KIT and abnormal expansion and accumulation of MCs in affected organs.^6,8,9^ Several tyrosine kinase inhibitors (TKIs) were reported to inhibit KIT activity in neoplastic MC.^10–14^ However, *KIT* D816V confers resistance to several of these TKIs, including imatinib.^11,15^ Other TKIs, such as midostaurin, suppress the growth of KIT D816V neoplastic MCs and recently, midostaurin was approved for treatment of advanced SM.^16–18^ More recently, two further compounds targeting KIT D816V, ripretinib (DCC-2618) and avapritinib (BLU-285), were identified and are being tested in clinical trials for advanced SM.^19–21^ However, none of the available drugs can induce complete long-lasting remission in all patients and therefore, further efforts to establish preclinical models and to identify new TKIs are needed.

Primary samples of SM patients represent a scarce and highly variable cell source for preclinical studies. Moreover, MCs often represent a minor fraction in BM aspirates. In addition, few human MC lines are available and among those, only two, HMC-1.2 and ROSA^D816V^, express *KIT* D816V.^8^ Additionally, engineered or established *KIT* D816V cell lines do not fully recapitulate *KIT* D816V SM as additional mutations in genes such as *ASXL1*, *CBL*, *RUNX1*, *SRSF2* and *TET2*, are commonly found in patients with *KIT* D816V ASM and MCL.^8,22–24^ These concurring mutations contribute to disease heterogeneity and progression. Therefore, to study SM pathology and to screen for compounds targeting SM cells, more authentic disease models are required that more faithfully recapitulate the genetic and functional features of SM.

Patient-derived iPSCs provide an inexhaustible cell source for disease modeling and drug screening while retaining the patient-specific genetic background, including disease-specific and/or associated mutations.^25–27^ Additionally, iPSCs and embryonic stem cells (ESCs) are readily subjected to genome engineering by CRISPR/Cas9 to precisely introduce or repair oncogenic mutations.^26,28,29^

We report here on the generation of *KIT* D816V iPSCs from SM patients. SM-derived *KIT* D816V iPSCs differentiated into hematopoietic progenitor cells (HPCs) and MCs with patient-specific features. Additionally, we introduced the *KIT* D816V mutation into human ESCs by CRISPR/Cas9n thus generating a panel of *KIT* D816V and *KIT*-unmutated iPSC and ESC lines. Compound screening identified the TKI nintedanib (Vargatef, Ofev) and its analogues as potent novel KIT D816V inhibitors. This finding was recapitulated in iPSC and ESC-derived *KIT* D816V hematopoietic cells and SM primary samples, thereby demonstrating KIT D816V specific targeting by nintedanib.

## Materials and methods

### iPSC generation and culture

Reprogramming of SM primary samples (supplemental Table 1) and cultivation of iPSCs was performed as described previously.^29^ *KIT* D816V mutation was detected by allele-specific PCR (supplemental Table 2).

### Generation of *KIT* D816V ESCs by CRISPR/Cas9n editing

Human HES-3 ESCs (ES03) were from WiCell Research Institute and cultured as described for iPSCs.^29^ ESC work was approved by German authorities, Robert Koch Institute, Berlin, Germany (permit no. 1710-79-1-4-79). *KIT* D816V ESCs were generated as described previously using the pX335 vector (Addgene 42335) and oligonucleotides listed in supplemental Table 3.^29^

### Cytogenetic analysis and Epi-Pluri test

All iPSCs clones generated in this work were subjected to karyotype analysis using GTG banding as before.^29^ Epi-Pluri-Score analysis was performed as described.^30^

### Immunofluorescence staining

iPSCs and ESCs were stained for pluripotency markers and endothelial cells were stained for CD31 and CD144 (supplemental Table 4) essentially as described previously.^29^

### Hematopoietic differentiation of iPSCs and ESCs

iPSCs were differentiated into hematopoietic progenitor cells and MCs using an embryonic body (EB)-based protocol.^31,32^ ESCs were differentiated towards the hematopoietic lineage by adapting a hypoxia-based protocol.^29^ CD34^+^ HPCs were isolated by MACS (Miltenyi Biotech) and expanded for up to 12 days at 37°C and 5% CO_2_ in hematopoietic differentiation medium.^31^

### Compound testing on hematopoietic cells, MCs and endothelial cells

MACS selected KIT^+^ or KIT^−^ hematopoietic cells derived from iPSCs or ESCs were cultured with compounds in 96-well format (Greiner) with 10^4^ cells/well in 90 μl drug screening medium (RPMI 1640 supplemented with 10% FCS, 2 mM L-glutamine, 100 U/ml penicillin and 100 μg/ml streptomycin (all Thermo Fisher Scientific)) for 66 h. Cell viability was determined with CellTiter-Glo Luminescent Cell Viability Assay and SpectraMAX i3 Plate Reader and Softmax Pro Software.

CD45^+^KIT^high^ iPSC-derived MCs were seeded at a density of 5×10^3^ cells/well and subjected to compound testing as above. Endothelial cells (10^4^ cells/well) were treated with compounds in fully supplemented EGM-2 medium (LONZA) for 48 h and then analyzed as above.

Detailed methods for NGS analysis, endothelial cell differentiation, proliferation assay, LDL uptake assay, flow cytometry analysis and sorting, apoptosis assay, Western blotting, CFU assay, cytospin preparations, DSRT assay, compound testing on primary cells, RT-PCR analysis and molecular docking are provided in the supplemental Methods (available on the Blood Web site).

## Results

### *KIT* D816V iPSCs express a constitutively active KIT receptor

*KIT* D816V iPSCs were generated from 14 *KIT* D816V/H SM patients by reprogramming PB or BM mononuclear cells (supplemental Table 1). More than 1000 iPSC lines were obtained and 5 *KIT* D816V iPSCs (3 of patient 1, 1 of patient 2 and 1 of patient 3) and 5 unmutated *KIT* iPSCs (2 of patient 1, 1 of patient 2 and 2 of patient 3, hereafter referred to as controls) were used in the present work. All 10 iPSC lines were pluripotent as determined by morphology, expression of the pluripotency markers OCT4, NANOG, TRA-1-60 and TRA-1-81 and by Epi-Pluri-Score analysis (Figure 1A and supplemental Figure 1, A and B).^30^ Karyotype analysis of all iPSCs showed normal GTG banding and no numeric abnormalities (supplemental Figure 1C).

**Figure 1.**
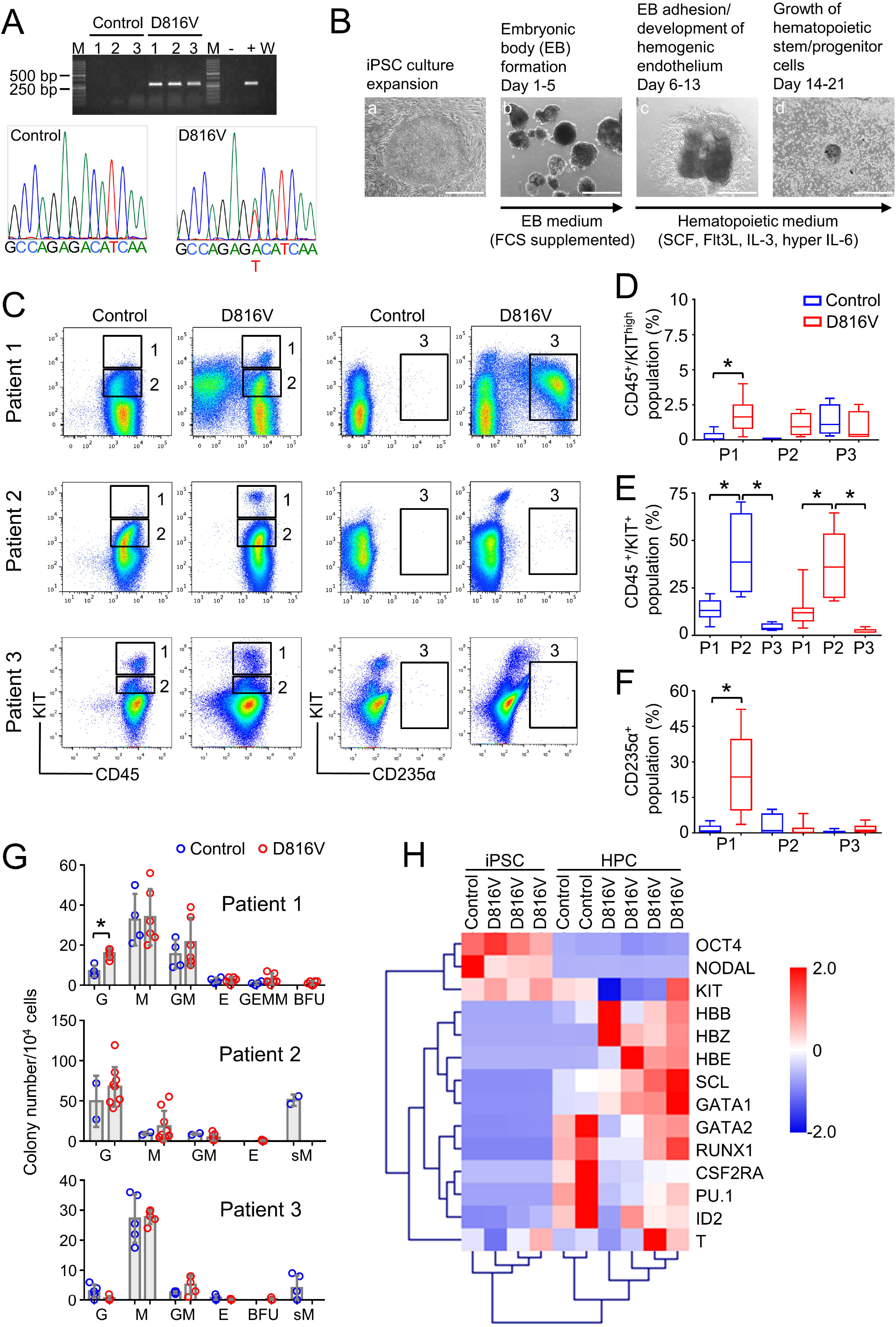
Hematopoietic differentiation of patient-specific *KIT* D816V iPSCs. (A) *KIT* D816V mutation in iPSCs by allele-specific PCR (top). Control 1-3: iPSCs without mutation. D816V 1-3: iPSCs with mutation. (-) HMC-1.1 cell line. (+) HMC-1.2 cell line. W: water control. M: molecular weight marker. *KIT* D816V mutation in iPSCs by Sanger sequencing (bottom). (B) Schematic representation of hematopoietic differentiation protocol. Embryonic bodies (EBs) were formed from iPSCs (a, b) and differentiated towards the hematopoietic lineage (c, d). Scale bar: 500 μm. (C) Representative flow cytometry analysis of *KIT* D816V and control iPSC-derived hematopoietic cells of patients 1-3. The plots represent synchronized hematopoietic differentiation experiments and show patient-specific phenotypes. Gates 1, 2 and 3 are for CD45^+^/KIT^high^, CD45^+^/KIT^+^ and CD235α^+^ cells, respectively. (D) Quantification of CD45^+^/KIT^high^ populations for *KIT* D816V and control iPSC-derived cells of all 3 patients (P1, P2, and P3). Flow cytometry was performed between 15 and 31 days of hematopoietic differentiation (n=4-21). *: p<0.0001. (E) Quantification of CD45^+^/KIT^+^ populations by flow cytometry as in (D) between 15 and 49 days of differentiation (n=4-30). *: p≤0.0003 (F) Quantification of CD235α^+^ population by flow cytometry as in (D) between 15 and 49 days of differentiation (n=4-30). *: p<0.0001. (G) Colony-forming unit (CFU) assay for *KIT* D816V and control iPSC-derived hematopoietic cells. Colony numbers and phenotype were evaluated 14 days after seeding. Bars indicate average colony numbers ±SD of at least 3 independent experiments, except for patient 2 control iPSC (n=2). *: p=0.02. (H) RT-qPCR data of patient 1-derived *KIT* D816V and control iPSCs (iPSC) and corresponding hematopoietic progenitor cells (HPC). Results of 2 and 4 independent hematopoietic differentiation experiments for control and *KIT* D816V HPC, respectively, are shown. Gene expression values were subjected to bidirectional hierarchical clustering and are shown in heatmap format (red and blue, high and low gene expression, respectively). Statistical analysis was performed with Welch’s t-test.

All *KIT* D816V iPSCs showed reduced KIT surface expression in comparison to unmutated *KIT* cells (supplemental Figure 1, D and E), which is in line with KIT D816V being confined to and signaling from intracellular compartments.^9^ Additionally, *KIT* D816V iPSCs showed strong phosphorylation of glycosylated and non-glycosylated forms of the receptor without SCF stimulation (supplemental Figure 1F). Control iPSCs showed very low phosphorylation of glycosylated KIT and upon SCF stimulation, receptor internalization and phosphorylation of unmutated glycosylated KIT, AKT and STAT3 were observed. These data are in accordance with KIT D816V representing a constitutively active receptor in iPSCs.

### *KIT* D816V and control iPSCs exhibit patient-specific phenotypes upon hematopoietic differentiation

*KIT* D816V and control iPSCs were induced to differentiate into hematopoietic cells in an EB-based protocol (Figure 1B).^31^ For both, *KIT* D816V and control iPSC-derived cells, a CD45^+^/KIT^high^ MC population was observed. In patient 1 and 2 samples, this population was more prominent in the *KIT* D816V genotype (Figure 1, C and D) and was observed at early time-points (>14d, supplemental Figure 2A). *KIT* D816V conferred a proliferative advantage to KIT^+^ cells, regardless of SCF stimulation compared to *KIT*-unmutated KIT^+^ cells (supplemental Figure 2B). *KIT* D816V also sustained cell viability upon cytokine withdrawal. Thus, *KIT* D816V promotes MC development *in vitro* in our iPSC model and confers higher proliferation capacity to hematopoietic cells.

Additionally, iPSCs exhibited patient-specific differentiation propensities. Patient 1 *KIT* D816V iPSC-derived cells showed a prominent erythroid CD235α^+^CD43^+^KIT^+^ population and, in colony forming unit (CFU) assays, BFU-E colonies were observed only in the mutated genotype (Figure 1, C, F and G, supplemental Figure 3, A and B). Cytospin preparations of *KIT* D816V CD235α^+^ cells revealed different stages of erythroid maturation (supplemental Figure 3C). Patient 2 iPSCs showed a prominent CD45^+^/KIT^+^ population during differentiation with myeloid progenitor morphology concomitantly with prominent apoptosis (Figure 1, C and E; supplemental Figure 4A and B). CFU assays showed a strong bias towards granulocytes and macrophages for both, *KIT* D816V and control cells, while abnormal CFU-M of small size (CFU-sM) were observed only in controls (Figure 1G). Patient 3 *KIT* D816V and control iPSC-derived hematopoietic cells showed a prominent CD45^+^/KIT^high^ MC population (Figure 1, C and D and supplemental Figure 5A). CFU assays revealed reduced colony forming potential and a bias towards the macrophage lineage, in agreement with the abundance of macrophages in cytospin preparations (Figure 1G and supplemental Figure 5B).

Finally, iPSC-derived cells exhibited hematopoietic gene expression profiles, and also recapitulated the erythroid bias observed in patient 1 *KIT* D816V iPSC-derived hematopoietic cells, shown by the high expression of hemoglobin genes *HBB*, *HBE* and *HBZ* (Figure 1H).

### *KIT* D816V iPSCs harbor patient specific mutation profiles

Previous studies have shown additional mutations further to *KIT* D816V in patients with advanced mastocytosis, most frequently in *ASXL1*, *CBL*, *RUNX1*, *SRSF2*, and *TET2*, which correlate with poor survival.^22,23^ In agreement with these observations, we identified patient-specific subset of mutations in ASM and MCL primary samples, which were recapitulated in the iPSCs derived thereof (supplemental Figure 6 and supplemental Table 5-7). Of particular relevance, concurrent to *KIT* D816V in patient 1 iPSCs, a *NFE2* truncating mutation was identified. In patient 2, a *SRSF2* in-frame 8 amino-acid deletion (P95_R102del) was detected in all iPSC cell lines while additional likely pathogenic mutations in *RUNX1* and *TET2* were found only in control iPSC. In patient 3 primary sample, no *KIT* D816V or other pathogenic or likely pathogenic mutations were detected, in agreement with the low number of *KIT* D816V iPSCs obtained (1 out of 129 screened).

### *KIT* D816V ESCs recapitulate phenotypes of *KIT* D816V iPSCs

To address the question of a potential influence of concurrent mutations on SM iPSC differentiation, we introduced the *KIT* D816V mutation into human ESCs by CRISPR/Cas9n technology (supplemental Figure 7). As observed for *KIT* D816V iPSCs, KIT receptor surface expression in *KIT* D816V ESCs was lower than in control ESCs, although no differences in *KIT* mRNA expression were detected (supplemental Figure 8A). Strong phosphorylation of non-glycosylated KIT D816V receptor was also observed in Western blot analysis (supplemental Figure 8, B and C).

*KIT* D816V ESCs differentiation yielded >95% CD45^+^ hematopoietic cells, including CD45^+^/KIT^high^ MCs (Figure 2A and supplemental Figure 9). Importantly, *KIT* D816V ESC-derived hematopoietic cells showed an erythroid bias, although not as prominent as *KIT* D816V iPSCs-derived cells of patient 1 (CD235^+^ ESC D816V 1 = 8.0±2.9, CD235^+^ ESC D816V 2 = 9.6±3.9, CD235^+^ Patient 1 D816V = 24.6±14.5). Accordingly, cytospin preparations showed higher numbers of nucleated erythrocytes for *KIT* D816V cells (Figure 2B). In CFU assays, CFU-E and BFU-E were only observed for mutated progenitors as well as higher expression of hemoglobin genes *HBB*, *HBE* and *HBZ* (Figure 2 A, C and D). No significant differences were observed for *KIT* D816V and control CD45^+^/KIT^high^ MC populations, in agreement with the reported weak oncogenic effect of *KIT* D816V alone.^33^ Taken together, these results demonstrate that *KIT* D816V ESCs recapitulate the phenotypes observed in patient-derived *KIT* D816V iPSCs, such as constitutive KIT phosphorylation, reduced surface expression of the mutated receptor and erythroid bias upon hematopoietic differentiation.

**Figure 2.**
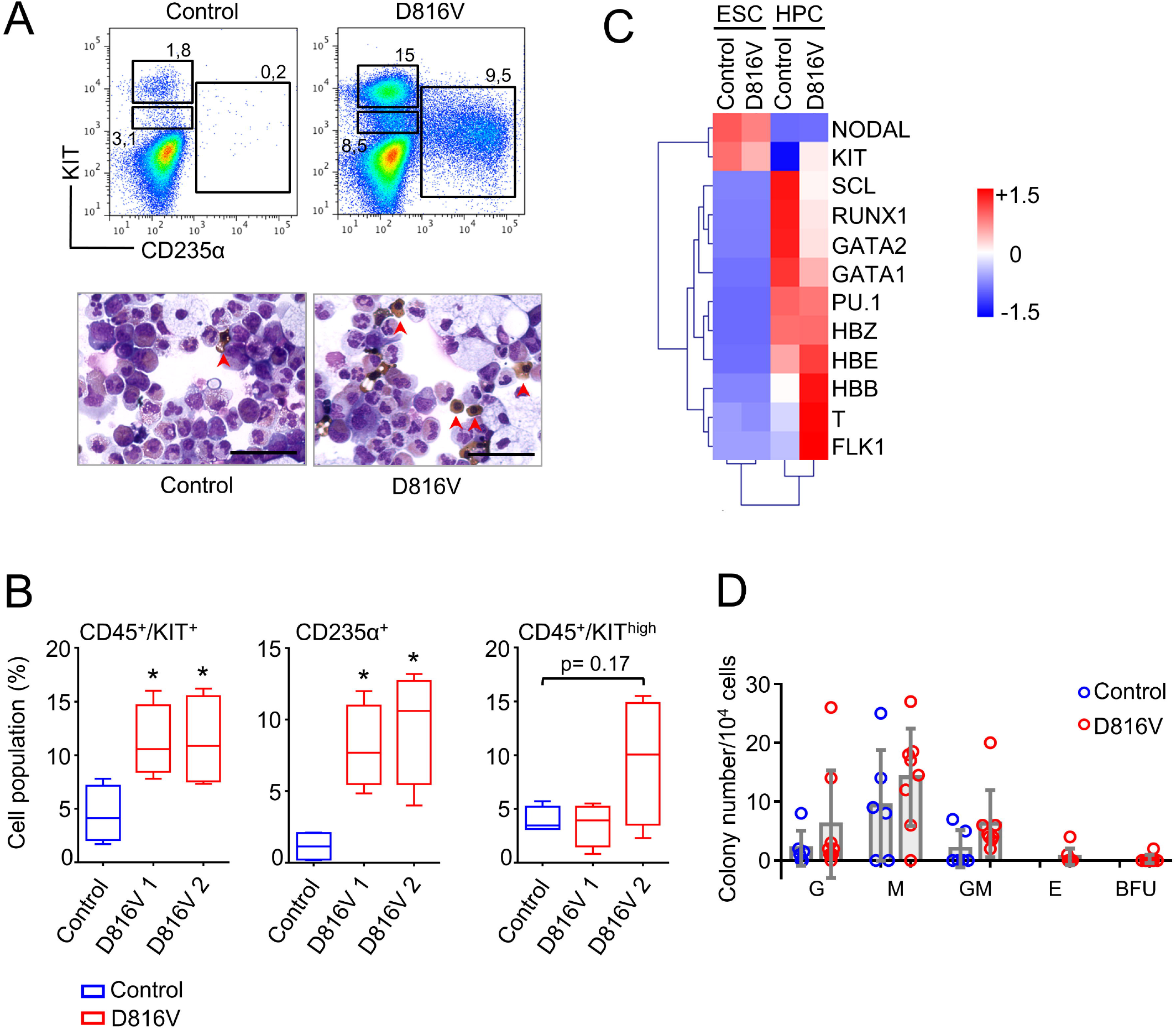
*KIT* D816V ESCs recapitulate the KIT^high^ and erythroid phenotype of patient *KIT* D816V iPSCs. (A) Flow cytometry analysis of *KIT* D816V and control ESC-derived hematopoietic cells 11 days after CD34^+^ MACS selection (top). Prominent CD235α^+^ and CD45^+^/KIT^high^ populations are observed in *KIT* D816V ESC-derived cells. Cytospin preparations of the same samples show nucleated erythrocytes (red arrowhead, bottom). Scale bar: 50 μm. (B) Quantification of hematopoietic cell populations (n=4) derived from *KIT* D816V clone 1 and D816V clone 2 ESCs and control ESCs. *KIT* D816V ESCs show prominent CD45^+^/KIT^+^ and CD235^+^ populations. CD45^+^/KIT^high^ MC population showed no statistically significant differences between *KIT* D816V and control. *: p<0.05. (C) RT-qPCR data of *KIT* D816V and control ESCs (ESC) and corresponding hematopoietic progenitor cells (HPC) 11 days after CD34^+^ enrichment by MACS. Gene expression values were subjected to bidirectional clustering and are shown in heatmap format (red and blue, high and low gene expression, respectively). (D) Colony forming unit (CFU) assay for hematopoietic cells obtained from *KIT* D816V and control ESCs. Bars show the mean ± SD of 3 independent experiments. Statistical analysis was performed with Welch’s t-test.

### The tyrosine kinase inhibitor nintedanib targets *KIT* D816V hematopoietic cells

We next focused on identifying compounds that selectively target *KIT* D816V iPSC-derived hematopoietic cells. A library of 459 compounds was first tested on HMC-1 cell lines (supplemental Figure 10 and supplemental Table 8) and selected compounds were further analyzed on iPSC-derived *KIT* D816V hematopoietic cells (Figure 3, A and B). We identified nintedanib as a potent KIT D816V inhibitor, significantly reducing cell viability of *KIT* D816V hematopoietic cells of all 3 patients in the nanomolar range (IC50_D816V_=27.5-104.9 nM, IC50_control_=262.1-541.7 nM). We also included midostaurin and imatinib in our studies as midostaurin is used to treat patients with advanced SM while imatinib is ineffective in *KIT* D816V SM.^15–17,20^ Nintedanib and midostaurin showed similar potency on *KIT* D816V cells and, as expected, imatinib was essentially ineffective. Rigosertib, a RAS mimetic compound, was also tested as oncogenic RAS mutations are commonly reported in hematologic malignancies, including SM.^2,34^ Therefore, simultaneous targeting of KIT and RAS might have positive therapeutic outcome. In this context, we observed that *KIT* unmutated cells derived from patient 1 expressing the *NRAS* G12D mutation (supplemental Figure 6) were highly responsive to rigosertib (Figure 3B).

**Figure 3.**
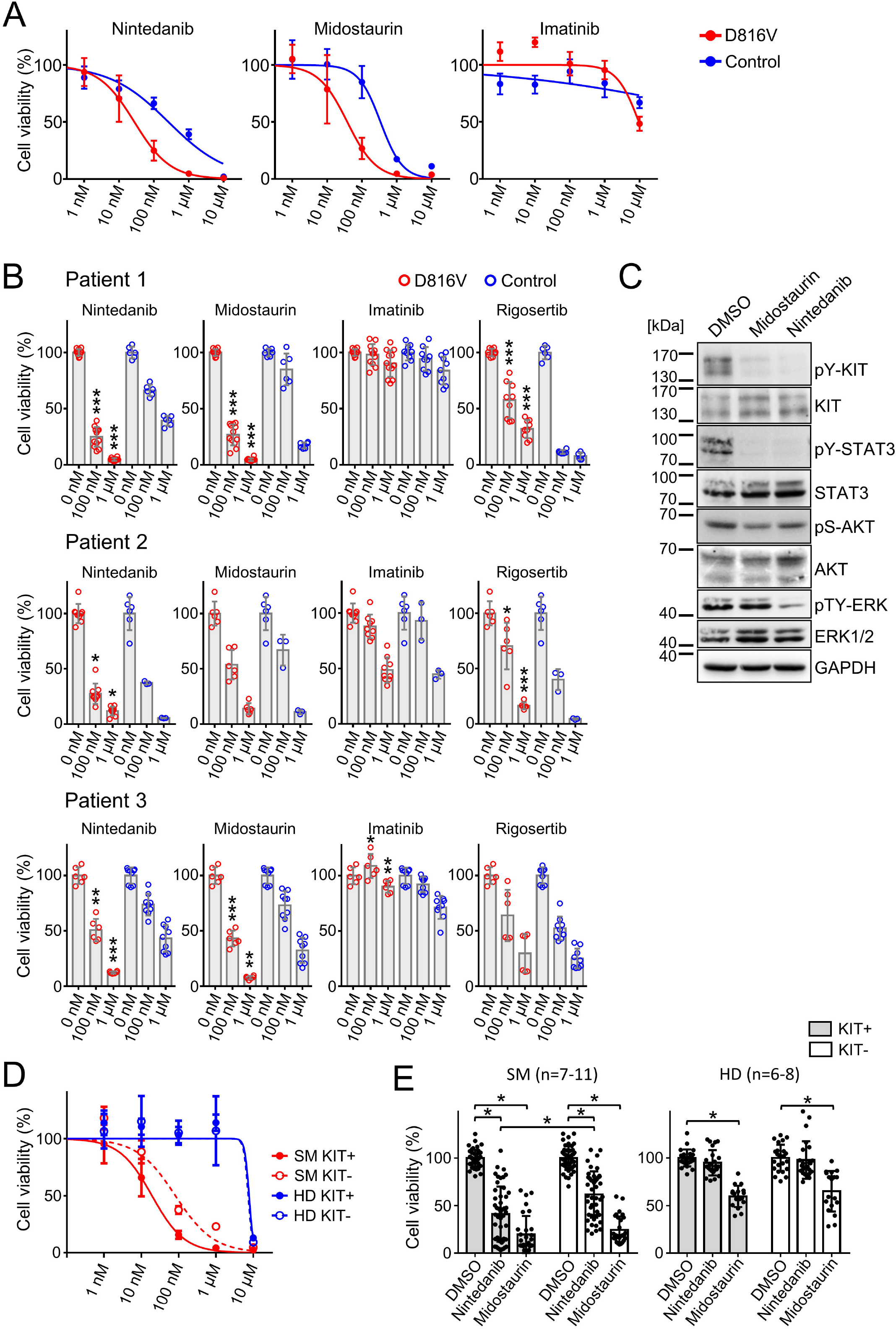
Compound testing on *KIT* D816V iPSC/ESC-derived hematopoietic cells and evaluation of nintedanib effect on SM patient samples. (A) Drug response curves (0 to 10 μM) for nintedanib, midostaurin and imatinib on *KIT* D816V (n=1 to 4) and control (n=2) iPSC-derived KIT^+^ hematopoietic cells of patient 1. IC50 values were calculated based on averaged titration curves obtained for cells derived from different iPSC lines. IC50 values for nintedanib were 27-105 nM for *KIT* D816V cells and 262-542 nM for control cells. (B) Averaged drug response ± SD of *KIT* D816V and control iPSC-derived KIT^+^ cells treated with 100 nM or 1 μM of nintedanib, midostaurin, imatinib or rigosertib for 66 h. Vehicle (DMSO) treated cells were used as control (0 nM). Statistical analysis was performed by Welch’s t-test comparing the drug responses of *KIT* D816V and control KIT^+^ cells at the same drug concentration. Patient 1: n=6-12. Patient 2: n=3-9. Patient 3: n=6-9. *: p≤0.05; **: p<0.001, ***: p≤0.0001. (C) Representative Western blot analysis of KIT receptor signaling upon nintedanib or midostaurin treatment of *KIT* D816V iPSC-derived hematopoietic cells. Cells were treated with 1 μM of compound for 4 h prior to analysis. Vehicle (DMSO) treated cells were used as control. Positions of molecular weight marker are indicated. (D) Nintedanib response curves (0 to 10 μM) for SM primary sample (patient 10, red) and healthy donor (n=2, blue). MNCs were subjected to MACS and KIT^+^ and KIT^−^ cells were treated with nintedanib for 66 h followed by viability measurement using CellTiter Glo assay. Vehicle (DMSO) treated cells were used as control. Nintedanib shows cytotoxicity to healthy donor cells only at concentrations closer to 10 μM while SM cell viability is severely compromised at concentrations higher than 100 nM. (E) Averaged response of 7-11 SM primary samples and 6-8 healthy donor (HD) primary samples to 1 μM nintedanib or midostaurin treatment. MNCs were treated as described in (D). Nintedanib treatment led to a significant decrease in the viability of KIT^+^ SM MNCs while midostaurin equally targeted KIT^+^ and KIT^−^ cells. Additionally, nintedanib had no significant impact on the viability of HD cells, in contrast to midostaurin that led to significant reduction in the viability of KIT^+^ and KIT^−^ HD cells. *: Welch’s t-test, p≤0.0003.

We further compared drug responses of KIT^+^ and KIT^−^ cells and observed selective targeting of KIT^+^ cells by nintedanib and midostaurin, which was more pronounced in *KIT* D816V cells than in control cells (supplemental Figure 11A). Western blot analysis of iPSC-derived cells treated with nintedanib or midostaurin revealed an efficient reduction in KIT and STAT3 phosphorylation and those effects were more pronounced in nintedanib treated samples, where a significant reduction in total KIT protein was observed (Figure 3C and supplemental Figure 12).

Importantly, nintedanib and midostaurin activities were recapitulated in *KIT* D816V ESC-derived cells (supplemental Figure 11, B and C). The higher selectivity of nintedanib and midostaurin for KIT^+^ over KIT^−^ cells was also observed in ESC-derived cultures (supplemental Figure 11D). Thus, data obtained with our *KIT* D816V ESCs fully recapitulate hematopoietic phenotypes and drug responses observed for *KIT* D816V iPSC-derived cells.

In order to evaluate whether the origin and differentiation stage of iPSC-derived cells impact drug responses, we MACS selected CD34^+^ HPCs from the hemogenic endothelium and expanded cells for further 10-20 days. Again, a strong reduction in viability of KIT^+^ mutated cells was observed upon treatment with nintedanib or midostaurin (supplemental Figure 13 A-C). Additionally, CD34^+^ HPCs displayed the same cell fate bias as cells obtained directly from iPSCs, such as the erythroid bias observed in patient 1 (supplemental Figure 13D).

We further compared the response of *KIT* D186V iPSC-derived hematopoietic cells to nintedanib, avapritinib (BLU-2815) or ripretinib (DCC-2618), as the last two compounds are currently being evaluated in clinical trials for advanced SM.^19–21,35^ Avapritinib and ripretinib effectively reduced viability of HMC-1.1 and HMC-1.2 cell lines and phosphorylation of KIT, STAT5, AKT and ERK (supplemental Figure 14), which is in accordance with previous studies.^19,21,35^ Finally, as observed for nintedanib, avapritinib and ripretinib preferentially reduced the viability of iPSC-derived *KIT* D816V cells (supplemental Figure 15). These results show selectivity of avapritinib and ripretinib for KIT D816V and validate the robustness of the *KIT* D816V iPSC system established in this study.

Next, we accessed the impact of nintedanib treatment on viability of KIT D816V/H and KIT S476I SM primary samples. A reduction ≥50% in cell viability was observed for 6 out of 9 primary MNCs samples (supplemental Figure 16A and supplemental Table 9). Additionally, nintedanib preferentially targeted KIT+ over KIT− primary MNCs leading to a reduction in cell viability ≥50% in 6 out of 11 primary samples. In contrast, midostaurin showed less selectivity and affected both KIT+ and KIT− primary cells from SM patients and healthy donors (Figure 3D and E, supplemental Figure 16B and supplemental Table 9). Nintedanib treatment also led to a reduction in KIT D816V allele burden in SM MNCs (supplemental Figure 16C). Western blot analysis showed strong reduction of KIT, STAT5 and ERK phosphorylation in KIT D816H SM primary sample upon 1μM nintedanib treatment (supplemental Figure 16D) in agreement with the data obtained with our iPSC-based model.

### *KIT* D816V iPSC-derived MCs are targeted by nintedanib

A CD45^+^/KIT^high^ MC population was obtained at later time points of hematopoietic differentiation for all *KIT* D816V and control iPSCs lines used in the present study (Figure 4A). *KIT* D816V MCs showed higher surface and gene expression of FCER1A than control cells, while no differences were observed for CD45 and KIT (Figure 4, B and E). FACS purified MCs revealed a homogenous cell population with multilobulated nuclei and some larger cells presenting metachromatic and tryptase positive cytoplasmic granules, features of immature MCs (Figure 4, C and D and supplemental Figure 17). qRT-PCR analysis indeed showed high expression of tryptase gene products TPSAB1 and TPSB2, while CPA3 (carboxypeptidase A3) and CMA1 (chymase 1) mRNA expression was rather low (Figure 4E).

**Figure 4.**
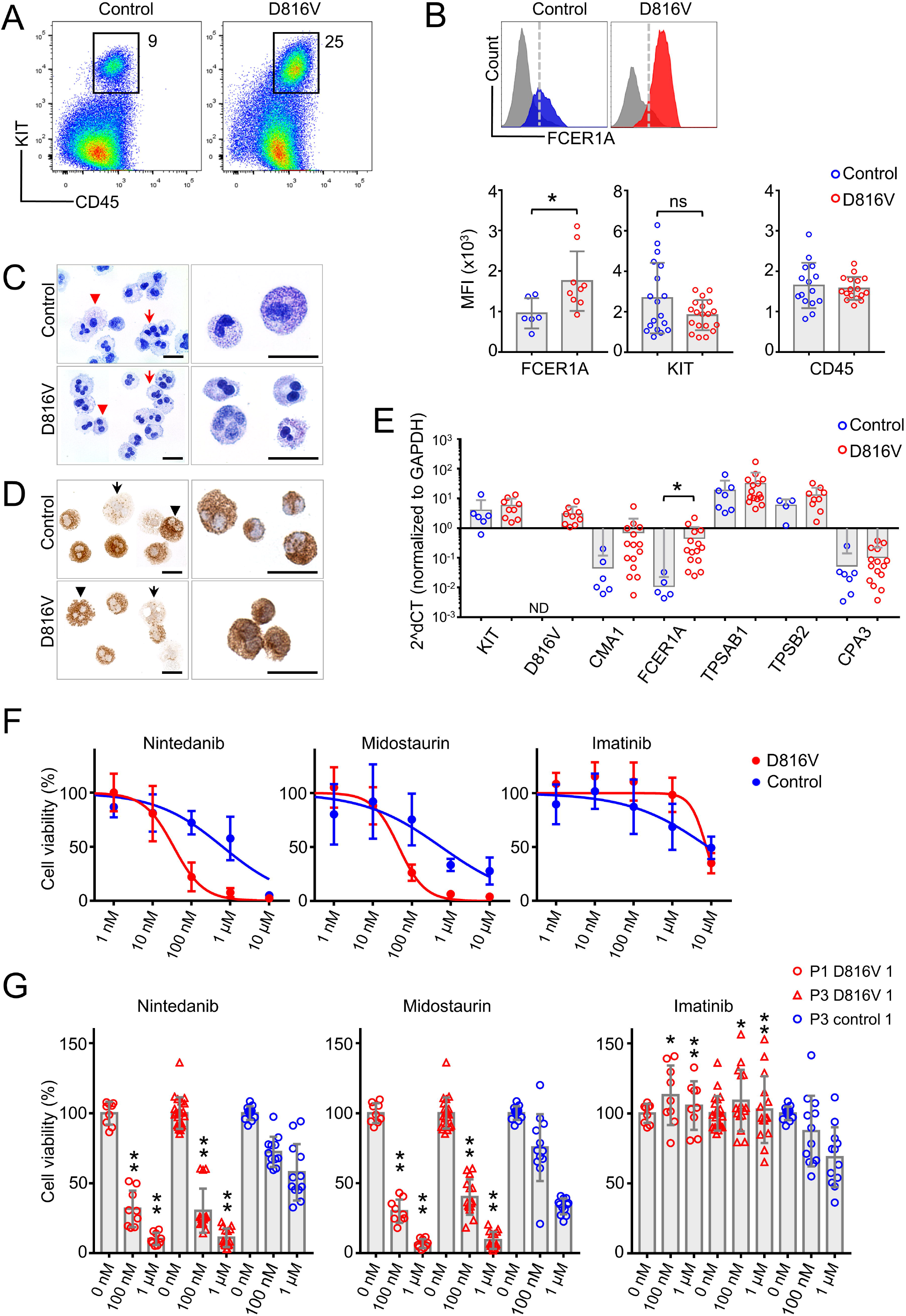
Nintedanib targets *KIT* D816V MCs. (A) Flow cytometry analysis of *KIT* D816V and control iPSC-derived CD45^+^/KIT^high^ MCs at >30 days of differentiation (patient 1). (B) *KIT* D816V MCs (red) show higher expression of FCER1A than control cells (blue, top). Mean fluorescence intensity (MFI) of FCER1A (n=6-9), KIT (n=18-19) and CD45 (n=15-17) on *KIT* D816V MCs and controls (bottom). *: Welch’s t-test, p=0.017. ns: not significant. (C) Representative composite images of acidic toluidine blue stained cytospins (left panels) or smears (right panels) of FACS sorted *KIT* D816V and control MCs. Homogenous population of multilobulated pro-mastocytes with (red arrowheads) or without (red arrows) metachromatic granules were observed. Scale bar: 25 μm. (D) Same as (C) but stained for tryptase. Cells with high (black arrowhead) and low (black arrow) number of tryptase-positive granules were observed. Scale bar: 25 μm. (E) RT-qPCR analysis for *KIT* D816V and control FACS sorted MCs (n=9-15 and n=3-7, respectively). mRNA expression of MC specific chymase 1 (*CMA1*), tryptase alpha/beta 1 (*TPSAB1*), tryptase beta 2 (*TPSB2*), carboxypeptidase A3 (*CPA3*), *FCER1A* and *KIT* are depicted. *KIT* D816V MCs show higher *FCER1A* mRNA expression than unmutated *KIT* cells. *: Welch’s t-test, p=0.014. (F) Drug response curves of FACS sorted *KIT* D816V and control MCs (n=5 and n=4, respectively) treated with nintedanib, midostaurin or imatinib. Nintedanib IC50 values were 34 and 633 nM for *KIT* D816V and control cells, respectively. Midostaurin IC50 values were 48 and 570 nM for *KIT* D816V and control cells, respectively. (G) Averaged drug response ± SD of *KIT* D816V and control iPSC derived FACS sorted MCs (n=9-15 and n=12, respectively) treated with 100 nM or 1 μM of nintedanib, midostaurin or imatinib. Statistical analysis was performed by Welch’s t-test comparing the drug responses of *KIT* D816V and control MCs at the same drug concentration. *: p=0.02. **: p≤0.0006.

Importantly, as observed for hematopoietic progenitor cells, *KIT* D816V MCs displayed remarkable reduction in cell viability upon nintedanib treatment in comparison to control cells (patient 1 IC50_D816V_= 34.7nM; patient 3 IC50_D816V_= 34.6 nM; patient 1 IC50_control_= 633.2 nM, Figure 4, F and G).

### Nintedanib preferentially targets iPSC-derived hematopoietic cells over iPSC-derived endothelial cells

Nintedanib was initially developed as an inhibitor of VEGF receptors, which are crucial for the development and function of endothelial cells.^36,37^ Increased angiogenesis in BM MC infiltrates is a key histopathological feature in SM patients.^38^ Therefore, we determined the impact of nintedanib on endothelial cells derived from SM KIT D816V and control iPSCs (Figure 5A). KIT D816V and control iPSC-derived endothelial cells expressed CD31, CD34, CD105, CD144 and KIT and were capable of lipid (acetylated LDL) uptake (Figure 5, B-E). Nintedanib reduced cell viability of both *KIT* D816V and control endothelial cells and significant differences in drug response were observed with 1 μM of the compound (Figure 5F). Importantly, cytotoxic effects were 20 to 100-fold more pronounced on iPSC-derived hematopoietic progenitor cells and MCs than on endothelial cells, regardless of the presence of *KIT* D816V mutation (Figure 5G).

**Figure 5.**
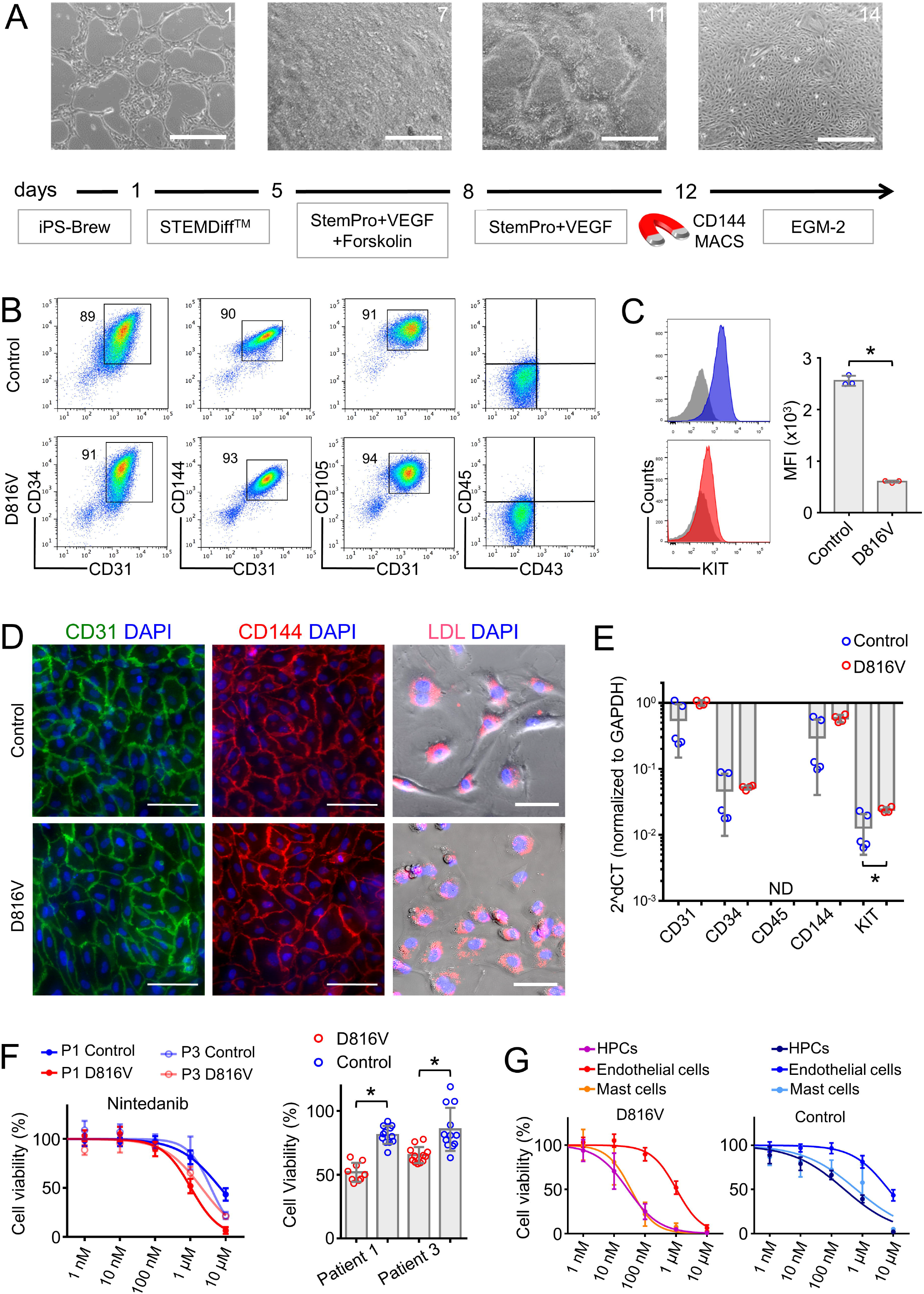
Evaluation of nintedanib activity on iPSC-derived endothelial cells. (A) Schematic representation of the endothelial differentiation for *KIT* D816V or control iPSCs. Representative phase contrast microscopy images of day 1, 7, 11 and 14 are shown. Scale bar: 500 μm. (B) *KIT* D816V and control iPSC-derived endothelial cells express CD34, CD31, CD105 and CD144 endothelial markers but lack CD45 and CD43 expression. (C) *KIT* D816V endothelial cells (red) have lower surface KIT receptor expression than control (blue) as shown by representative histogram plots; isotype control in grey. *: n= 3, p=0.0005. (D) Immunofluorescence staining of *KIT* D816V and control endothelial cells showed homogenous surface expression of CD31 (left panels) and CD144 (middle panels). Cells effectively absorbed Dil-conjugated acetylated-LDL (right panels). Nuclei were stained with DAPI (blue). Scale bar: 100 μm. (E) qRT-PCR data show similar expression of CD31, CD34 and CD144 in *KIT* D816V and control endothelial cells. *KIT* mRNA expression was slightly higher in mutated cells. *: n_=_4-5, p=0.03. ND= not detected. (F) *KIT* D816V and control endothelial cells derived from patients 1 and 3 iPSCs showed similar nintedanib response curves (left, n=2-3, patient 1 (P1), IC50_D816V_=1632 nM, IC50_control_=7003 nM; patient 3 (P3), IC50_D816V_=2073 nM, IC50_control_=3803 nM). *KIT* D816V endothelial cells were more affected by 1 μM nintedanib treatment than unmutated cells (right). *: Welch’s t-test, n=3-4, p<0.01. (G) Comparison of drug response curves obtained for *KIT* D816V (left) or control (right) endothelial cells (n=2-3), hematopoietic progenitor cells (HPC, from Figure 3) and MCs (from Figure 6) treated with nintedanib (1 nM to 10 μM).

### Nintedanib occupies the ATP-binding site in KIT D816V

Nintedanib is an indolinone derivative type II kinase inhibitor of VEGFR, PDFGR and FGFR.^36,37^ It binds to the ATP-binding site cleft between C- and N-terminal lobes of VEGFR kinase domain.^36^ Thus, we compared the nintedanib binding site in VEGFR2 (PDB code 3C7Q) with the ATP binding sites of all available structures of KIT (unmutated and KIT D816V, supplemental Figure 18A). Our analysis revealed geometrical similarity of VEGFR2 and KIT and steric fitting of nintedanib into the inactive KIT receptor, in agreement with the biological data reported in the present work. Induced fit molecular docking studies showed that, on average, nintedanib has a higher affinity (evaluated here as Glide Score) for inactive KIT D816V (based on PDB code 3G0F) than for unmutated KIT, either in its inactive or active state (PDB code 3G0E and 1PKG, respectively, supplemental Figure 18B). Nintedanib preferentially binds to KIT D816V by orienting the indole moiety towards the A-loop and the methylpiperazin moiety towards the C-terminal region (Figure 6).

**Figure 6.**
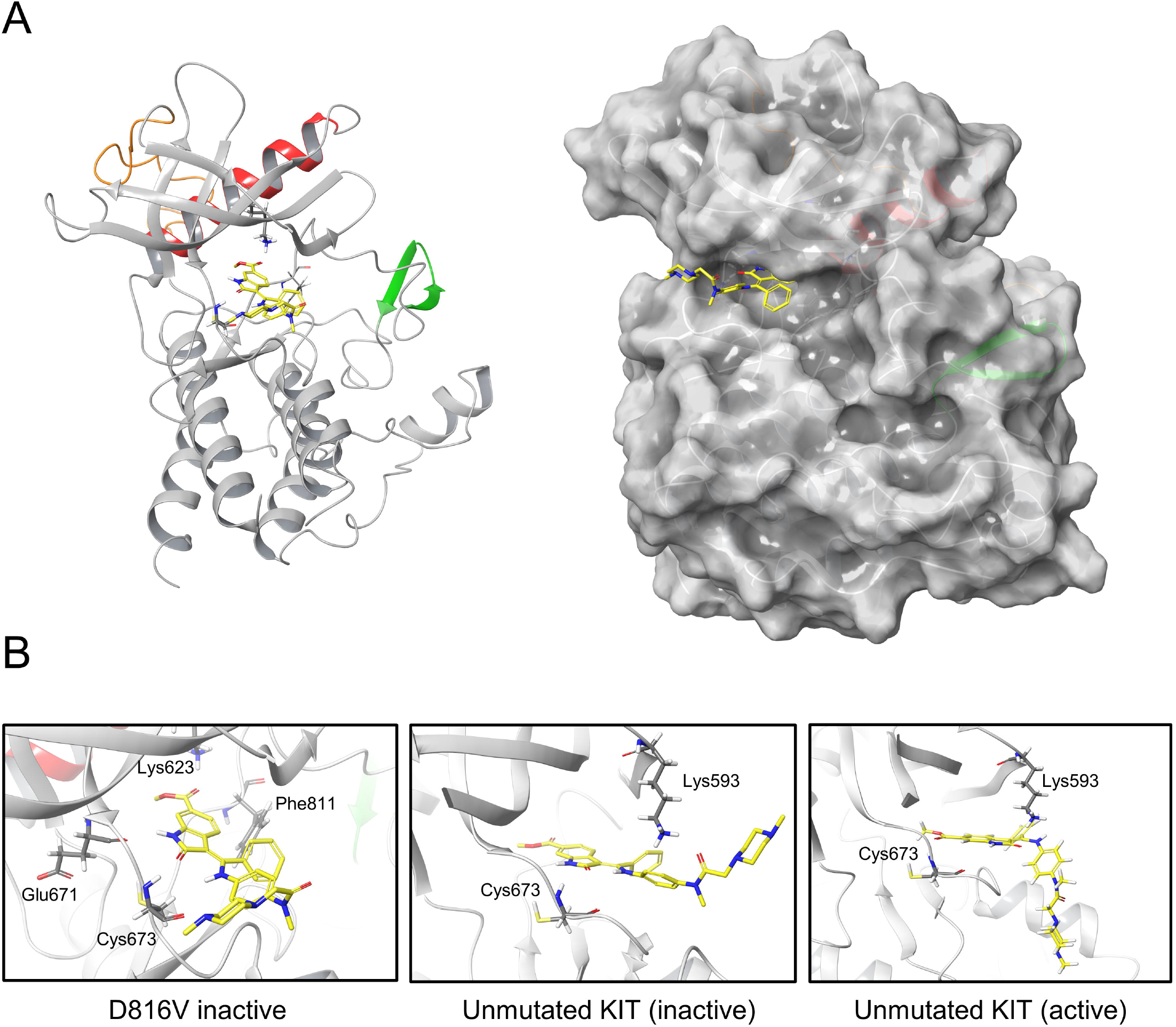
Molecular docking of nintedanib in KIT D816V. (A) KIT D816V ribbon structure with nintedanib in cartoon (left) and surface representation (right). The JMR, A-loop and C-helix are indicated in orange, green and red, respectively. Nintedanib is represented in yellow licorice. (B) Protein-ligand interaction details for nintedanib docked in the inactive KIT D816V (left), inactive and active unmutated KIT (middle and right, respectively) are shown in 3D ribbon structure. Amino acids that are key to ligand-receptor interaction are highlighted. The positioning of nintedanib indole moiety is similar to that observed for sunitinib when bound to KIT (PDB code 3G0E or 3G0F) and it establishes the same hydrogen bonds with Glu671 and Cys673 plus an additional pi-stacking with Phe811. Other hydrophobic contacts are reported in Supplemental Figure 19.

Further analysis revealed that midostaurin and avapritinib display higher affinity for the active unmutated KIT receptor and poor affinity for inactive KIT (unmutated or KIT D816V), while ripretinib and imatinib showed preference for the inactive KIT (unmutated or KIT D816V) over the active conformation (supplemental Figures 18B and 19), in agreement with previous reports.^19,39^

### Nintedanib analogues are potent KIT D816V inhibitors

To further extend the structure-function relation of nintedanib and KIT D816V, we screened a library of 43 nintedanib analogues with 80-98% structural similarity to nintedanib (supplemental Table 10). Compounds were tested on *KIT* D816V and control iPSC-derived hematopoietic cells and 7 of them preferentially targeted *KIT* D816V cells (Figure 7A). Four compounds (BIBG1724TF, BIBF1496XX, BIBE3315BS, and BIBE3316BS) were further evaluated and showed IC50 values similar to nintedanib (Figure 7 B-C).

**Figure 7.**
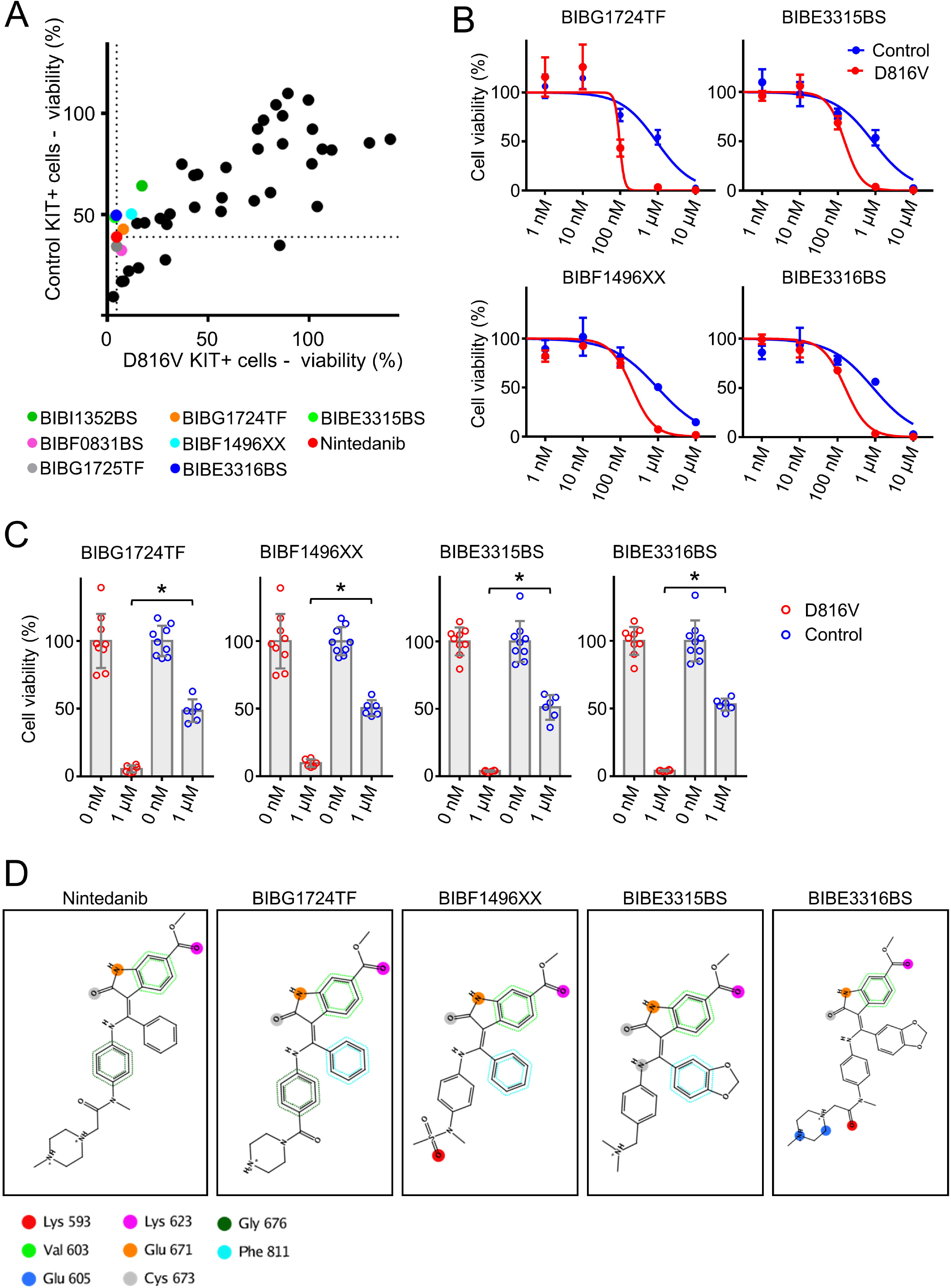
Nintedanib analogues target *KIT* D816V hematopoietic cells. (A) The impact of 43 nintedanib analogues on *KIT* D816V and control iPSC-derived hematopoietic cells was determined in CellTiter Glo assays (1 μM of compound) and is shown as percent of untreated cells. KIT D816V selective compounds are highlighted and color coded. Dotted lines indicate response to nintedanib treatment. (B) Representative drug response curves for *KIT* D816V and control iPSC-derived hematopoietic cells treated with BIBG1724TF, BIBE3315BS, BIBF1496XX or BIBE3316BS. IC50 values were 96-203 nM and 849-917 nM for *KIT* D816V and control cells, respectively. (C) Averaged drug response ± SD of *KIT* D816V iPSC-derived hematopoietic cells treated with 1 μM of BIBG1724TF, BIBF1496XX, BIBE3315BS or BIBE3316BS show significant decrease in cell viability in comparison to treated control cells. *: Welch’s t-test, n=6 to 9, p<0.0001. (D) Structures and interactions based on protein-ligand interaction fingerprint (PLIF) as described in Supplemental Figure 20B. Lys593, Glu605, Lys623, Glu671, and Cys673 establish H-bond interactions with the ligands. Val603/Gly676 and Phe811 establish H-arene interactions and aromatic interactions, respectively, with the aromatic moieties of the ligands. These interactions are represented as dotted points since they are spread over all the atoms of the aromatic rings. The colors uniquely identify each residue, but do not refer to the kind of interaction established.

To rationalize at the molecular level the observed activity of nintedanib derivatives, we modeled these compounds against KIT D816V and unmutated KIT structures by performing induced fit molecular docking calculations. All nintedanib derivatives displayed highest Glide Scores for KIT D816V in comparison to unmutated KIT, suggesting preferential binding to the mutated receptor (supplemental Figure 20A), which agrees with our assays on *KIT* D816V cells. When bound to KIT D816V, BIBG1724TF, BIBF1496XX, BIBE3315BS, and BIBE3316BS located their indole-like moiety toward the A-loop in a similar position as the indole moiety of nintedanib and they established similar molecular contacts with the same amino acid residues in KIT D186V (Figure 7D and supplemental Figure 20B).

## Discussion

We report here on *KIT* D816V iPSCs of ASM and MCL patients that upon differentiation into hematopoietic progenitor cells and MCs recapitulate the SM phenotype. In this model, drug screening identified nintedanib (Vargatef, Ofev) as potent TKI targeting KIT D816V neoplastic cells from patient-specific iPSCs, gene-edited ESCs and primary SM samples.

KIT D816V iPSCs harbor concurring mutations in genes such as TET2, NEF2, NRAS, SRSF2, RUNX1, thereby reflecting the clonal composition of patient samples and SM heterogeneity. The impact of these mutations was observed during *in vitro* differentiation as iPSCs harboring *KIT* D816V and *NFE2* mutations presented an erythroid biased hematopoiesis, while mutations in *TET2*, *RUNX1* and *SRSF2* led to myeloid biased hematopoiesis and pronounced apoptosis, in agreement with published data.^40–44^

Nintedanib is a type II kinase inhibitor currently being used for treatment of non-small cell lung cancer and idiopathic pulmonary fibrosis.^36,37,45^ Importantly, nintedanib has a manageable safety and tolerability profile in long-term use in pulmonary fibrosis patients.^46^ Nintedanib treatment decreased the viability of KIT-expressing iPSC-derived hematopoietic cells and strongly reduced KIT and STAT3 phosphorylation, in agreement with the important role of STAT proteins in downstream signaling of oncogenic KIT.^47^ *KIT* D816V iPSC-derived cells were also targeted by midostaurin, avapritinib (BLU-285) and ripretinib (DCC-2618), in agreement with previous reports and clinical trials for these compounds.^17,19–21^ Importantly, nintedanib compromised cell viability of SM patient samples and blocked KIT D816V phosphorylation and signaling in primary patient cells.

Molecular docking studies further corroborated our drug screening data and revealed nintedanib binding preferentially to the ATP binding pocket of inactive KIT D816V. We also identified further nintedanib analogues with KIT D816V selectivity and mapped key residues in the ATP binding pocket for the interaction with the indole moiety of these compounds. This information should be useful for studies on guided chemical enhancement of nintedanib-based compounds by tailoring specific ligand-receptor interactions to increase compound affinity and specificity.

The iPSC-based disease model also enabled us to evaluate the impact of *KIT* D816V on MCs. *KIT* D816V was correlated with more accelerated and more prominent development of MCs during hematopoietic differentiation of iPSCs, in agreement with the abnormal MC expansion and infiltration observed in SM. No morphological differences were observed between *KIT* D816V and control MCs, but *KIT* D816V MCs presented higher expression and surface levels of FCER1A, indicating accelerated MC development and maturation. Most importantly, *KIT* D816V iPSC-derived MCs were also efficiently targeted by nintedanib.

We further exploited our SM iPSC panel to evaluate potential cytotoxic effects of nintedanib on endothelial cells, as this compound was initially developed as an inhibitor for VEGFR, a key receptor to endothelial cell development and function.^36,37^ Interestingly, nintedanib preferentially targeted iPSC-derived hematopoietic cells and MCs over iPSC-derived endothelial cells. These results demonstrate the versatility of the iPSC-based disease model developed in the present work.

One limitation of our SM disease model is the permanent presence of the *KIT* D816V and associated mutations in iPSCs and in all cells derived thereof. Thus, KIT D816V iPSCs represent a snapshot of a particular disease state and their differentiation into KIT D816V hematopoietic progenitors and MCs and does not mirror SM disease progression. In this context, it is currently under debate whether the *KIT* D816V mutation is an early or late event in SM pathogenesis. Jawhar et al identified *KIT* D816V as a late event following *TET2*, *SRSF2* and *ASXL1* mutations.^23^ Grootens et al., by using single cell RNA-Seq of SM patient samples, identified *KIT* D816V in hematopoietic stem cells and also in lineage-primed progenitors and MCs.^48^ More recently, leukemic stem cells for MCL have been reported to reside in the CD34^+^/CD38^−^ fraction of the malignant clone.^49^ In light of these reports our iPSC-based SM disease model stands as a powerful tool for the systematic evaluation of *KIT* D816V impact on the different stages of hematopoiesis.

In summary, we report on the first iPSC-based model for SM and envision an expansion of our library of SM-derived iPSCs with *KIT* D816V and associated mutations to provide an even more comprehensive array of SM disease models. Additionally, nintedanib, identified in this study as effective KIT D816V inhibitor, is already in clinical use and thus should be considered as an additional and/or alternative option for treatment of advanced SM.

## Acknowledgments

We acknowledge the support of U. Gollan for cytogenetic analysis, S. Mitzka for gene expression analysis, the Interdisciplinary Center for Clinical Research Aachen (IZKF Aachen) FACS Core Facility, Faculty of Medicine, RWTH Aachen University, Aachen, Germany, G. Aydin for cell sorting and E. Mierau for expert administrative assistance. The High-Throughput Biomedicine Unit, Institute of Molecular Medicine Finland, Helsinki, Finland is acknowledged for assistance. We would like to thank M. Huber and M. Begemann for discussion and advice. We also thank the support of A. Rajendiran and S. Dreschers for providing healthy donor buffy coat samples. MAST was funded by CAPES-Alexander von Humboldt postdoctoral fellowship (99999.001703/2014-05) and donation by U. Lehmann. MG, NC and MZ were funded by the IZKF Aachen. SM was supported by Finnish Cancer Organizations, Sigrid Juselius Foundation and Finnish special governmental subsidy for health sciences, research and training, Helsinki, Finland. SK was supported by a grant of the Deutsche José Carreras Leukaemie-Stiftung (DJCLS 16 R/2017). PV is supported by the Austrian Science Fund (FWF) - SFB project F4704-B20. Part of this work was supported by funds of ERS, RWTH Aachen University to MZ.

## Author contributions

MAST: designed and performed the experiments and wrote the manuscript. MG: designed and performed the experiments. SS and FK: performed experiments on iPSC generation and provided support on hematopoietic differentiation. KVG and GE: provided patient samples, performed compound screening on primary samples, and provided support on data analysis. KF: performed Western blot experiments. RG and GR: performed molecular docking studies. ASS: provided support on immunohistochemistry analysis. OMJD and SMM: performed high-throughput drug screening experiments and data analysis. AM: performed NGS analysis. HMS: performed cytogenetic analysis. RG and WW: performed DNA methylation analysis. TB: performed histological analysis. JP and AS: provided patient samples and support on NGS data analysis. MJ and AR: provided patient samples. FH and PE: provided nintedanib analogues and support on data analysis. SK and THB: provided patient samples and support on data analysis. PV: provided patient samples, support on data analysis and wrote the manuscript. NC: designed the experiments, performed Western blot analysis and provide support on data analysis. MZ: designed the experiments, analyzed the data and wrote the manuscript. All authors approved the final version of the manuscript for submission.

