## Supplemental Materials and Methods for "Nintedanib Targets KIT D816V Neoplastic Cells Derived from Induced Pluripotent Stem cells of Systemic Mastocytosis"

**Statistics**

All data are presented as means ±SD of at least three independent experiments, unless otherwise indicated. Statistical analysis was performed with GraphPad Prism 7 using t-test or Welch´s t-test. P values ≤0.05 were considered as indicative of statistical significance. IC50 values were calculated by nonlinear regression analysis using GraphPad Prism 7.

**Study approval**

Bone marrow (BM) and/or peripheral blood (PB) samples were obtained from *KIT* D816V, *KIT* D816H and *KIT* S476I SM patients (supplemental Table 1 and 9) after written informed consent (RWTH Aachen University Hospital ethics board reference number EK206/09, Medical University of Vienna ethics board reference number EK1184/2014 and Medical Faculty of Mannheim, Heidelberg University, ethics board reference numbers 2007-087N-MA, 2013-514N-MA and 2013-509N-MA).

**iPSC generation**

iPSCs were generated from BM or PB samples with Oct4, Sox2, Klf4 and cMyc in Sendai virus vectors (CytoTune™-iPS 2.0 Sendai Reprogramming Kit, Thermo Fisher Scientific) ^1^ or with 4-in-1 lentivirus vector ^2^ (Supplemental Table 1). iPSC culture was performed on irradiated mouse embryo fibroblasts (MEF) as previously described.^1^

**Next generation sequencing (NGS) analysis**

Genomic DNA (gDNA) of BM or PB samples and of iPSCs derived thereof was extracted with NucleoSpin Tissue Kit (Macherey-Nagel, Düren, Germany) according to the manufacturer instructions. DNA quantification was done with NanoDrop ND-1000 UV-Vis Spectrophotometer (Thermo Fisher Scientific) and 250 ng gDNA was used for library preparation with a customized, clinically validated panel (TruSeq Custom Amplicon Kit, Illumina) covering 31 genes associated with myeloproliferative disease (*ABL1*, *ASXL1*, *BARD1*, *CALR*, *CBL*, *CHEK2*, *CSF3R*, *DNMT3A*, *ETNK1*, *ETV6*, *EZH2*, *IDH1*, *IDH2*, *JAK2*, *KIT*, *KRAS*, *MPL*, *NF-E2*, *NRAS*, *PDGFRA*, *PTPN11*, *RUNX1*, *SETBP1*, *SF3A1*, *SF3B1*, *SH2B3*, *SRSF2*, *TCF12*, *TET2*, *TP53*, *U2AF1*). Sequencing was conducted on a MiSeq Sequencing System (Illumina). For demultiplexing and FastQ file generation the MiSeq onboard software was used (MiSeq Control Software v2.6 and Real time analysis software v1.18.54, Illumina). Alignment and variant calling were performed with the SeqNext-Module of the SeqPilot-Software (JSI medical systems, Version 4.4.0 Build 509). Variants were called with a bidirectional frequency of >5% and non-synonymous mutations identified were classified as pathogenic (5), likely pathogenic (4), uncertain (3), likely not pathogenic or little clinical significance (2) and not pathogenic or of no clinical significance (1).^3^

**Endothelial cell differentiation**

iPSCs were differentiated into endothelial cells essentially as previously described.^4^ Briefly, iPSCs were treated with Accutase (StemCell Technologies) and 5x10^5^ single cells/well were seeded on Matrigel-coated 6-well plates in StemMACS iPS-Brew XF (Miltenyi Biotech) supplemented with 10 µM Rho-Kinase inhibitor (Y-27632, Abcam). From day 1 to 5, cells were cultured in STEMdiff Mesoderm Induction Medium (StemCell Technologies) from day 6 to 8, in StemPro-34 SFM (Thermo Fisher Scientific) supplemented with 200 ng/ml VEGF (Peprotech) and 2 µM Forskolin (Sigma-Aldrich) and from day 9 to 12 in StemPro-34 SFM supplemented with 50 ng/mL VEGF. Medium change was performed daily. Cells were then harvested by Accutase treatment, selected for CD144 surface expression by MACS (Miltenyi Biotech) and further cultured in Endothelial Cell Growth Medium-2 (EGM-2, Lonza). Endothelial differentiation was monitored by flow cytometry with antibodies specific for CD31, CD34, CD43, CD45, CD105, CD144 and KIT (supplemental Table 4).

**Proliferation assay**

The proliferation assay based on the reduction of MTT (3-(4,5-dimethylthiazol-2-yl)-2,5-diphenyltetrazolium bromide) into purple formazan by cellular NAD(P)H-dependent oxidoreductase, also known as MTT assay, was used to assess the proliferation potential of iPSC-derived KIT^+^ and KIT^-^ hematopoietic cells. Cells were subjected to MACS (Miltenyi Biotech) and 1x10^4^ KIT^+^ or KIT^-^ cells were seeded per well in a flat bottom 96 well-plate in 100 µl of StemPro-34 SFM supplemented with 100 U/ml penicillin, 100 µg/ml streptomycin (all Thermo Fisher Scientific), 100 ng/ml stem cell factor (SCF), 50 ng/ml fms-related tyrosine kinase 3 ligand (FLT3L) and 30 ng/ml interleukin 3 (IL-3, all Peprotech), and 10 ng/ml interleukin 6/soluble interleukin 6 receptor fusion protein (hyper-IL-6).^5^ Alternatively, cells were seeded in medium as described above but without SCF or without any cytokines. Cell proliferation was measured after 24, 48, 72 and 96 h incubation at 37°C and 5% CO_2_ by adding 10 μl of 5 mg/ml MTT solution (Sigma Aldrich) followed by 4 h incubation. Cell lysis was performed by adding 100 µl of 1:50 isopropanol-2M HCl solution (Sigma Aldrich) and absorption was measured at 550 nm.

**LDL uptake assay**

Endothelial cells were grown on gelatin-coated 24-well plates in supplemented EGM-2 medium (Lonza) until full confluency was reached. Cells were then incubated for 4 h with 10 µg/ml Dil-Ac-LDL (Alfa Aesar) in EGM-2 medium at 37°C and 5% CO_2_ and washed 3 times with PBS (without calcium and magnesium, Thermo Fisher Scientific). Nuclei were stained with 5 µg/ml DAPI (Vector Laboratories) for 10 min at RT. Image acquisition was performed on EVOS Fluorescence Digital Inverted Microscope (AMG-Advanced Microscopy Group, WA, USA).

**Flow cytometry analysis and cell sorting**

Hematopoietic differentiation was monitored by staining with specific antibodies and flow cytometry (supplemental Table 4). Briefly, cells were harvested, passed through a 40 µm cell strainer (Falcon) and washed once with PBS (Thermo Fisher Scientific). Blocking of unspecific binding was done with 1% human IgG solution (Privigen, CSL Behring) in PBS supplemented with 1% BSA and 2 mM EDTA for 30 min at 4°C followed by staining with specific antibodies under the same conditions. Single cell suspensions of iPSCs, ESCs or endothelial cells were obtained by Accutase treatment and cells were analyzed by flow cytometry as described above. In some experiments iPSCs and ESCs were stimulated with 250 ng/ml SCF for 15 min prior to FACS analysis. Flow cytometry analysis was performed with a FACS Canto II and FACS sorting with a FACS Aria II 3L (both BD Bioscience) and data analysis was done with FlowJo software (Tree Star).

**Apoptosis assay**

iPSC-derived hematopoietic cells were analyzed for apoptosis by flow cytometry using FITC Annexin V Apoptosis Detection Kit (BD Pharmingen) following manufacturer’s instructions. Briefly, suspension cells were harvested, washed once with PBS and 1x10^5^ cells were resuspended in 100 µl of 1X Binding Buffer followed by addition of 5 µl FITC Annexin V and 5µl of propidium iodide (PI). Samples were incubated at RT for 15 min in the dark, 400 µl 1X Binding Buffer was added and cells were analyzed by flow cytometry using a FACS Canto II (BD Bioscience).

**Western blotting**

Western blot (WB) analysis was performed as previously described^6^ using the specific antibodies listed in supplemental Table 4.

HMC-1.1 and HMC-1.2 cells (5x10^6^) or iPSC-derived HPCs (3-4x10^6^) were cultured in 3 ml RPMI 1640 supplemented with 10% FCS, 2 mM L-glutamine, 100 U/ml penicillin and 100 µg/ml streptomycin (all Thermo Fisher Scientific) and treated with 1 µM nintedanib, midostaurin, avapritinib or ripretinib (all Selleckchem, Munich, Germany) for 4 h. Dimethyl sulfoxide (DMSO, Sigma-Aldrich) treated cells were used as control. Cells were then harvested and processed for WB analysis as above.

Patient 10 MNCs (supplemental Table 1) were cultured in StemSpan SFEM (StemCell Technologies) supplemented with 2 mM L-glutamine, 100 U/ml penicillin, 100 µg/ml streptomycin, 100 ng/ml SCF, 50 ng/ml FLT3L, 20 ng/ml TPO, 10 ng/ml hyper-IL-6 for 24h followed by 4 h starvation in cytokine-free medium. Cells were then treated with 1 µM nintedanib for 4 h and DMSO treated cells were used as controls. Cells were harvested and processed for WB analysis as above.

To investigate the response to SCF stimulation, iPSCs and ESCs were cultured on Matrigel-coated 6-well plates in supplemented StemMACS iPS-Brew XF, treated with 250 ng/ml SCF for 15 min, harvested by Accutase treatment and further processed for WB analysis as above.

**Colony forming unit (CFU) assay**

iPSC and ESC-derived suspension cells of the hemogenic endothelium layer were harvested in regular time intervals (4-5 days), washed with PBS and resuspended in StemPro-34 SFM without cytokine supplementation. Cells (10^4^) were seeded in 1 ml of StemMACS HSC-CFU lite with Epo (Miltenyi Biotech) in a 3.5 cm dish (Greiner Bio-One) and incubated for 14 days at 37°C and 5% CO_2._ Colony classification and counting was performed by microscopy inspection.

**Cytospin/smear preparations**

iPSC and ESC-derived hematopoietic and FACS sorted cells were centrifuged onto glass slides in Shandon Cytospin 4 cytocentrifuge (Thermo Fisher Scientific) followed by fixation with methanol at RT. Cells were stained with Benzidine (Sigma-Aldrich) and Diff Quik (Medion Diagnostics, Düdingen, Switzerland) or with acidic Toluidine Blue O (Sigma-Aldrich). Mounting was performed with Entellan (Merck) and image acquisition was done with Leica DMRX microscope (Leica) and Leica Application Suite software (Leica Microsystems). Image J was used for image handling. Smear preparations of FACS sorted MCs were done on a glass slide followed by acetone fixation at RT. Cells were stained with acidic Toluidine Blue O as described above.

For tryptase staining sorted MCs (cytospin or smear preparations) were fixed in acetone and subjected to immunohistochemical staining against MC tryptase with Flex Kit DAKO/Agilent (DAKO, Carpinteria, CA, USA). Briefly, endogenous peroxidase was blocked with specific blocking reagent and samples were incubated with anti-tryptase monoclonal mouse antibody (M7052, DAKO) for 30 min. Next, the staining enhancer was applied, followed by staining with secondary antibody and visualization with horseradish peroxidase and DAB (DAKO). Counterstaining was done with hematoxylin (DAKO). Slides were then dehydrated, placed in xylene and mounted on coverslips. Image acquisition and handling was performed as described above.

**Drug sensitivity and resistance testing (DSRT) on HMC1.1 and HMC1.2 cells**

A library of 459 FDA/EMA approved or investigational anti-cancer and other drugs was used (supplemental Table 8). All compounds were purchased from commercial chemical vendors and dissolved in either DMSO or water. Each compound was pre-printed on 384-well plates (Corning) in five different concentrations covering a 10,000-fold concentration range with an acoustic liquid handling device (Echo 550, Labcyte Inc.). Compounds were dissolved in 5 µl culture medium on a shaker for 10 min. 20 µl of single-cell suspensions of HMC-1.1 or HMC-1.2 cells (2,500 cells per well) in culture medium as above were dispensed using a Multi-Drop Combi peristaltic dispenser (Thermo Fisher Scientific). Plates were incubated at 37°C and 5% CO_2_ for 72 h and cell viability was measured with CellTiter-Glo 2.0 reagent (Promega) according to the instructions of the manufacturer with a Pherastar FS plate reader (BMG Labtech). Cell viability luminescence data were normalized to DMSO-only wells (negative control) and 100 mM benzethonium chloride-containing wells (positive control). Data were quantified using the drug sensitivity score (DSS).^7^

Alternatively, HMC-1.1 and HMC-1.2 cells (10^4^ cells/well) were seeded in white flat bottom 96 well-plates (Greiner) in 90 µl of compound screening medium (RPMI 1640 supplemented with 10% FCS, 2 mM L-glutamine, 100 U/ml penicillin and 100 µg/ml streptomycin). Compounds were dissolved in DMSO at stock concentrations of 10 mM and diluted to concentrations ranging from 1 nM to 10 µM in compound screening medium. Cells were incubated with compounds for 66 h and cell viability was determined with CellTiter-Glo Luminescent Cell Viability Assay (Promega). Fluorescence measurement was performed with SpectraMAX i3 Plate Reader and Softmax Pro Software (Molecular Devices).

**Compound testing on primary samples**

PB or BM mononuclear cells (MNCs) from SM patients were obtained by Ficoll density gradient centrifugation. Buffy coat samples from 8 healthy donors were used as control MNCs. Cells were expanded in StemSpan SFEM (StemCell Technologies) supplemented with 2 mM L-glutamine, 100 U/ml penicillin, 100 µg/ml streptomycin, 100 ng/ml SCF, 50 ng/ml FLT3L, 20 ng/ml TPO, 10 ng/ml hyper-IL-6 for 24-48 h. When indicated primary MNCs were enriched for KIT expressing cells by MACS using CD117 MicroBead Kit (Miltenyi Biotec). MNCs were then seeded in 96-well plates (10^4^ cells/well) in compound screening medium (see above) supplemented with 1 µM nintedanib or midostaurin and incubated at 37°C and 5% CO_2_. DMSO treated cells were used as control. After 66 h cell viability was measured with CellTiter-Glo Luminescent Cell Viability Assay.

MNCs of SM patients 25-29 (supplemental Table 9) were treated with 1 µM nintedanib for 48 h and cell proliferation was measured by incubation with ^3^H-thymidine (0.5 µCi) for 16 h. Cells were then harvested on filter membranes and bound radioactivity was measured in a β-counter (Top-Count NXT, Packard Bioscience). All measurements were performed in triplicates and untreated cells were used as control.

For the measurement of nintedanib effect on *KIT* D816V allele burden by RT-PCR analysis, MNCs from SM patients were expanded as described above. Cells were then seeded in compound screening medium (see above) supplemented with 1 µM nintedanib and incubated at 37°C and 5% CO_2_ for 48 h. Cells were harvested and processed for RNA isolation as described (RT-PCR analysis section).

**RT-PCR analysis**

RNA isolation was performed with NucleoSpin RNA Kit (Macherey-Nagel) or MagMAX 96 Total RNA Isolation Kit (Thermo Fisher Scientific) following the instructions of the manufacturers. cDNA was synthesized using MultiScribe reverse transcriptase (High Capacity cDNA Reverse Transcriptase Kit, Thermo Fisher Scientific). RT-qPCR was performed on StepOnePlus Real Time cycler using FAST SYBR Green master mix (Thermo Fisher Scientific). Primers are listed in supplemental Table 11 and were synthesized by Eurofins Genomics, Ebersberg, Germany. Gene expression data were normalized to GAPDH expression levels, subjected to hierarchical clustering (bidirectional clustering considering Euclidian distance measure and average linkage method) and are represented in heatmap format with dendrograms. Data analysis and heatmap plots were done with MeV-Multiple Experiment Viewer (http://mev.tm4.org/).

**Molecular Docking**

The modeling study has been entirely performed using the Schrodinger 2019-1suite. All the ligands were structurally pre-processed using LigPrep tool of the suite. Two different isomeric forms were found at pH 7 for each ligand.

Molecular docking was conducted on the following KIT structures: unmutated (PDB code 3G0E) and D816V KIT. The latter was produced starting from the structure of D816H KIT (PDB code 3G0F) replacing histidine with valine at the mutation site. Both the unmutated and the mutant proteins are resolved in complex with Sunitinib and correspond to inactive, autoinhibited geometries. Most of the analyzed ligands are close in chemical structure to nintedanib, which is crystalized within the vascular endothelial growth factor receptor 2 (VEGFR2) kinase domain (PDB code 3C7Q). The smallest RMSD with the KIT systems and the VEGFR2 was obtained for 3G0E and 3G0F and therefore those structures were used for initial docking of the ligands on KIT (supplemental Figure 18A).

Protein structures were pre-processed with the Protein Preparation Wizard within the Schrodinger suite. The protonation states of each side chain were generated using the software Epik for pH 7 in Schroedinger 2019-2. Protein energy minimization was performed using the OPLS3e force field.^8^

Glide^9^ was used for all docking calculations. Ligands are often known to induce conformational changes in the active site upon binding. We therefore used the Schrödinger induced fit docking (IFD) protocol of the suite to account for these changes. The receptor grid center was built around the co-crystallized ligands found in the X-ray structures. In the first stage of the IFD protocol, softened-potential docking is performed to generate 20 initial poses. The scaling factors to soften the potentials of the receptors and ligands were set to 0.7 and 0.5, respectively. A 2.5 kcal/mol energy window was used for ligand conformational sampling. For each of the top 20 poses (with respect to Glide Score) from the initial softened-potential docking step, all residues within 7.0 Å of ligand poses were refined using the Prime molecular dynamics.

Refinement was performed with the Prime package and the OPLS3e force field^8^ to accommodate the ligand by reorienting nearby side chains. The complexes were ranked by Prime energy (molecular mechanics plus solvation) and those within 20 kcal/mol of the minimum energy structure were passed through for a final round of Glide docking and scoring. The ligands were then re-docked into their corresponding receptor structures using XP scoring in Glide. Each docking result was analyzed by comparing the Glide Score.^10^

For each ligand, all the poses generated were visually inspected to eliminate binding geometries not compatible with experimental evidences and producing artificially high values of Glide Score. For all the poses retained, the average value of the Glide Score was computed together with its standard error.

Limitations: As with any modelling study, our models also have limitations. First, we used a single static structure for each KIT state and the performance of a single protein conformation in docking is not always reliable. We partially overcame this limitation by performing induced fit docking. The latter simulates structural changes occurring upon ligand binding, mimicking protein flexibility at the binding pocket. Protein mobility is also most probably increased by the presence of moving and displaceable water molecules. A second limitation is that we do not have explicit water molecules, which are also critical in binding process. Solvation effects can be accounted up to 100-fold difference in binding affinity (corresponding to ~3 kcal/mol in binding free energy). However, the molecular docking software we used, Glide, is able to consider the presence of water molecules at least to some extent, by including statistics about the number of hydrogen bonds formed by polar and apolar groups and an implicit solvent model. Finally, we rely on scoring functions to rank and select the best binding poses. Current docking/scoring methods were suggested to provide reasonable predictions of ligand binding modes, but their performance is often disappointing in predicting ligand binding affinities. Additionally, those methods are often system-dependent, making it very hard to decide which scoring function is suitable for the chosen target protein. Here, to extract the best binding pose for each ligand, Glide score was used.

**Supplemental Tables:**

**Supplemental Table 1.** **KIT D816V patient samples used for iPSC reprogramming.**

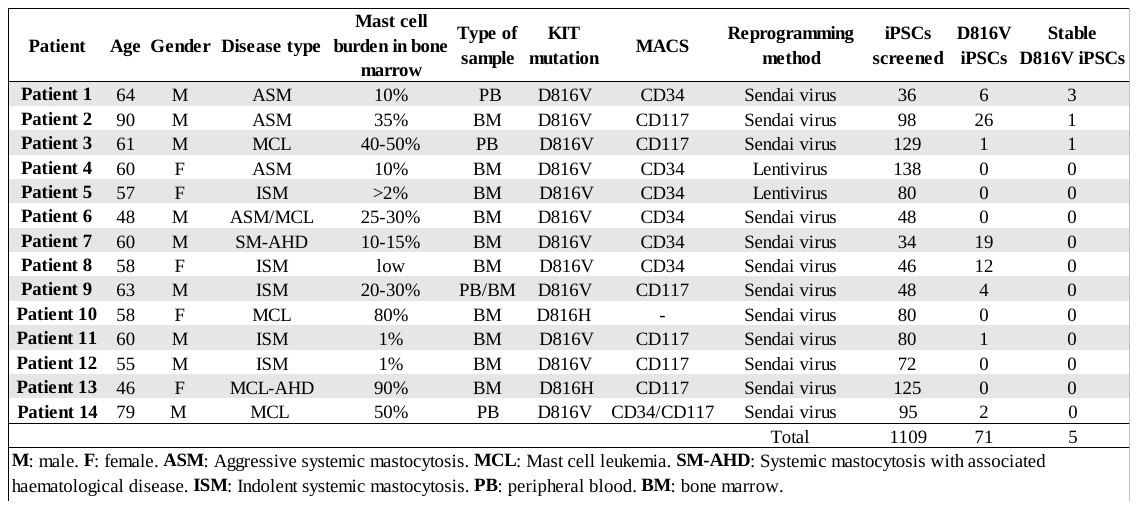

Overview of SM patient samples used for iPSC reprogramming and number of iPSC clones obtained.

**Supplemental Table 2.** **Oligonucleotides for PCR analysis.**
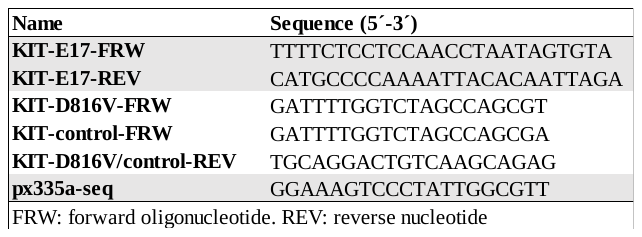

**Supplemental Table 3.** **Oligonucleotides for CRISPR/Cas9n gRNA cloning into pX335a vector and donor template.**

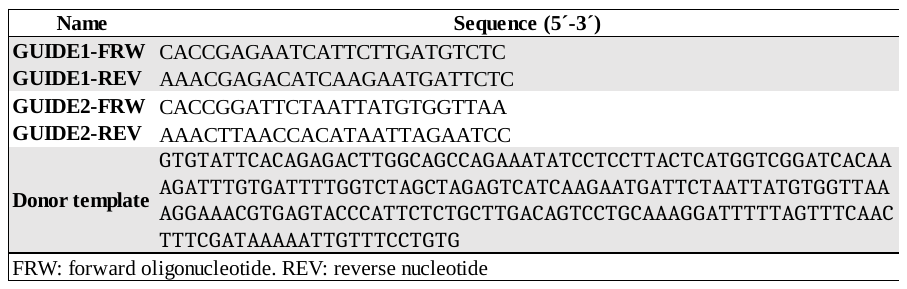

**Supplemental Table 4.** **List of antibodies.**

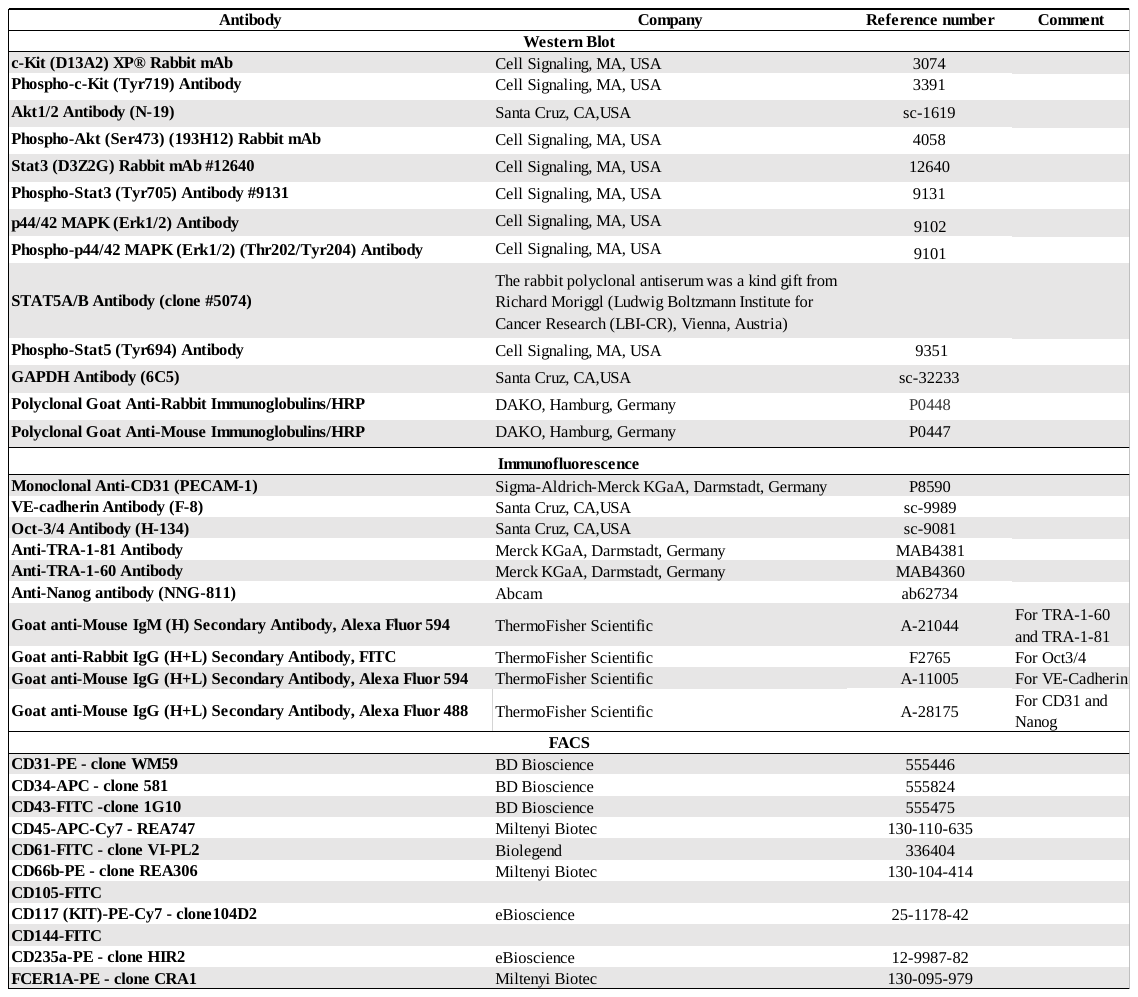

Antibodies used in the present work for Western blot, immunofluorescence and FACS analysis.

**Supplemental Table 5. Patient 1 mutations detected by NGS and their allele frequency in primary sample and in iPSCs derived thereof (KIT D816V 1-3 and control 1 and 2).**

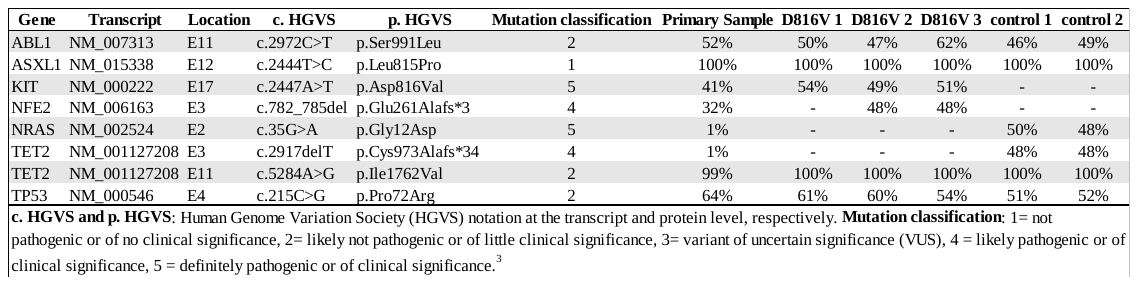

**Supplemental Table 6. Patient 2 mutations detected by NGS and their allele frequency in primary sample and in iPSCs derived thereof (KIT D816V 1 and control 1).**

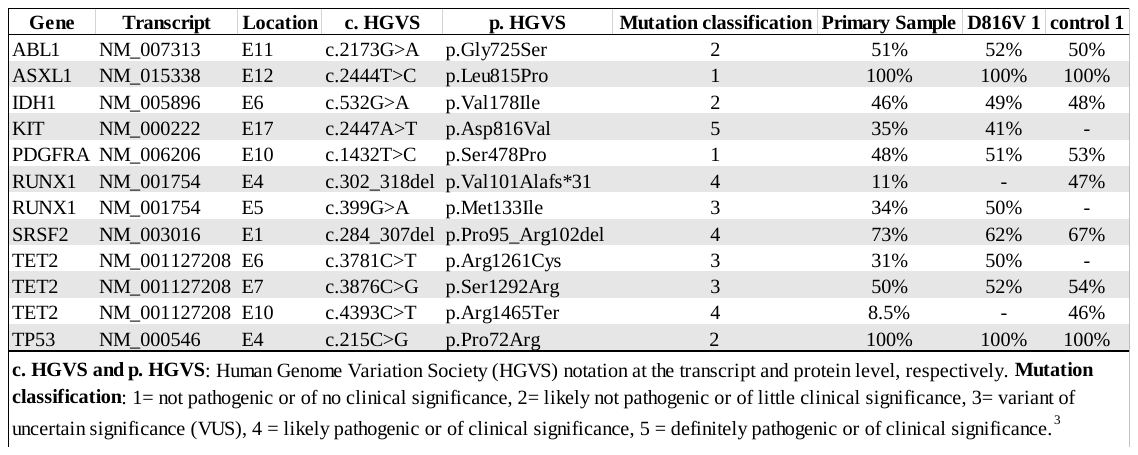

**Supplemental Table 7.** **Patient 3 mutations detected by NGS and their allele frequency in primary sample and in iPSCs derived thereof (KIT D816V 1 and control 1 and 2).**

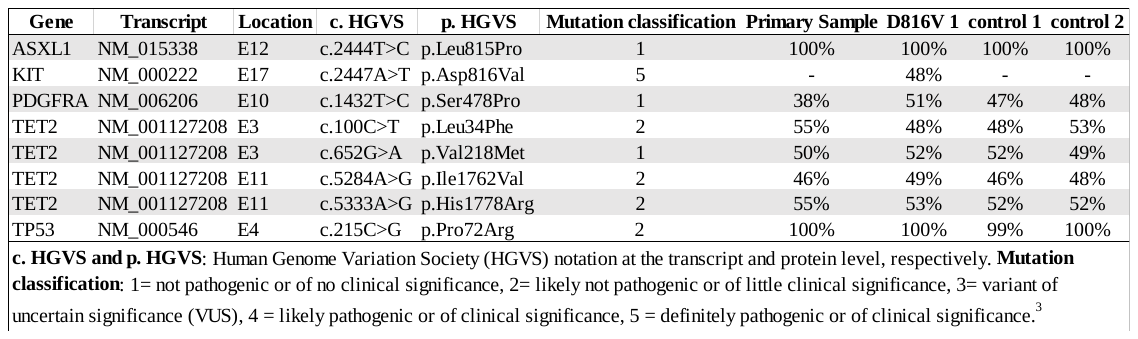

**Supplemental Table 8.** **List of compounds tested on HMC-1 cell lines.**

| **Drug name** | **Mechanism/Targets** | **Class** |
| --- | --- | --- |
| **GSK-461364** | PLK1 inhibitor | B. Kinase inhibitor |
| **Gemcitabine** | Antimetabolite; Nucleoside analog | A. Conv. Chemo |
| **Cytarabine** | Anti-metabolite, interferes with DNA synthesis | A. Conv. Chemo |
| **Clofarabine** | Anti-metabolite; Purine analog | A. Conv. Chemo |
| **BI 2536** | PLK1 inhibitor | B. Kinase inhibitor |
| **RG7388** | MDM2/p53, 2nd gen MDM2 inhibitor | G. Apoptotic modulator |
| **PF-03758309** | PAK inhibitor | B. Kinase inhibitor |
| **ARRY-520** | KSP/Eg5 inhibitor | I. Kinesin inhibitor |
| **Cladribine** | Anti-metabolite; Purine analog | A. Conv. Chemo |
| **Pevonedistat** | NAE inhibitor | H. Metabolic modifier |
| **Sirolimus** | binds FKBP12, causes inhibition of mTORC1 | C. Rapalog |
| **MK1775** | Wee1 inhibitor | B. Kinase inhibitor |
| **GSK-2334470** | PDK1 inhibitor | B. Kinase inhibitor |
| **Patupilone** | Mitotic inhibitor, epothilone microtubule stabilizer | A. Conv. Chemo |
| **SB 743921** | Mitotic inhibitor. Eg5/KSP inhibitor | I. Kinesin inhibitor |
| **PHA-793887** | CDK inhibitor | B. Kinase inhibitor |
| **Topotecan** | Topoisomerase I inhibitor. Camptothecin analog | A. Conv. Chemo |
| **Temsirolimus** | binds FKBP12, causes inhibition of mTORC1 | C. Rapalog |
| **Birinapant** | IAPs, SMAC mimetic | G. Apoptotic modulator |
| **Volasertib** | PLK1 inhibitor | B. Kinase inhibitor |
| **Everolimus** | binds FKBP12, causes inhibition of mTORC1 | C. Rapalog |
| **Ridaforolimus** | binds FKBP12, causes inhibition of mTORC1 | C. Rapalog |
| **Dinaciclib** | CDK inhibitor | B. Kinase inhibitor |
| **Rigosertib** | PLK1 inhibitor, non-ATP-comp inhibitor | B. Kinase inhibitor |
| **CH-5132799** | pan-PI3K inhibitor | B. Kinase inhibitor |
| **AZD-8186** | PI3Kbeta inhibitor | B. Kinase inhibitor |
| **UCN-01** | PKCbeta, PDK1, Chk, Cdk2 inhibitor | B. Kinase inhibitor |
| **Paclitaxel** | Mitotic inhibitor, taxane microtubule stabilizer | A. Conv. Chemo |
| **Mitoxantrone** | Topoisomerase II inhibitor | A. Conv. Chemo |
| **Daunorubicin** | Topoisomerase II inhibitor | A. Conv. Chemo |
| **Daporinad** | NAMPT inhibitor | H. Metabolic modifier |
| **AZD-6482** | PI3Kbeta-selective inhibitor | B. Kinase inhibitor |
| **Floxuridine** | Antimetabolite; Analog of 5-fluorouracil | A. Conv. Chemo |
| **AR-42** | HDAC inhibitor | E. Differentiating/ epigenetic modifier |
| **Vinblastine** | Mitotic inhibitor. Vinca alkaloid microtubule depolymerizer | A. Conv. Chemo |
| **NVP-BGT226** | PI3K/mTOR inhibitor | B. Kinase inhibitor |
| **Camptothecin** | Topoisomerase I inhibitor | A. Conv. Chemo |
| **AZD1208** | PIM1, 2, 3 kinase inhibitor | B. Kinase inhibitor |
| **TGX-221** | p110beta selective PI3K inhibitor | B. Kinase inhibitor |
| **8-chloro-adenosine** | Nucleoside analog, RNA synthesis inhibitor | A. Conv. Chemo |
| **Bryostatin 1** | PKC activator | B. Kinase inhibitor |
| **Belinostat** | HDAC inhibitor | E. Differentiating/ epigenetic modifier |
| **CUDC-101** | HDAC & EGFR, Her2 inhibitor | B. Kinase inhibitor |
| **Pracinostat** | HDAC inhibitor | E. Differentiating/ epigenetic modifier |
| **Pimasertib** | MEK inhibitor | B. Kinase inhibitor |
| **Cobimetinib** | MEK1/2 inhibitor | B. Kinase inhibitor |
| **Abemaciclib** | CDK4 and 6 inhibitor | B. Kinase inhibitor |
| **Triapine** | ribonucleotide reductase inhibitor | H. Metabolic modifier |
| **MK-2206** | AKT inhibitor | B. Kinase inhibitor |
| **Lenalidomide** | Immunomodulatory | D. Immunomodulatory |
| **Selinexor** | CRM1 inhibitor | G. Apoptotic modulator |
| **KX2-391** | non-ATP competitive Src inhibitor | B. Kinase inhibitor |
| **Idelalisib** | PI3K inhibitor, p110δ-selective | B. Kinase inhibitor |
| **CUDC-907** | HADC1/2/3/10, PI3Kalpha inhibitor | E. Differentiating/ epigenetic modifier |
| **Idarubicin** | Topoisomerase II inhibitor | A. Conv. Chemo |
| **Quisinostat** | HDAC inhibitor | E. Differentiating/ epigenetic modifier |
| **GSK2636771** | p110beta selective PI3K inhibitor | B. Kinase inhibitor |
| **MK-8745** | Aurora A inhibitor | B. Kinase inhibitor |
| **GSK-690693** | AKT, PKA, PKC inhibitor | B. Kinase inhibitor |
| **Mocetinostat** | HDAC inhibitor (HDAC1 & 2-selective) | E. Differentiating/ epigenetic modifier |
| **SNS-032** | Cdk inhibitor | B. Kinase inhibitor |
| **Duvelisib** | PI3K inhibitor | B. Kinase inhibitor |
| **AZD-8330** | MEK inhibitor | B. Kinase inhibitor |
| **Nutlin-3** | mdm2 inhibitor | G. Apoptotic modulator |
| **GSK-1059615** | PI3K/mTOR inhibitor | B. Kinase inhibitor |
| **GDC-0623** | MEK1/2 inhibitor | B. Kinase inhibitor |
| **Etoposide** | Topoisomerase II inhibitor | A. Conv. Chemo |
| **LY-294002** | PI3K inhibitor | B. Kinase inhibitor |
| **Vinorelbine** | Mitotic inhibitor. Vinca alkaloid microtubule depolymerizer | A. Conv. Chemo |
| **SCH772984** | ERK1 & 2 inhibitor | B. Kinase inhibitor |
| **Selumetinib** | MEK inhibitor | B. Kinase inhibitor |
| **Panobinostat** | HDAC inhibitor | E. Differentiating/ epigenetic modifier |
| **Vincristine** | Mitotic inhibitor. Vinca alkaloid microtubule depolymerizer | A. Conv. Chemo |
| **2-methoxyestradiol** | Angiogenesis inhibitor | X. Other |
| **Pomalidomide** | Immunomodulatory agent, anti-angiogenic | D. Immunomodulatory |
| **Entinostat** | HDAC inhibitor | E. Differentiating/ epigenetic modifier |
| **Ruboxistaurin** | PKCbeta inhibitor | B. Kinase inhibitor |
| **Methotrexate** | Antimetabolite; Anti-folate agent | H. Metabolic modifier |
| **TAK-733** | MEK inhibitor | B. Kinase inhibitor |
| **Trametinib** | MEK1/2 inhibitor | B. Kinase inhibitor |
| **Teniposide** | Topoisomerase II inhibitor | A. Conv. Chemo |
| **Rocilinostat** | HDAC-6 selective inhibitor | E. Differentiating/ epigenetic modifier |
| **Doxorubicin** | Topoisomerase II inhibitor | A. Conv. Chemo |
| **MLN-8054** | AURa AURb FLT3 KIT (PDGFR) | B. Kinase inhibitor |
| **Mitomycin C** | Antineoplastic anatibiotic; DNA crosslinker | A. Conv. Chemo |
| **Amonafide** | Topoisomerase II inhibitor / DNA intercalator | A. Conv. Chemo |
| **AVN944** | IMPDH inhibitor | H. Metabolic modifier |
| **Oxaliplatin aq** | Platinum-based antineoplastic agent | A. Conv. Chemo |
| **Serdemetan** | HDM2-p53 antagonist | G. Apoptotic modulator |
| **Lestaurtinib** | FLT3, JAK2, TrkA, TrkB, TrkC inhibitor | B. Kinase inhibitor |
| **Docetaxel** | Mitotic inhibitor, taxane microtubule stabilizer | A. Conv. Chemo |
| **Bleomycin** | Glycopeptide antibiotic; causes DNA breaks | A. Conv. Chemo |
| **PF-04691502** | PI3K/mTOR inhibitor | B. Kinase inhibitor |
| **RG-7603** | pan-PI3K inhibitor | B. Kinase inhibitor |
| **Pictilisib** | PI3K inhibitor, pan-class I | B. Kinase inhibitor |
| **AMG-900** | pan-Aurora inhibitor | B. Kinase inhibitor |
| **Dactolisib** | PI3K/mTOR inhibitor | B. Kinase inhibitor |
| **PD0325901** | MEK1/2 inhibitor | B. Kinase inhibitor |
| **PKI-402** | PI3K/mTOR inhibitor | B. Kinase inhibitor |
| **Tamoxifen** | Estrogen receptor antagonist | F. Hormone therapy |
| **Resminostat** | HDAC1, 3, 6 inhibitor | E. Differentiating/ epigenetic modifier |
| **AZD-5438** | CDK1,2,9 inhibitor | B. Kinase inhibitor |
| **PHA 408** | IKK-2 inhibitor | B. Kinase inhibitor |
| **Chloroquine aq** | Antimalaria agent; chemo/radio sensitizer | A. Conv. Chemo |
| **Gedatolisib** | PI3K/mTOR inhibitor | B. Kinase inhibitor |
| **Voxtalisib** | mTOR/PI3K inhibitor | B. Kinase inhibitor |
| **Tretinoin** | Retinoic acid receptor agonist | E. Differentiating/ epigenetic modifier |
| **NVP-LEE011** | CDK4/6 inhibitor | B. Kinase inhibitor |
| **Omacetaxine** | Protein synthesis inhibitor (80 S ribosome) | A. Conv. Chemo |
| **UNC0642** | G9a/GLP inhibitor | E. Differentiating/ epigenetic modifier |
| **Triciribine** | Akt inhibitor | B. Kinase inhibitor |
| **Pelitinib** | EGFR inhibitor | B. Kinase inhibitor |
| **Stattic** | STAT3 SH2 domain inhibitor | X. Other |
| **Imiquimod** | Immunomodulatory agent | D. Immunomodulatory |
| **Vorinostat** | HDAC inhibitor | E. Differentiating/ epigenetic modifier |
| **Indibulin** | Mitotic inhibitor. Microtubule depolymerizer | A. Conv. Chemo |
| **PD184352** | MEK1/2 inhibitor | B. Kinase inhibitor |
| **GDC-0068** | Akt inhibitor | B. Kinase inhibitor |
| **ABT-751** | Mitotic inhibitor. Colchicine site binding microtubule depolymerizer. | A. Conv. Chemo |
| **C646** | p300/CREB-binding protein (CBP) inhibitor | E. Differentiating/ epigenetic modifier |
| **Alpelisib** | PI3Kalpha inhibitor | B. Kinase inhibitor |
| **Milciclib** | CDK2 inhibitor | B. Kinase inhibitor |
| **Tubastatin A** | HDAC6 inhibitor | E. Differentiating/ epigenetic modifier |
| **KU-60019** | ATM inhibitor | B. Kinase inhibitor |
| **AZD8055** | mTOR inhibitor | B. Kinase inhibitor |
| **ZSTK474** | p110gamma selective PI3K inhibitor | B. Kinase inhibitor |
| **Apitolisib** | PI3K/mTOR inhibitor | B. Kinase inhibitor |
| **Irinotecan** | Topoisomerase I inhibitor. Camptothecin prodrug analog | A. Conv. Chemo |
| **Bexarotene** | Antineoplastic agent; retinoid specifically selective for retinoid X receptors | E. Differentiating/ epigenetic modifier |
| **Mepacrine aq** | Unclear. PLA2 inhibitor. NF-kB inhibitor, p53 activator | X. Other |
| **Refametinib** | MEK1/2 inhibitor | B. Kinase inhibitor |
| **MK-8776** | CHEK1 inhibitor | B. Kinase inhibitor |
| **Aldoxorubicin** | Topo II, albumin | A. Conv. Chemo |
| **GSK-1070916** | AURb, AURc inhibitor | B. Kinase inhibitor |
| **Talazoparib** | PARP1/2 inhibitor | E. Differentiating/ epigenetic modifier |
| **StemRegenin 1** | AHR antagonist, stem cell regenerating | E. Differentiating/ epigenetic modifier |
| **NSC348884** | NPM1 oligomerization inhibitor | X. Other |
| **Sotrastaurin** | PKC inhibitor | B. Kinase inhibitor |
| **UNC0638** | G9a/GLP inhibitor | E. Differentiating/ epigenetic modifier |
| **XL-647** | EGFR, ERBB2, VEGFR, EPHB4 | B. Kinase inhibitor |
| **Rabusertib** | Chk1 inhibitor | B. Kinase inhibitor |
| **Radicicol** | HSP90 inhibitor | K. HSP inhibitor |
| **Canertinib** | pan-ErbB inhibitor | B. Kinase inhibitor |
| **PAC-1** | procaspase-3 activator | G. Apoptotic modulator |
| **Binimetinib** | MEK inhibitor | B. Kinase inhibitor |
| **INK128** | mTOR inhibitor | B. Kinase inhibitor |
| **Alvocidib** | Cdk inhibitor | B. Kinase inhibitor |
| **TG100-115** | PI3K gamma/delta inhibitor | B. Kinase inhibitor |
| **LCL161** | IAPs, SMAC mimetic | G. Apoptotic modulator |
| **Fludarabine** | Antimetabolite; Purine analog | A. Conv. Chemo |
| **VX-11E** | ERK1 & 2 inhibitor | B. Kinase inhibitor |
| **Pilaralisib** | PI3K inhibitor. Pan-class I | B. Kinase inhibitor |
| **8-amino-adenosine** | Nucleoside analog, RNA synthesis inhibitor | A. Conv. Chemo |
| **Valrubicin** | Topoisomerase II inhibitor | A. Conv. Chemo |
| **Clomifene** | Selective estrogen receptor modulator | F. Hormone therapy |
| **AZD2014** | mTOR inhibitor, ATP-competitive | B. Kinase inhibitor |
| **OSI-027** | mTOR inhibitor | B. Kinase inhibitor |
| **4-hydroxytamoxifen** | Selective estrogen receptor modulator | F. Hormone therapy |
| **Palbociclib** | Cdk inhibitor (Cdk4/6) | B. Kinase inhibitor |
| **Sonolisib** | PI3K inhibitor, pan-class I. Irreversible | B. Kinase inhibitor |
| **Galiellalactone** | STAT3-DNA interaction inhibitor | X. Other |
| **Itraconazole** | antifungal, hedgehog signaling inhibitor | X. Other |
| **Seliciclib** | CDK2/7/9 inhibitor | B. Kinase inhibitor |
| **GSK343** | EZH2 inhibitor | E. Differentiating/ epigenetic modifier |
| **OSU-03012** | PDPK1 inhibitor | B. Kinase inhibitor |
| **Azacitidine** | DNMT inhibitor | E. Differentiating/ epigenetic modifier |
| **IOX-2** | PHD2 inhibitor | E. Differentiating/ epigenetic modifier |
| **AT-406** | XIAP, cIAP1, cIAP2 inhibitor | G. Apoptotic modulator |
| **PF-00477736** | Chk1 inhibitor | B. Kinase inhibitor |
| **Navitoclax** | Bcl-2 inhibitor | G. Apoptotic modulator |
| **Afatinib** | EGFR inhibitor | B. Kinase inhibitor |
| **Thioguanine** | Antimetabolite; Purine analog | A. Conv. Chemo |
| **Rucaparib** | PARP inhibitor | E. Differentiating/ epigenetic modifier |
| **Deferoxamine aq** | Iron chelator | X. Other |
| **Tipifarnib** | Farnesyltransferase inhibitor | E. Differentiating/ epigenetic modifier |
| **OTX015** | BRD2, 3, 4 | E. Differentiating/ epigenetic modifier |
| **Dactinomycin** | RNA and DNA synthesis inhibitor | A. Conv. Chemo |
| **PF-670462** | CK1epsilon and CK1delta inhibitor | B. Kinase inhibitor |
| **Fingolimod** | S1PR antagonist | X. Other |
| **Abiraterone** | P450 17alpha-hydroxylase-17,20-lyase inhibitor | F. Hormone therapy |
| **PS-1145** | IKK-2 inhibitor | B. Kinase inhibitor |
| **RGFP966** | HDAC3 inhibitor | E. Differentiating/ epigenetic modifier |
| **Valproic acid aq** | HDAC inhibitor | E. Differentiating/ epigenetic modifier |
| **I-BET151** | BET family inhibitor | E. Differentiating/ epigenetic modifier |
| **Cisplatin aq** | Platinum-based antineoplastic agent | A. Conv. Chemo |
| **Neratinib** | EGFR inhibitor | B. Kinase inhibitor |
| **Olaparib** | PARP inhibitor | E. Differentiating/ epigenetic modifier |
| **Plicamycin** | RNA synthesis inhibitor | A. Conv. Chemo |
| **Tivantinib** | MET inhibitor | B. Kinase inhibitor |
| **Ceritinib** | ALK inhibitor | B. Kinase inhibitor |
| **GSK-J4** | JMJD3 (histone demethylase) inhibitor | E. Differentiating/ epigenetic modifier |
| **Auranofin** | Antirheumatic agent | A. Conv. Chemo |
| **BX-912** | PDK1 inhibitor | B. Kinase inhibitor |
| **AT7519** | CDK1, 2, 4, 6 and 9 inhibitors | B. Kinase inhibitor |
| **PF-00562271** | FAK inhibitor | B. Kinase inhibitor |
| **CUDC-305** | HSP90 inhibitor | K. HSP inhibitor |
| **GSK650394** | SGK1 & 2 inhibitor | B. Kinase inhibitor |
| **PF 431396** | FAK/PYK2 inhibitor | B. Kinase inhibitor |
| **AT 101** | Bcl family inhibitor | G. Apoptotic modulator |
| **PFI-1** | Selective chemical probe for BET Bromodomains | E. Differentiating/ epigenetic modifier |
| **Raloxifene** | Selective estrogen receptor modulator | F. Hormone therapy |
| **AZD-5363** | AKT inhibitor | B. Kinase inhibitor |
| **Niraparib** | PARP inhibitor | E. Differentiating/ epigenetic modifier |
| **Fasudil** | Rho kinase, PKA, PKG, PRK inhibitor, prodrug | B. Kinase inhibitor |
| **Ixabepilone** | Mitotic inhibitor. Epothilone microtubule stabilizer. | A. Conv. Chemo |
| **Lonafarnib** | Farnesyl transferase inhibitor | E. Differentiating/ epigenetic modifier |
| **Decitabine** | Nucleoside analog DNA methyl transferase inhibitor | E. Differentiating/ epigenetic modifier |
| **Fluorouracil** | Antimetabolite | A. Conv. Chemo |
| **Obatoclax** | Bcl2 inhibitor | G. Apoptotic modulator |
| **Buparlisib** | PI3K inhibitor, pan-class I | B. Kinase inhibitor |
| **Tanzisertib** | JNK1, 2, 3 inhibitors | B. Kinase inhibitor |
| **Thalidomide** | Immunosuppressant | D. Immunomodulatory |
| **Chlorambucil** | Nitrogen mustard alkylating agent | A. Conv. Chemo |
| **E7438** | EZH2 inhibitor | E. Differentiating/ epigenetic modifier |
| **AZD1480** | JAK1/2, FGFR inhibitor | B. Kinase inhibitor |
| **Veliparib** | PARP inhibitor | E. Differentiating/ epigenetic modifier |
| **Lomeguatrib** | O6-methylguanine-DNA methyltransferase inhibitor | E. Differentiating/ epigenetic modifier |
| **FG-4592** | HIF prolyl hydroxylase inhibitor | E. Differentiating/ epigenetic modifier |
| **Dacarbazine** | Alkylating agent | A. Conv. Chemo |
| **Talmapimod** | p38alpha selective inhibitor | B. Kinase inhibitor |
| **Celecoxib** | Selective COX-2 inhibitor | J. NSAID |
| **Crizotinib** | ALK, c-Met inhibitor | B. Kinase inhibitor |
| **ARRY-380** | HER2 inhibitor | B. Kinase inhibitor |
| **Mechlorethamine** | Nitrogen mustard alkylating agent | A. Conv. Chemo |
| **Pilocarpine** | Non-selective muscarinic receptor agonist | X. Other |
| **Metformin aq** | AMPK activator | H. Metabolic modifier |
| **Anagrelide** | PDE-3, PLA2 inhibitor | X. Other |
| **Goserelin** | Gonadotropin releasing hormone super agonist | F. Hormone therapy |
| **Rofecoxib** | COX-2 inhibitor | J. NSAID |
| **Lapatinib** | HER2, EGFR inhibitor | B. Kinase inhibitor |
| **Letrozole** | Aromatase inhibitor | F. Hormone therapy |
| **Anastrozole** | Aromatase inhibitor | F. Hormone therapy |
| **Bicalutamide** | Nonsteroidal antiandrogen | F. Hormone therapy |
| **Lomustine** | Alkylating nitrosourea compound | A. Conv. Chemo |
| **Altretamine** | Formaldehyde release, alkylating agent | A. Conv. Chemo |
| **Aminoglutethimide** | Anti-steroid, aromatase inhibitor | F. Hormone therapy |
| **Cyclophosphamide** | Alkylating agent | A. Conv. Chemo |
| **Finasteride** | type II 5-alpha reductase inhibitor | F. Hormone therapy |
| **Flutamide** | Nonsteroidal antiandrogen | F. Hormone therapy |
| **Ifosfamide** | Nitrogen mustard alkylating agent | A. Conv. Chemo |
| **Levamisole** | Immunomodulatory agent | D. Immunomodulatory |
| **Melphalan aq** | Nitrogen mustard alkylating agent | A. Conv. Chemo |
| **Procarbazine** | Alkylating agent | A. Conv. Chemo |
| **Temozolomide** | Alkylating agent | A. Conv. Chemo |
| **Fulvestrant** | Estrogen receptor antagonist | F. Hormone therapy |
| **Nilutamide** | Nonsteroidal antiandrogen | F. Hormone therapy |
| **Mitotane** | Antineoplastic agent | A. Conv. Chemo |
| **Allopurinol** | Xanthine oxidase inhibitor | A. Conv. Chemo |
| **Busulfan** | Alkylating antineoplastic agent | A. Conv. Chemo |
| **Hydroxyurea** | Antineoplastic agent | A. Conv. Chemo |
| **Carmustine** | Alkylating agent | A. Conv. Chemo |
| **Pipobroman** | Alkylating agent | A. Conv. Chemo |
| **Pravastatin aq** | HMG CoA reductase inhibitor | H. Metabolic modifier |
| **Perifosine aq/PBS** | AKT/PI3K inhibitor | B. Kinase inhibitor |
| **Tarenflurbil** | Gamma-secretase inhibitor | X. Other |
| **Bimatoprost** | Prostaglandin analog | D. Immunomodulatory |
| **Tacedinaline** | HDAC inhibitor | E. Differentiating/ epigenetic modifier |
| **Sonidegib** | Smoothened (Hh) inhibitor | X. Other |
| **Iniparib** | PARP inhibitor | E. Differentiating/ epigenetic modifier |
| **Nelarabine** | Nucleoside analog, DNA, RNA synth inhibitor | A. Conv. Chemo |
| **Plerixafor aq** | CXCR4 antagonist | X. Other |
| **EMD1214063** | c-Met inhibitor | B. Kinase inhibitor |
| **Tofacitinib** | JAK3, JAK2(V617F) inhibitor | B. Kinase inhibitor |
| **Enzastaurin** | PKCbeta inhibitor | B. Kinase inhibitor |
| **Vismodegib** | Smoothened (Hh) inhibitor | X. Other |
| **Pentostatin** | Antimetabolite; Purine analog | A. Conv. Chemo |
| **Estramustine aq** | Alkylating agent | A. Conv. Chemo |
| **Streptozocin** | Alkylating glucosamine-nitrosourea agent | A. Conv. Chemo |
| **Uracil mustard** | Alkylating agent | A. Conv. Chemo |
| **Arsenic(III) oxide aq** | Thioredoxin reductase inhibitor; cytotoxic chemotherapeutic | E. Differentiating/ epigenetic modifier |
| **APR-246** | p53 activator, thioredoxin reductase 1 inhibitor | G. Apoptotic modulator |
| **Bendamustine** | Nitrogen mustard alkylating agent | A. Conv. Chemo |
| **XAV-939** | Tankyrase-1 and -2 | E. Differentiating/ epigenetic modifier |
| **Lasofoxifene** | Selective estrogen receptor modulator | F. Hormone therapy |
| **Galunisertib** | TGF-B/Smad inhibitor | B. Kinase inhibitor |
| **MK-0752** | gamma-secretase/notch inhibitor | X. Other |
| **Ibrutinib** | Btk inhibitor | B. Kinase inhibitor |
| **NVP-BGJ398** | FGFR inhibitor | B. Kinase inhibitor |
| **Varespladib** | Secretory phospholipase A2 inhibitor | X. Other |
| **15D-PGJ2** | Endogenous PPARγ ligand, prostaglandin, NFkB signaling inhibitor | X. Other |
| **RD162** | AR antagonist | F. Hormone therapy |
| **Enzalutamide** | AR antagonist | F. Hormone therapy |
| **CPI-613** | pyruvate dehydrogenase, alpha-ketoglutarate dehydrogenase inhibitor | H. Metabolic modifier |
| **Rebastinib** | Allosteric ABL, FLT3, TIE2, TRKA inhibitor | B. Kinase inhibitor |
| **Venetoclax** | Selective Bcl-2 inhibitor | G. Apoptotic modulator |
| **UNC1215** | L3MBTL3 inhibitor | E. Differentiating/ epigenetic modifier |
| **Alectinib** | ALK (incl gatekeeper mut) inhibitor | B. Kinase inhibitor |
| **GSK-1838705A** | IGF1R, INSR, ALK inhibitor | B. Kinase inhibitor |
| **TRAM-34** | intermediate-conductance Ca2+-activated K+ channel inhibitor | X. Other |
| **Baricitinib** | JAK inhibitor | B. Kinase inhibitor |
| **Orteronel** | CYP17A1, androgen synth inhibitor | F. Hormone therapy |
| **EPZ-5687** | EZH2 inhibitor | E. Differentiating/ epigenetic modifier |
| **ARN 509** | AR antagonist | F. Hormone therapy |
| **Tideglusib** | GSK3 inhibitor | B. Kinase inhibitor |
| **Cilengitide** | alphaVbeta3 integrin inhibitor | X. Other |
| **Varlitinib** | EGFR HER2 inhibitor | B. Kinase inhibitor |
| **Sapitinib** | Pan-HER inhibitor | B. Kinase inhibitor |
| **AGI-5198** | IDH1 R132H/R132C inhibitor | H. Metabolic modifier |
| **VX 745** | p38MAPK inhibitor | B. Kinase inhibitor |
| **TAK-960** | PLK1 inhibitor | B. Kinase inhibitor |
| **PH-797804** | p38MAPK inhibitor | B. Kinase inhibitor |
| **AMG-208** | MET inhibitor | B. Kinase inhibitor |
| **NVP-INC280** | MET inhibitor | B. Kinase inhibitor |
| **Palomid-529** | AKT, MTOR, PI3K inhibitor | B. Kinase inhibitor |
| **TAK-285** | HER2 inhibitor | B. Kinase inhibitor |
| **SGX-523** | MET inhibitor | B. Kinase inhibitor |
| **JNJ-38877605** | MET inhibitor | B. Kinase inhibitor |
| **PF-04217903** | MET inhibitor | B. Kinase inhibitor |
| **EPZ-5676** | DOT1L inhibitor | E. Differentiating/ epigenetic modifier |
| **IRAK1/4 inhibitor** | IRAK1/4 inhibitor | B. Kinase inhibitor |
| **GSK2801** | BAZ2B/A bromodomain inhibitor | E. Differentiating/ epigenetic modifier |
| **VGX-1027** | Immunomodulator | D. Immunomodulatory |
| **Encorafenib** | B-RAF(V600E) | B. Kinase inhibitor |
| **URB597** | FAAH inhibitor | H. Metabolic modifier |
| **NLG919** | IDO inhibitor | D. Immunomodulatory |
| **VE-821** | ATR inhibitor | B. Kinase inhibitor |
| **Tasquinimod** | S100A9, immunomodulatory, anti-angiogenic | D. Immunomodulatory |
| **WEHI-539** | Bcl-XL inhibitor | G. Apoptotic modulator |
| **Gefitinib** | EGFR inhibitor | B. Kinase inhibitor |
| **Exemestane** | Aromatase inhibitor | F. Hormone therapy |
| **Silmitasertib** | CSNK2A1 inhibitor | B. Kinase inhibitor |
| **(+)JQ1** | BET family inhibitor | E. Differentiating/ epigenetic modifier |
| **Ralimetinib** | p38MAPK inhibitor | B. Kinase inhibitor |
| **Omipalisib** | PI3K/mTOR inhibitor | B. Kinase inhibitor |
| **AZ-23** | Trk inhibitor | B. Kinase inhibitor |
| **Dacomitinib** | pan-HER inhibitor | B. Kinase inhibitor |
| **Icotinib** | EGFR inhibitor | B. Kinase inhibitor |
| **VER 155008** | HSP70 inhibitor | K. HSP inhibitor |
| **Crenolanib** | PDGFRA and PDGFRB inhibitor | B. Kinase inhibitor |
| **Bortezomib** | Proteasome inhibitor (26S subunit) | A. Conv. Chemo |
| **Momelotinib** | JAK1 & 2 inhibitor | B. Kinase inhibitor |
| **Danusertib** | Aurora, Ret, TrkA, FGFR-1 inhibitor | B. Kinase inhibitor |
| **Dabrafenib** | B-Raf(V600E) inhibitor | B. Kinase inhibitor |
| **GSK923295** | CENP-E inhibitor | I. Kinesin inhibitor |
| **OSI-930** | KIT, VEGFR inhibitor | B. Kinase inhibitor |
| **deltarasin** | Ras-PDEdelta inhibitor | X. Other |
| **GSK269962** | ROCK1 and ROCK2 inhibitor | B. Kinase inhibitor |
| **PF-4800567** | CK1epsilon inhibitor | B. Kinase inhibitor |
| **TH588** | MTH1 inhibitor | H. Metabolic modifier |
| **Hydroxyfasudil** | ROCK inhibitor | B. Kinase inhibitor |
| **Tubacin** | HDAC6 inhibitor | E. Differentiating/ epigenetic modifier |
| **SGI-1776** | PIM kinase inhibitor | B. Kinase inhibitor |
| **AZD-1080** | GSK3 inhibitor | B. Kinase inhibitor |
| **Brivanib** | VEGFR inhibitor | B. Kinase inhibitor |
| **TAK-715** | p38MAPK inhibitor | B. Kinase inhibitor |
| **Tanespimycin** | HSP90 inhibitor | K. HSP inhibitor |
| **Thio-TEPA** | Alkylating agent | A. Conv. Chemo |
| **1-methyl-D-tryptophan aq** | Indolamine 2,3-dioxygenase 1 and 2 inhibitor | X. Other |
| **Capecitabine** | 5-FU prodrug | A. Conv. Chemo |
| **AGI-6780** | IDH2-R140Q inhibitor | H. Metabolic modifier |
| **Bosutinib** | Abl, Src inhibitor | B. Kinase inhibitor |
| **Tacrolimus** | Binds FKBP12, causes inhibition of calcineurin | C. Rapalog |
| **SGC0946** | DOT1L inhibitor | E. Differentiating/ epigenetic modifier |
| **XL019** | JAK2 inhibitor | B. Kinase inhibitor |
| **PF-04708671** | p70S6K inhibitor | B. Kinase inhibitor |
| **Ganetespib** | HSP90 inhibitor | K. HSP inhibitor |
| **CP-724714** | EGFR ERBB2 inhibitor | B. Kinase inhibitor |
| **Toremifene** | selective estrogen receptor modulator | F. Hormone therapy |
| **Alisertib** | Aurora A inhibitor | B. Kinase inhibitor |
| **IOX-1** | 2-Oxoglutarate Oxygenase Inhibitor | E. Differentiating/ epigenetic modifier |
| **Linsitinib** | IGF1R, IR inhibitor | B. Kinase inhibitor |
| **Carboplatin aq** | Platinum-based antineoplastic agent | A. Conv. Chemo |
| **Bentamapimod** | JNK inhibitor | B. Kinase inhibitor |
| **Erlotinib** | EGFR inhibitor | B. Kinase inhibitor |
| **Amuvatinib** | Broad spectrum TK inhibitor | B. Kinase inhibitor |
| **AZ191** | DYRK1A inhibitor | B. Kinase inhibitor |
| **BIIB021** | HSP90 inhibitor | K. HSP inhibitor |
| **AZD7762** | Chk1 inhibitor | B. Kinase inhibitor |
| **Lovastatin** | HMG-CoA reductase inhibitor | H. Metabolic modifier |
| **Ruxolitinib** | JAK1&2 inhibitor | B. Kinase inhibitor |
| **BGB324** | Axl inhibitor | B. Kinase inhibitor |
| **NMS-873** | p97/VCP inhibitor | X. Other |
| **Alvespimycin** | HSP90 inhibitor | K. HSP inhibitor |
| **Luminespib** | HSP90 inhibitor | K. HSP inhibitor |
| **PF-3845** | FAAH inhibitor | X. Other |
| **AZ 3146** | Mps1 kinase (TTK) inhibitor | B. Kinase inhibitor |
| **Tozasertib** | pan-Aurora inhibitor | B. Kinase inhibitor |
| **Atorvastatin** | HMG CoA reductase inhibitor | H. Metabolic modifier |
| **MK-2461** | MET inhibitor | B. Kinase inhibitor |
| **Mercaptopurine** | Antimetabolite | A. Conv. Chemo |
| **Simvastatin** | HMG CoA reductase inhibitor | H. Metabolic modifier |
| **(5Z)-7-Oxozeaenol** | TAK1 inhibitor | B. Kinase inhibitor |
| **BMS-911543** | JAK2 inhibitor | B. Kinase inhibitor |
| **BMS-754807** | IGF1R inhibitor | B. Kinase inhibitor |
| **SGC-CBP30** | CREBBP/EP300 bromodomain inhibitor | E. Differentiating/ epigenetic modifier |
| **GSK-1904529A** | IGF1R, INSR inhibitor | B. Kinase inhibitor |
| **AZD1152-HQPA** | Aurora B inhibitor | B. Kinase inhibitor |
| **Oprozomib** | Irreversible proteasome (20 S) inhibitor | A. Conv. Chemo |
| **SNS-314** | AURa, AURb inhibitor | B. Kinase inhibitor |
| **Midostaurin** | PKC, PKA, S6K and EGFR inhibitor | B. Kinase inhibitor |
| **Orantinib** | KDR, FGFR, PDGFR inhibitor | B. Kinase inhibitor |
| **BMS-599626** | Pan-HER inhibitor | B. Kinase inhibitor |
| **AT9283** | Aurora A & B, Jak2, Flt, Abl inhibitor | B. Kinase inhibitor |
| **Tosedostat** | Aminopeptidase inhibitor | X. Other |
| **Vandetanib** | VEGFR,EGFR, RET inhibitor | B. Kinase inhibitor |
| **Disulfiram** | alcohol dehydrogenase inhibitor | H. Metabolic modifier |
| **Carfilzomib** | Proteasome inhibitor (20S subunit) | A. Conv. Chemo |
| **Megestrol** | Progestogen | F. Hormone therapy |
| **TAK-901** | Aurora B inhibitor | B. Kinase inhibitor |
| **NVP-AEW541** | IGF1R inhibitor | B. Kinase inhibitor |
| **ASP3026** | ALK inhibitor | B. Kinase inhibitor |
| **PF-03814735** | AURa, AURb inhibitor | B. Kinase inhibitor |
| **Pemetrexed aq** | Dihydrofolate reductase inhibitor | H. Metabolic modifier |
| **BMS-777607** | Met, Axl, Ron and Tyro3 inhibitor | B. Kinase inhibitor |
| **NVP-AEE788** | EGFR, VEGFR, ABL, SRC inhibitor | B. Kinase inhibitor |
| **TPCA-1** | IKK-2 inhibitor | B. Kinase inhibitor |
| **Fostamatinib aq** | Syk inhibitor | B. Kinase inhibitor |
| **Vemurafenib** | B-Raf(V600E) inhibitor | B. Kinase inhibitor |
| **Tamatinib** | Syk inhibitor | B. Kinase inhibitor |
| **SP600125** | pan-JNK inhibitor | B. Kinase inhibitor |
| **Doramapimod** | p38 inhibitor | B. Kinase inhibitor |
| **ONX-0914** | LMP7 (immunoproteasome) | X. Other |
| **LY-2874455** | FGFR inhibitor | B. Kinase inhibitor |
| **Mubritinib** | ERBB2 inhibitor | B. Kinase inhibitor |
| **Gandotinib** | JAK2 inhibitor | B. Kinase inhibitor |
| **CYC-116** | Aurora and VEGFR2 inhibitor | B. Kinase inhibitor |
| **Saracatinib** | Src, Abl inhibitor | B. Kinase inhibitor |
| **Dovitinib** | FGFR inhibitor | B. Kinase inhibitor |
| **RAF265** | C-Raf inhibitor | B. Kinase inhibitor |
| **Pacritinib** | FLT3/JAK2 | B. Kinase inhibitor |
| **SAR302503** | JAK2-selective inhibitor | B. Kinase inhibitor |
| **AZD4547** | FGFR inhibitor | B. Kinase inhibitor |
| **Prednisone** | Immunomodulatory agent | D. Immunomodulatory |
| **NVP-LGK974** | PORCN inhibitor | X. Other |
| **KW-2449** | AURa AURb FLT3 inhibitor | B. Kinase inhibitor |
| **YM155** | Survivin inhibitor | G. Apoptotic modulator |
| **Linifanib** | VEGFR, PDGFR, CSF-1R, FLT3 inhibitor | B. Kinase inhibitor |
| **Tandutinib** | FLT3, PDGFR, KIT inhibitor | B. Kinase inhibitor |
| **ENMD-2076** | pan-Aurora inhibitor | B. Kinase inhibitor |
| **Semaxanib** | VEGFR inhibitor | B. Kinase inhibitor |
| **Apatinib** | VEGFR inhibitor | B. Kinase inhibitor |
| **Sorafenib** | B-Raf, FGFR-1, VEGFR-2 & -3, PDGFR-beta, KIT, and FLT3 inhibitor | B. Kinase inhibitor |
| **Bafetinib** | Abl, Lyn inhibitor | B. Kinase inhibitor |
| **Cediranib** | KDR/Flt/VEGFR inhibitor | B. Kinase inhibitor |
| **Vatalanib** | VEGFR-1 & -2 inhibitor | B. Kinase inhibitor |
| **Regorafenib** | B-Raf, c-Kit, VEGFR2 inhibitor | B. Kinase inhibitor |
| **Motesanib** | VEGFR, PDGFR, Ret, Kit inhibitor | B. Kinase inhibitor |
| **Prednisolone** | Immunomodulatory agent | D. Immunomodulatory |
| **Ponatinib** | Broad TK inhibitor | B. Kinase inhibitor |
| **KRN-633** | VEGFR inhibitor | B. Kinase inhibitor |
| **CEP-32496** | BRAF inhibitor | B. Kinase inhibitor |
| **Methylprednisolone** | Immunosuppressant | D. Immunomodulatory |
| **Dexamethasone** | Immunosuppressant; glucocorticoid | D. Immunomodulatory |
| **Nintedanib** | VEGFR, PDGFR, FGFR inhibitor | B. Kinase inhibitor |
| **Telatinib** | VEGFR, KIT, PDGFR inhibitor | B. Kinase inhibitor |
| **Quizartinib** | FLT3 inhibitor | B. Kinase inhibitor |
| **Foretinib** | MET, VEGFR2 inhibitor | B. Kinase inhibitor |
| **Dasatinib** | BCR/Abl, Src, Kit, EphR ... Inhibitor | B. Kinase inhibitor |
| **Sunitinib** | Broad TK inhibitor | B. Kinase inhibitor |
| **Golvatinib** | MET, VEGFR2 inhibitor | B. Kinase inhibitor |
| **Nilotinib** | BCR/Abl inhibitor | B. Kinase inhibitor |
| **Imatinib** | Abl, Kit, PDGFRB inhibitor | B. Kinase inhibitor |
| **Masitinib** | KIT inhibitor | B. Kinase inhibitor |
| **Cabozantinib** | VEGFR2, Met, FLT3, Tie2, Kit and Ret inhibitor | B. Kinase inhibitor |
| **Lenvatinib** | VEGFR inhibitor | B. Kinase inhibitor |
| **Pazopanib** | VEGFR inhibitor | B. Kinase inhibitor |
| **Lucitanib** | FGFR1, VEGFR inhibitor | B. Kinase inhibitor |
| **Axitinib** | VEGFR, PDGFR, KIT inhibitor | B. Kinase inhibitor |
| **Tivozanib** | VEGFR1, 2, 3, c-Kit, PDGFRB inhibitor | B. Kinase inhibitor |

Name, molecular targets and classification of 459 compounds used in DSRT (Drug Sensitivity and Resistance Testing) using HMC-1 cell lines.

**Supplemental Table 9.** **Nintedanib effect on primary SM patient samples.**

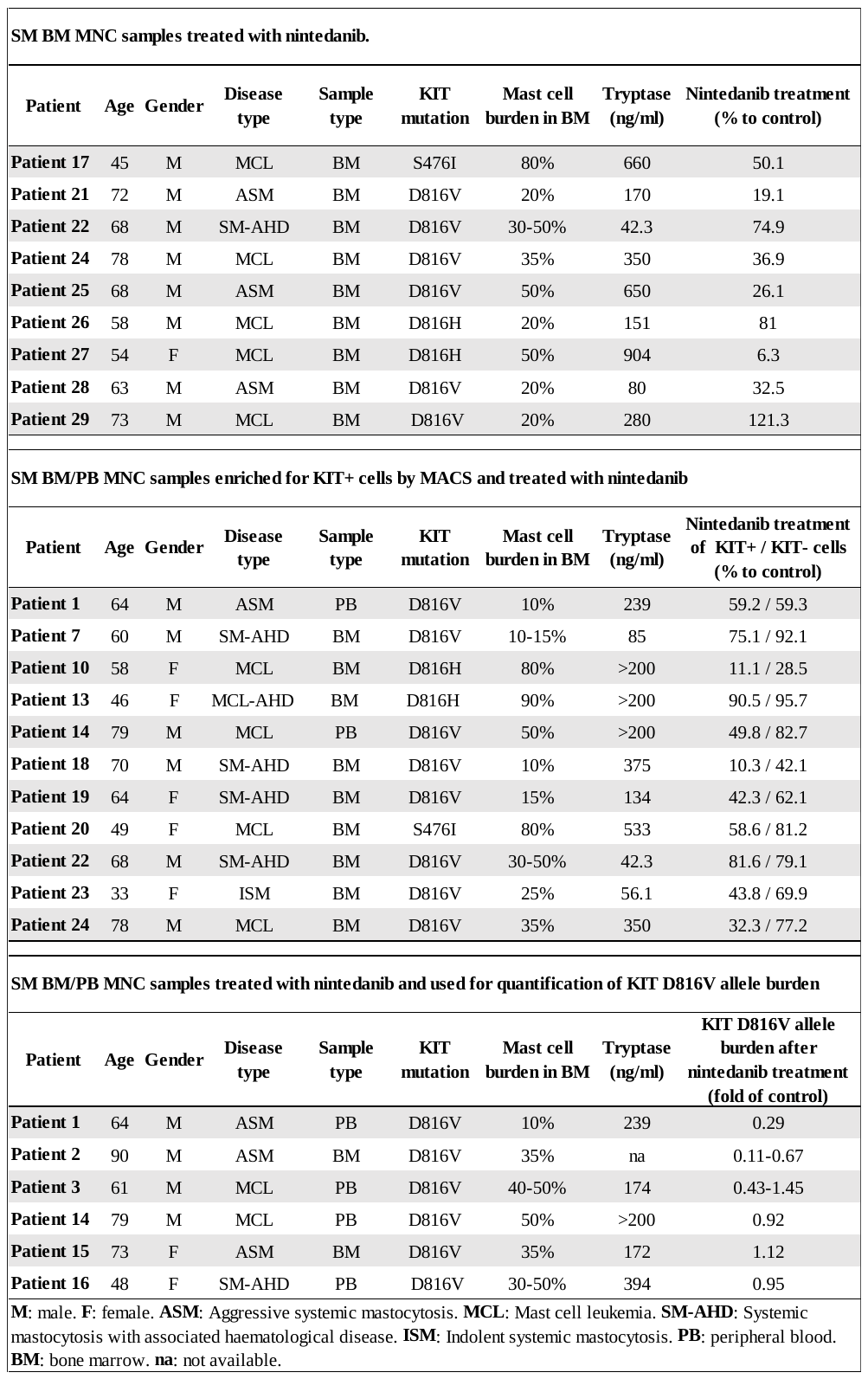

**Supplemental Table 10.** **Similarity index of 43 nintedanib analogues tested.**

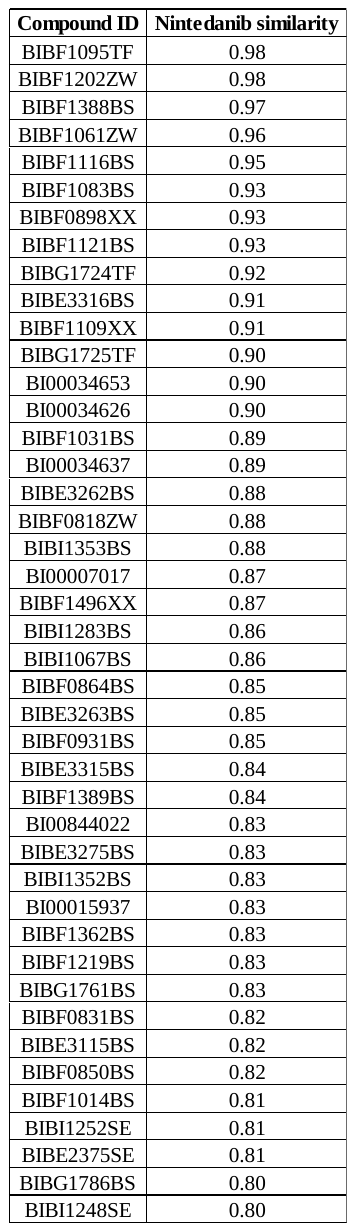

The structural similarity of each compound to nintedanib is represented as a similarity index with the maximum value set as 1. All 43 compounds used in the present work have a similarity index ranging from 0.8 up to 0.98.

**Supplemental Table 11.** **Oligonucleotides for RT-PCR analysis.**
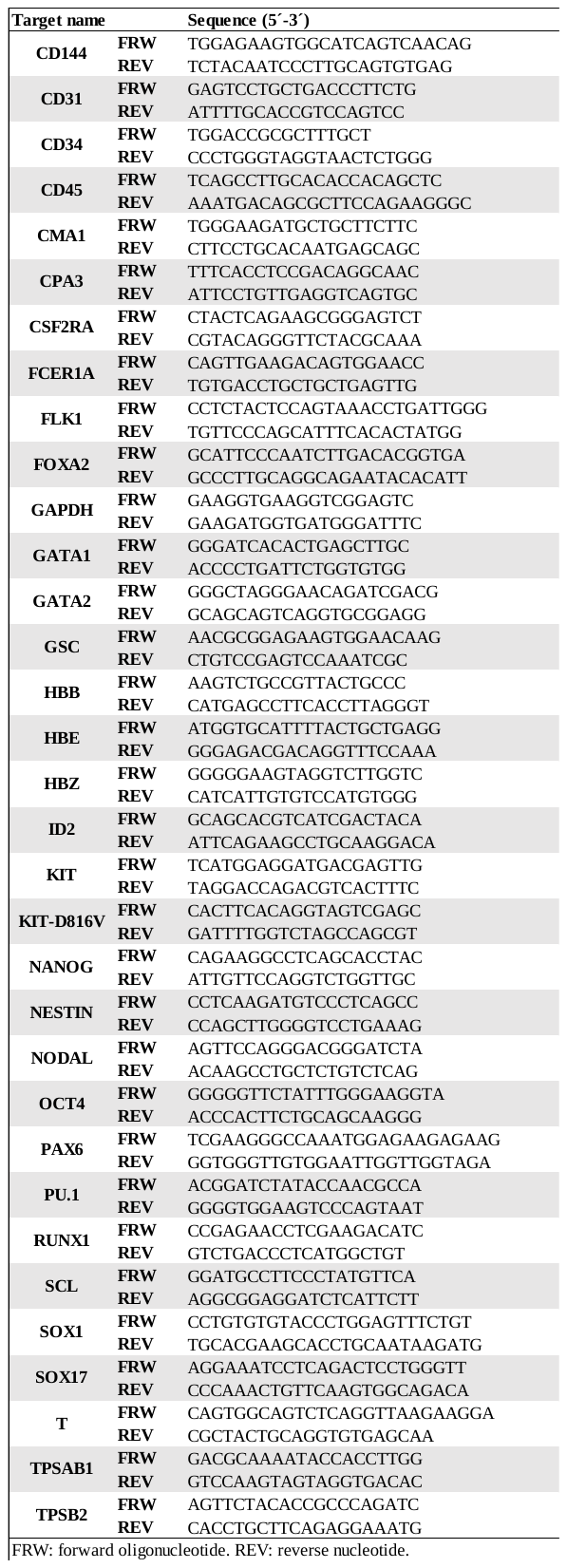

**Supplemental Figure Legends**

**Supplemental Figure 1. Characterization of patient-specific iPSCs.**

(A) *KIT* D816V and control iPSCs express the pluripotency markers OCT4, NANOG, TRA-1-60 and TRA-1-81 as assessed by immunofluorescence. Nuclei were stained with DAPI (blue). Scale bar 100 μm. Images are representative for all iPSCs generated in the study. (B) Pluripotency assessment of iPSCs using Epi-Pluri-Score analysis. Blue area represents DNA methylation profiles of 1951 non-pluripotent cell samples and red area represents DNA methylation profiles of 264 pluripotent samples profiles (all Illumina HumanMethylation27 BeadChip platform). All *KIT* D816V iPSCs and control iPSCs (red and blue dots, respectively) generated in the present work (patient 1-3, P1-P3) revealed a positive Epi-Pluri-Score and are therefore classified as pluripotent. (C) Karyotype analysis of *KIT* D816V iPSC of patient 1. The image shown is representative for the karyotype of all iPSCs generated in the present work and no chromosomal aberrations were identified. (D) Analysis of KIT expression on the surface of iPSCs by flow cytometry (top) reveals that *KIT* D816V iPSCs have lower amount of cell surface KIT receptor in comparison to control iPSCs. Bottom: RT-qPCR analysis of *KIT* mRNA expression in *KIT* D816V iPSC (red, n=8) and control iPSCs (blue, n=5). No statistically significant difference was found (p=0.40; ns, not significant). (E) The amount of KIT receptor on the surface of mutated and control iPSCs is reduced upon SCF stimulation (open curves, without SCF; filled curves with SCF). Bottom: KIT receptor mean fluorescence intensity (MFI, n=4 for each condition) on *KIT* D816V and control iPSCs before and after SCF stimulation (250 ng/ml for 15 min). *: p<0.05. **: p=0.0002. (F) Western blot analysis of KIT signaling in iPSCs upon SCF stimulation. Left: Representative Western blot analysis of KIT downstream signaling in *KIT* D816V and control iPSCs (patient 1) before (-) and after (+) SCF stimulation (250 ng/ml for 15 min). Position of molecular weight marker is indicated on the left in kDa. Right: Quantification of glycosylated and not glycosylated phospho-KIT bands is shown as ratio of SCF stimulated vs not stimulated (+SCF/-SCF ratio, n=3-4). *: p<0.05. All the statistical analyses were performed with Welch´s t-test.

**Supplemental Figure 2. Impact of *KIT* D816V on iPSC-derived hematopoietic cells.**

(A) CD45^+^/KIT^high^ MCs appear earlier in *KIT* D816V hematopoiesis than in unmutated control. Kinetics of *KIT* D816V iPSC differentiation (patient 1) shows emerging CD45^+^/KIT^high^ MCs/MC progenitors at early time points compared to unmutated control. The flow cytometry analysis is representative for all iPSC clones of patient 1. (B) *KIT* D816V mutation confers a proliferative advantage to KIT^+^ cells assessed by MTT assay. Mutated KIT^+^ hematopoietic cells have a proliferative advantage in comparison to unmutated cells regardless of the presence of SCF (top and middle plots). Upon complete cytokine withdrawal, mutated cells demonstrated sustained cell survival, which was not observed for control cells. *KIT* D816V and control KIT^-^ cells showed similar proliferation under the conditions tested. P1: patient 1. P2: patient 2. Statistical analysis was performed for 96 h values using Welch´s t-test (n=3). *: p<0.01, **: p<0.001.

**Supplemental Figure 3. Characterization of hematopoietic cells differentiated from *KIT* D816V and control iPSCs.**

(A) Surface marker phenotype of *KIT* D816V iPSC-derived erythroid cells (patient 1) by flow cytometry. CD235α^+^ *KIT* D816V cells also express CD43 and KIT. CD45 is expressed on a minor fraction of the CD235α^+^ population. Stained sample is shown as red line. Isotype control is showed as gray filled curve. (B) Flow cytometry characterization of *KIT* D816V and control iPSCs-derived hematopoietic cells. Box plots for each population analyzed are shown. **P1**: Box plots for patient 1 show the averaged values of eight independent differentiation experiments of three *KIT* D816V iPSCs (red) analyzed between 13 and 49 days of hematopoietic differentiation (n=30) and six independent differentiation experiments of two control iPSCs (blue) analyzed between 15 and 49 days of hematopoietic differentiation (n=19). *KIT* D816V hematopoietic cell populations (except for CD31/CD34) show statistically significant differences in comparison to control populations (*=p ≤ 0.007). **P2**: Box plots for patient 2 show the averaged values of three independent differentiation experiments of one *KIT* D816V iPSC (red) analyzed between 16 and 48 days of hematopoietic differentiation (n=12) and one independent differentiation experiment of one control iPSC (blue) analyzed between 29 and 59 days of hematopoietic differentiation (n=4). *KIT* D816V and control hematopoietic cell populations show no statistically significant differences. **P3**: Box plots for patient 3 show the averaged values of three independent differentiation experiments of one *KIT* D816V iPSC (red) analyzed between 15 and 45 days of hematopoietic differentiation (n=7) and three independent differentiation experiments of two control iPSCs (blue) analyzed between 17 and 44 days of hematopoietic differentiation (n=14). *KIT* D816V and control hematopoietic cell populations show no statistically significant differences. Statistical analysis was performed with Welch´s t-test. (C) Flow cytometry cell sorting experiment for patient 1 *KIT* D816V iPSC-derived hematopoietic cells (left panel). Cytospin preparations of sorted CD235α^+^/KIT^+/-^ cells show different stages of erythroblast maturation. Basophilic (red arrow), polychromatic (black arrowhead) and orthochromatic (black arrow) erythroblasts were observed. Scale bars: 50 μm.

**Supplemental Figure 4. Impaired hematopoietic differentiation of patient 2 iPSCs.**

(A) Patient 2 iPSC-derived hematopoietic cells show a myeloid biased phenotype. Cytospins of *KIT* D816V and control iPSC-derived cells exhibit an immature myeloid (myeloblast-like) phenotype (green arrowheads) in agreement with the prominent CD45^+^/KIT^+^ population observed by flow cytometry (Figure 1, C and E). The *KIT* D816V sample also shows some nucleated erythrocytes (normoblasts, red arrowheads) and erythrocytes precursors (erythroblasts, red arrows). Scale bars: 50 μm. (B) Annexin V/PI staining of iPSC-derived hematopoietic cells. Patient 2 *KIT* D816V and control iPSCs at day 10-20 of hematopoietic differentiation (n=3-6) show prominent apoptosis in comparison to patients 1 and 3 (left). Representative annexin V/PI flow cytometry plots for iPSC-derived hematopoietic cells, showing prominent Annexin+/PI+ population (quadrant III) in patient 2 sample (right). *: p≤0.0001. Statistical analysis was performed with Welch´s t-test.

**Supplemental Figure 5. CD45^+^/KIT^high^ cells in patient 3 iPSC-derived hematopoietic cells.**

(A) Kinetics of *KIT* D816V and control iPSC differentiation shows CD45^+^/KIT^high^ MCs in both genotypes. (B) Representative images of cytospin preparations of *KIT* D816V iPSC-derived hematopoietic cells. Macrophages (red arrowheads), immature myeloid (myeloblast-like, green arrowheads) and more differentiated myeloid cells (green arrows) are shown. Scale bars: 50 μm.

**Supplemental Figure 6. NGS analysis of primary patient samples and of iPSCs derived thereof shows patient specific mutation profiles.**

(A) Mutations identified by NGS in primary PB or BM samples of patients 1-3 and in *KIT* D816V and control iPSCs generated. P.S.: primary sample. 1-3: *KIT* D816V or control iPSCs generated for each patient. (B) Allele burden of mutations identified in primary samples of patient 1-3. Please note that for some genes, more than one mutation was identified (supplemental Table 5-7) and these are also represented.

**Supplemental Figure 7. Introduction of the *KIT* D816V mutation into ESCs by CRISPR/Cas9n.**

(A) Schematic representation of CRISPR/Cas9n mediated introduction of the *KIT* D816V mutation into HES-3 ESCs (ES03). Two gRNAs were designed for targeting exon 17 of the KIT gene, generating a double strand break (DSB)*.* Expected nick sites are indicated by yellow arrowheads and codon 816 is in bold letters. A 200 nucleotide single stranded oligonucleotide (ssODN) was used as donor template harboring the A-T nucleotide change in codon 816 plus a silent mutation (C-T) in codon 814 to disrupt the PAM sequence. (B) Allele-specific PCR identifies ESCs with *KIT* D816V introduced by CRISPR/Cas9n. HMC1.1 and HMC1.2 MC lines without and with *KIT* D816V mutation, respectively, were used as negative (-) and positive (+) controls. *KIT* editing was observed in 27.5% (66/240) of the screened clones of which homology directed repair (HDR) was observed in 13.6% (9/66). Heterozygous *KIT* D816V was observed in 6% (4/66) of the edited clones. Two KIT D816V ESC heterozygous cell lines were selected for further studies. W: water control. M: molecular weight marker. (C) Sanger sequencing shows heterozygous *KIT* D816V mutation in ESCs. Clone D816V 1 shows only the A-T substitution in codon 816 (D816V mutation) and clone D816V 2 shows the D816V mutation and additionally the C-T silent mutation in codon 814. (D) ESC pluripotency was not compromised by the introduction of the KIT D816V mutation (in both KIT D816V ESC lines evaluated) as demonstrated by Epi-Pluri-Score analysis. *KIT* D816V and control ESCs showed a positive Epi-Pluri-Score and are thus classified as pluripotent. (E) *KIT* D816V 1 and 2 ESCs express the pluripotency markers OCT4, NANOG, TRA-1-60 and TRA-1-81. Immunofluorescence analysis was as in supplemental Figure 1A. Scale bar: 100 μm.

**Supplemental Figure 8. Characterization of *KIT* D816V ESCs.**

(A) Representative phase contrast image of *KIT* D816V ESCs on MEF layer with typical pluripotent morphology (top left). Scale bar: 500 μm. Representative flow cytometry histogram plots for KIT receptor on *KIT* D816V and control ESCs (red and blue, respectively) show reduced expression of surface KIT receptor on mutated cells (top right). RT-qPCR analysis of *KIT* mRNA expression in *KIT* D816V ESCs (red, n=3) and control ESCs (blue, n=3) (bottom). No statistically significant difference was found (p=0.68; ns, not significant). (B) Representative Western blot analysis of KIT signaling before (-) and after (+) SCF stimulation (250 ng/ml for 15 min) in *KIT* D816V 1 and 2 ESCs and control ESCs (left). Size marker as in supplemental Figure 1F. Quantification of glycosylated and not glycosylated phospho-KIT bands is shown (n=3-6) as SCF stimulated / SCF not stimulated ratios (right). *: p≤0.05. (C) Flow cytometry histogram plots for KIT receptor on the surface of *KIT* D816V 1 and 2 ESCs (red) and control ESCs (blue) before and after SCF stimulation (open and filled curves, respectively) shows reduction in cell surface KIT expression upon SCF stimulation (250 ng/ml for 15 min). KIT receptor MFI on *KIT* D816V and control ESCs upon SCF stimulation (right). *: n=4, p=0.01. **: n=6, p=0.0002. Statistical analysis was performed with Welch´s t-test.

**Supplemental Figure 9. Hematopoietic differentiation of *KIT* D816V ESCs.**

Quantification of hematopoietic cell populations derived from *KIT* D816V ESCs 1 and 2 and from unmutated control. No significant differences between KIT D816V and unmutated cells were observed for hematopoietic cells (CD45^+^) or granulocyte (CD45+/CD66b+) populations. Statistical analysis was performed using Welch´s t-test by comparing *KIT* D816V and control cell populations. *: n=4, p≤0.05; **: n=4, p=0.001.

**Supplemental Figure 10. Compound screening on HMC-1 cell lines.**

(A) Screening of 459 compounds on HMC-1.2 (*KIT* D816V/V560G) and control HMC-1.1 (*KIT* V560G) cell lines. The relative inhibition upon drug treatment is shown for HMC-1.1 and HMC-1.2. Drug responses for nintedanib, midostaurin, imatinib and rigosertib are highlighted. (B) Drug response curves (0 to 10 μM, n=3) of HMC-1.2 (red) and HMC-1.1 (blue) cell lines treated with nintedanib (IC50_HMC-1.2_: 585 nM, IC50_HMC-1.1_: 4 nM), midostaurin (IC50_HMC-1.2_: 448 nM, IC50_HMC-1.1_: 321 nM), imatinib (IC50_HMC-1.2_: >100 µM, IC50_HMC-1.1_: 34 nM) and rigosertib (IC50_HMC-1.2_: 230 nM, IC50_HMC-1.1_: 101 nM) for 66 h.

**Supplemental Figure 11. Nintedanib targets KIT^+^ D816V iPSC and ESC-derived hematopoietic cells.**

(A) Drug response analysis of KIT^+^ and KIT^-^ hematopoietic cells derived from *KIT* D816V and control iPSCs of patients 1-3. Cells were treated with 1μM of midostaurin, nintedanib, imatinib or rigosertib for 66 h and subjected to CellTiter Glo assay (n=3-12). Cell viability reduction is more prominent on KIT^+^ cells upon nintedanib or midostaurin treatment. Imatinib treatment shows less activity on KIT^+^ and KIT^-^ cells compared to nintedanib and midostaurin while rigosertib targets both mutated and unmutated KIT^+^ cells. *: Welch´s t-test, p<0.05; **: Welch´s t-test, p<0.01; ***: Welch´s t-test, p<0.0001. (B) Drug response curves (0 to 10 µM) for nintedanib, midostaurin and imatinib on *KIT* D816V 1 and 2 ESC-derived KIT^+^ hematopoietic cells. Control: hematopoietic cells of unmutated ESCs. Each curve represents an independent experiment performed in triplicates. Nintedanib IC50 values were 302 and 2430 nM for *KIT* D816V and control cells, respectively. Midostaurin IC50 values were 200 and 3695 nM for *KIT* D816V and control cells, respectively. (C) Averaged drug response ± SD of *KIT* D816V 1, *KIT* D816V 2 and control ESC-derived KIT^+^ cells treated with 100 nM or 1 μM of nintedanib, midostaurin, imatinib or rigosertib for 66 h (n=9). Vehicle (DMSO) treated cells were used as control (0 nM). Statistical analysis was performed by Welch´s t-test comparing the drug responses of *KIT* D816V and control KIT^+^ cells at the same drug concentration. *: p≤0.05, **: p<0.001, ***: p<0.0001. (D) Drug response analysis of KIT^+^ and KIT^-^ hematopoietic cells derived from *KIT* D816V 1 and 2 ESCs and control ESCs. Cells were treated with 1μM of midostaurin, nintedanib, imatinib or rigosertib for 66 h as in (A) (n=9). Similar to iPSC-derived cells, nintedanib and midostaurin effects are more pronounced on KIT^+^ D816V cells. *: Welch´s t-test, p≤0.05; **: Welch´s t-test, p≤0.0001.

**Supplemental Figure 12. Nintedanib blocks KIT D816V signaling.**

Quantification of Western blot bands obtained for *KIT* D816V iPSC-derived hematopoietic cells treated for 4 h with 1 µM of nintedanib or midostaurin as shown in Figure 3C. Vehicle (DMSO) treated cells were used as control. At least 6 independent experiments were quantified. Compound treatment strongly reduced KIT receptor and STAT3 phosphorylation in comparison to control. For nintedanib a significant reduction of total KIT protein was also observed. Statistical analysis was performed with Welch´s t-test. *: p≤0.0033; **: p≤ 0.0004.

**Supplemental Figure 13. Nintedanib targets KIT^+^ cells derived from CD34^+^ HPCs.**

(A) Schematic representation of compound testing on iPSC-derived hematopoietic cells. *KIT* D816V and control iPSCs were differentiated into hematopoietic cells. Suspension cells were enriched for KIT^+^ cells by MACS and subjected to compound testing as in supplemental Figure 11. Alternatively, the adherent layer containing hemogenic endothelium was harvested and enriched for CD34^+^ HPCs by MACS. Those cells were expanded for 10-20 days followed by enrichment for KIT^+^ cells by MACS and used for compound testing. (B) Drug response curves for nintedanib, midostaurin and imatinib on patient 1 and patient 3 iPSC-derived HPCs. *KIT* D816V and control KIT^+^ cells were treated with compounds for 66 h before analysis. IC50 values for nintedanib were 84-95 nM and 807-895 nM for *KIT* D816V and control cells, respectively. For midostaurin, IC50 values were 45-63 nM and 312-808 nM for *KIT* D816V and control cells, respectively. (C) Drug response of patient 1 and patient 3 iPSC-derived *KIT* D816V and control HPCs treated with 100 nM or 1 µM nintedanib, midostaurin or imatinib. Significant reduction in viability of mutated cells, in comparison to control cells, was observed after nintedanib and midostaurin treatment. Bars represent the average of 2 independent experiments in triplicates (n=6). *: p<0.05; **: p<0.0001. (D) Representative flow cytometry dot plots of *KIT* D816V iPSC-derived CD34^+^ HPCs of patients 1 - 3 after 10, 15 and 20 days. The biased hematopoiesis, observed for suspension cells upon hematopoietic differentiation (Figure 1, C-H), was recapitulated in expanded CD34+ HPCs: erythroid bias in patient 1 cells, prominent CD45^+^/KIT^+^ population and apoptosis in patient 2 cells and prominent CD45^+^/KIT^high^ MCs in patient 3 cells. Statistical analysis was performed with Welch´s t-test.

**Supplemental Figure 14. Activity of avapritinib (BLU-285) and ripretinib (DCC-2618) on HMC-1.1 and HMC-1.2 cell lines.**

(A) Drug response curves (0 to 10 μM) for avapritinib (BLU-2815) and ripretinib (DCC-2618) on HMC-1.1 and HMC-1.2 cell lines (n=3) assessed by CellTiter Glo assay. IC50 values for avapritinib were 172 and 244 nM for HMC-1.1 and HMC-1.2 cells respectively. IC50 values for ripretinib were 4 and 120 nM for HMC-1.1 and HMC-1.2 cells respectively. (B) Western blot analysis of HMC-1.1 and HMC-1.2 cell lines treated with 1µM nintedanib, midostaurin, ripretinib or avapritinib for 4 h. Vehicle (DMSO) treated cells were used as control. (C) Quantification of pY-KIT, pS-AKT and pTY-ERK1/2 Western blot bands from 3 independent experiments. Avapritinib and ripretinib treatment strongly reduces KIT, STAT5, AKT and ERK1/2 phosphorylation in both cell lines. Statistical analysis was performed with Welch´s t-test (n=3). *: p≤0.05, **: p≤0.001.

**Supplemental Figure 15. Activity of avapritinib (BLU-285) and ripretinib (DCC-2618) on *KIT* D816V iPSC-derived hematopoietic cells.**

(A) Drug response curves (0 to 10 μM) for avapritinib (BLU-285) and ripretinib (DCC-2618) on *KIT* D816V and control iPSC-derived KIT^+^ hematopoietic cells (n=2 to 4) of patients 1 and 3. Avapritinib IC50 values were 31-56 nM and 1991-2585 nM for *KIT* D816V and control cells, respectively. Ripretinib IC50 values were 22-63 nM and 607-3870 nM for *KIT* D816V and control cells, respectively. (B) Averaged drug response ± SD of *KIT* D816V and control iPSC-derived KIT^+^ cells treated with 1 μM of nintedanib, avapritinib, ripretinib or midostaurin for 66 h. Vehicle (DMSO) treated cells were used as control. Statistical analysis was performed using Welch´s t-test (n=3-5) comparing the drug responses of *KIT* D816V and control KIT^+^ cells. *: p≤0.05, **: p≤0.001, ***: p≤0.0001.

**Supplementary Figure 16. Nintedanib decreases viability of SM primary samples.**

(A) Representative bar plots of BM MNCs from 4 SM patients harboring the *KIT* D816V mutation (patients 21, 22 and 24) or the *KIT* S476I mutation (patient 17) treated with 1 µM nintedanib or midostaurin for 66 h. Cell viability was measured with CellTiter Glo assay and normalized to the viability of DMSO treated cells. Strong response (≥50%) to nintedanib and midostaurin was observed in 3 SM samples. (B) Primary samples, where enough cells were available after MNC isolation, were evaluated for the response of KIT^+^ and KIT^-^ cells to nintedanib or midostaurin treatment. DMSO treated cells were used as control. For patients 1, 7, 14 and 20, cells were treated only with nintedanib due to limitation in cell numbers. For 6 SM primary samples (patients 10, 14, 18, 19, 23 and 24), a reduction in cell viability ≥ 50% was observed upon nintedanib treatment. For those samples, we also observed that nintedanib preferentially targeted KIT expressing cells. The observed variability of nintedanib response among samples might be due to differences in *KIT* D816V allele burden, cell composition and/or the genetic background including concurrent mutations. (C) The impact of nintedanib treatment on *KIT* D816V mutation burden was measured by allele-specific RT-qPCR. Primary MNCs were treated with nintedanib for 48 h and DMSO treated cells were used as control. Total *KIT* expression (left panel) indicates that *KIT* expressing cells comprised a small fraction of the primary sample. Nintedanib treatment caused a reduction in *KIT* D816V expressing cells (right panel), which was however not statistically significant. (D) Western blot analysis revealed efficient inhibition of KIT phosphorylation upon nintedanib treatment of *KIT* D816H SM primary cells from patient 10. Cells were prepared as described in (B) and treated with 1µM nintedanib for 4 h. DMSO treated cells were used as control. Nintedanib treatment led to pronounced reduction in KIT, STAT5 and ERK phosphorylation in KIT^+^ cells. As expected, KIT or phosphorylated KIT was not observed in KIT^-^ cells. Further information on the primary samples used in (A-D) are listed in Supplemental Table 9.

**Supplemental Figure 17. *KIT* D816V and control iPSC-derived MCs.**

(A) Bright field microscopy images of acidic Toluidine Blue stained cytospin preparations of sorted *KIT* D816V and control MCs. Metachromatic granules in the cytoplasm (red arrowheads) characteristic of MCs and cells lacking those granules (more immature MCs, red arrows) were observed. Scale bar: 50 μm. (B) Bright field microscopy images of tryptase stained cell smear preparations of sorted *KIT* D816V and control MCs. A homogenous population of cells with variable number of nucleus lobulations (orange arrowheads) and positive staining for tryptase were observed. Scale bar: 50 μm.

**Supplemental Figure 18. Molecular docking of nintedanib onto KIT D816V.**

(A) Root Main Square Deviation (RMSD) map of the pocket region belonging to seven kinases crystalized in the presence of ADP or inhibitor molecules as indicated. The values shown are expressed in angstrom (A) and they are re-weighted on the side chain symmetry and the secondary structures similarity. For each of the residues in the pocket, all the atoms are considered for the RMSD calculation. The RMSD of the atoms belonging to the KIT binding sites with respect VEGFR2 is below 2.23 A. (B) Glide Score of tautomers of ligands docked onto KIT D816V inactive and onto unmutated KIT in the inactive or active conformations. For nintedanib, both the isomeric states (cis/trans) at pH=7 were used and the cis-isomer displayed better binding poses than the trans-isomer. Therefore, we focused our analysis on the cis-isomer of nintedanib. Top: Averaged Glide Score obtained from different binding poses explored by each ligand. Number in parenthesis indicates the SD. *: cis conformation as in PDB code 3C7Q). **: trans conformation. Bottom: Bar plot showing the averaged Glide Scores (±SD) obtained by each ligand binding pose in the different conformations of unmutated or KIT D816V receptor. Low Glide Score values indicate higher binding affinity between ligand and receptor. Midostaurin and avapritinib show higher affinity to the active KIT conformation while ripretinib and imatinib docking studies suggest preference for the inactive conformations of KIT (unmutated or KIT D816V). Interestingly, ripretinib was recently co-crystalized with unmutated KIT^11^ Although a structure is not yet deposited, our model reproduces the reported ligand-receptor interaction: the hydrophobic interactions with L644, L656, and H790, as well as the HB with D810.

**Supplemental Figure 19. Molecular docking of nintedanib, midostaurin and imatinib onto inactive KIT D816V and unmutated KIT in the inactive and active conformations.**

Protein-ligand interaction details for nintedanib, midostaurin and imatinib are shown in 2D scheme. In the diagram, residues are represented as colored spheres, labelled with the residue name and residue number. Additional labels are shown in the figure.

**Supplemental Figure 20. Molecular docking of nintedanib analogues onto KIT D816V.**

(A) Glide Score of tautomers of ligands docked onto KIT D816V and unmutated KIT. Left: Averaged score obtained from different binding poses explored by each ligand. Number in parenthesis indicates the SD. *: cis conformation (as in PDB code 3C7Q).**: trans conformation. Right: Bar plot showing the average scores (±SD) obtained by each ligand binding pose. The average is computed over all the poses of all the tautomers for each molecule. (B) Protein-ligand interaction fingerprint (PLIF) generated with MOE (Molecular Operating Environment, 2013.08; Chemical Computing Group ULC, Montreal, Canada). Schematic representation of KIT receptor amino acids (columns, color coded) that mediate at least one protein-ligand interaction (black boxes) with each ligand (rows, up to 2 isomers each). Several interactions (e.g. salt bridges, p-stacking) may be formed between the KIT receptor amino acid residue and the ligand. Therefore, varying number of columns are observed for each residue. Compounds further analyzed in Figure 7 are indicated in red.

**Supplemental Figures****:**

**Supplemental Figure 1**
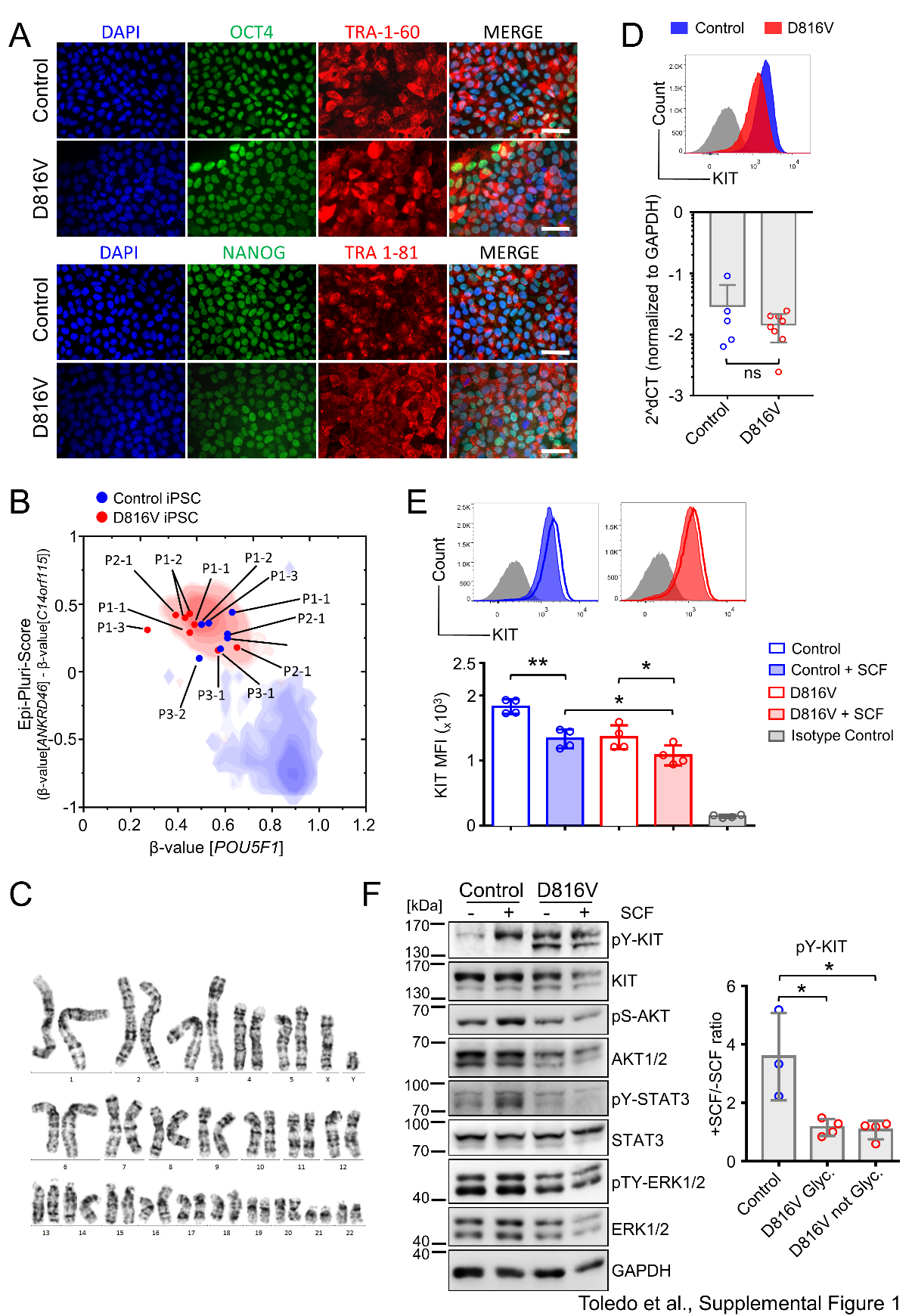

**Supplemental Figure 2**

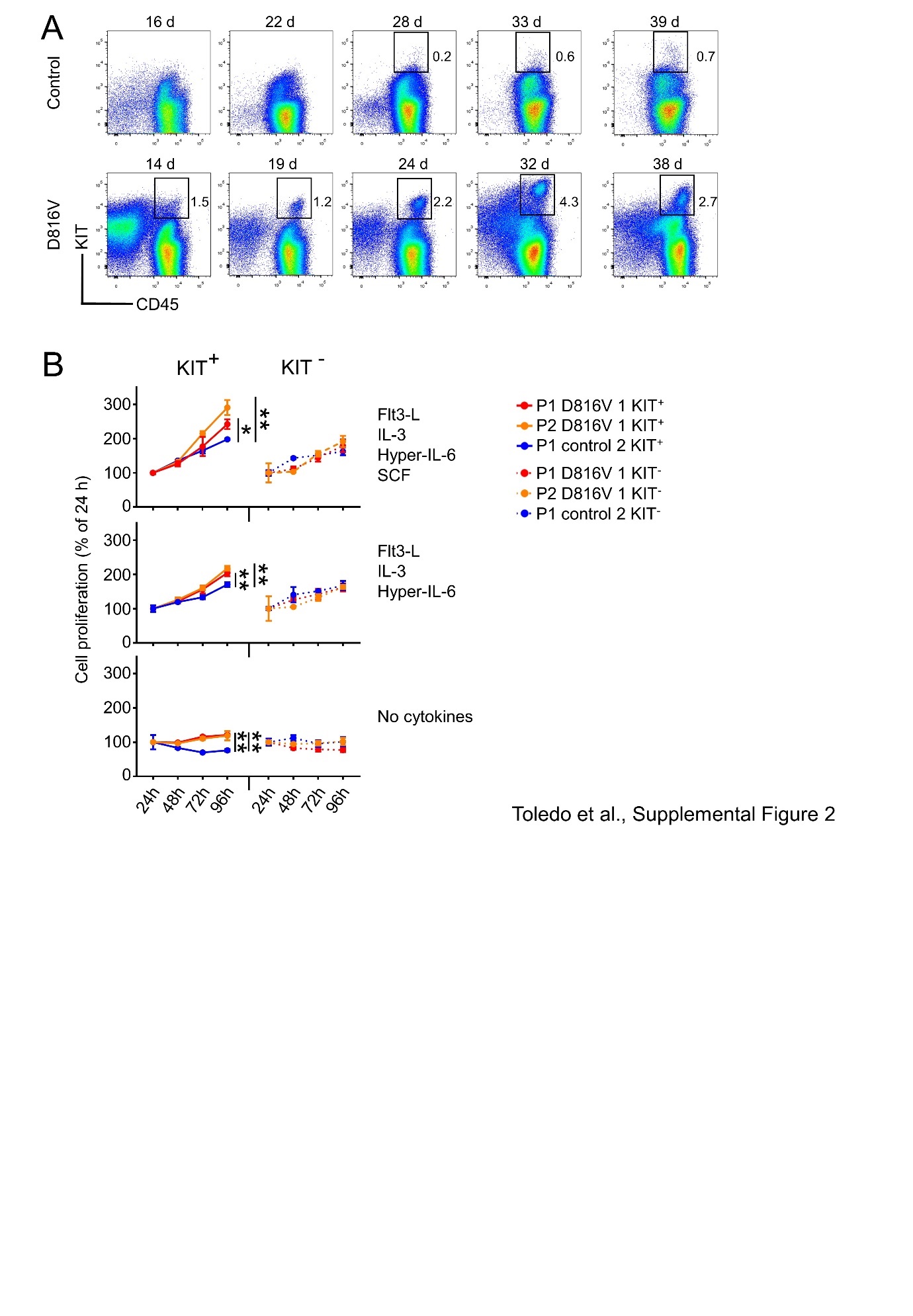

**Supplemental Figure 3**

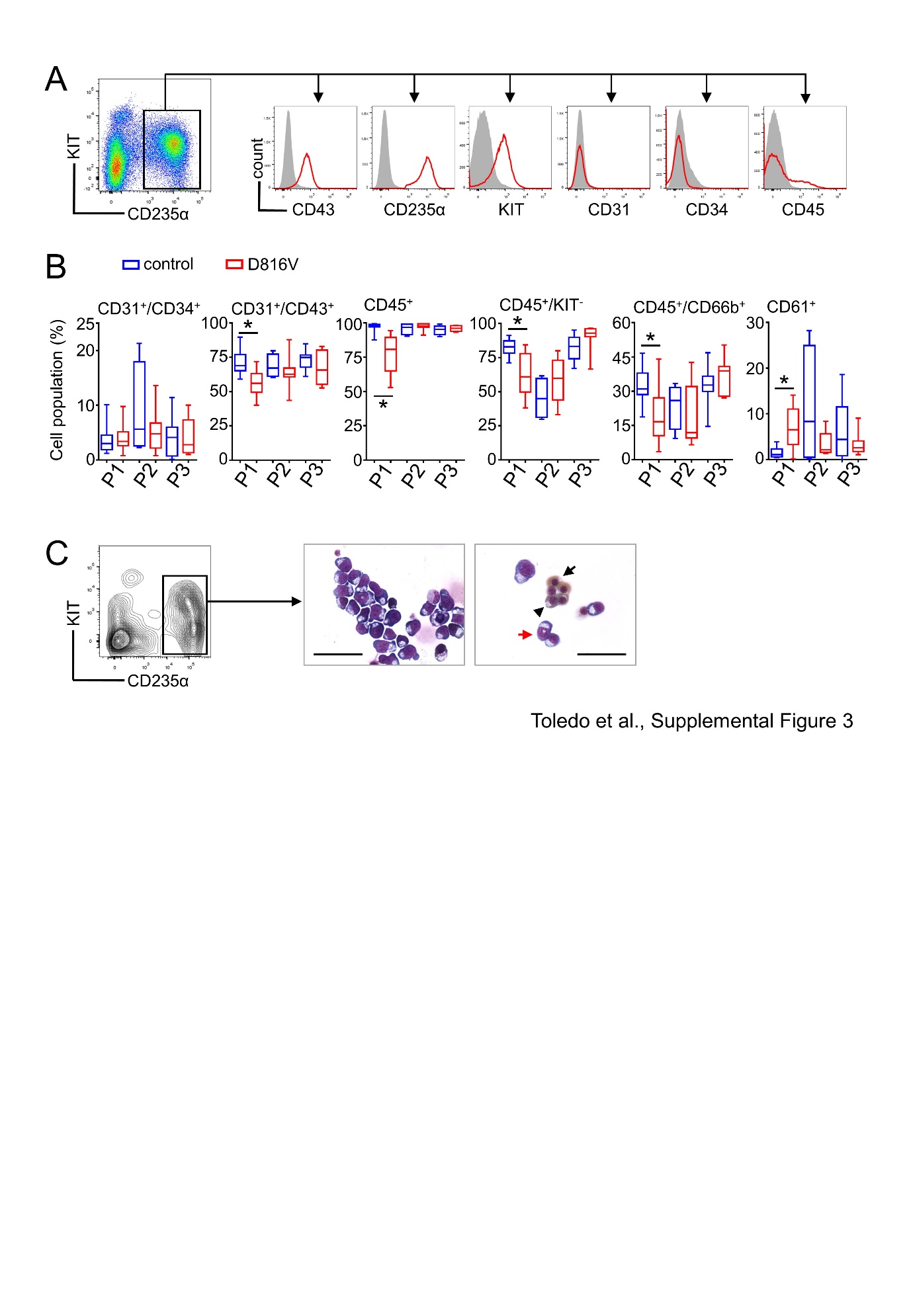

**Supplemental Figure 4**

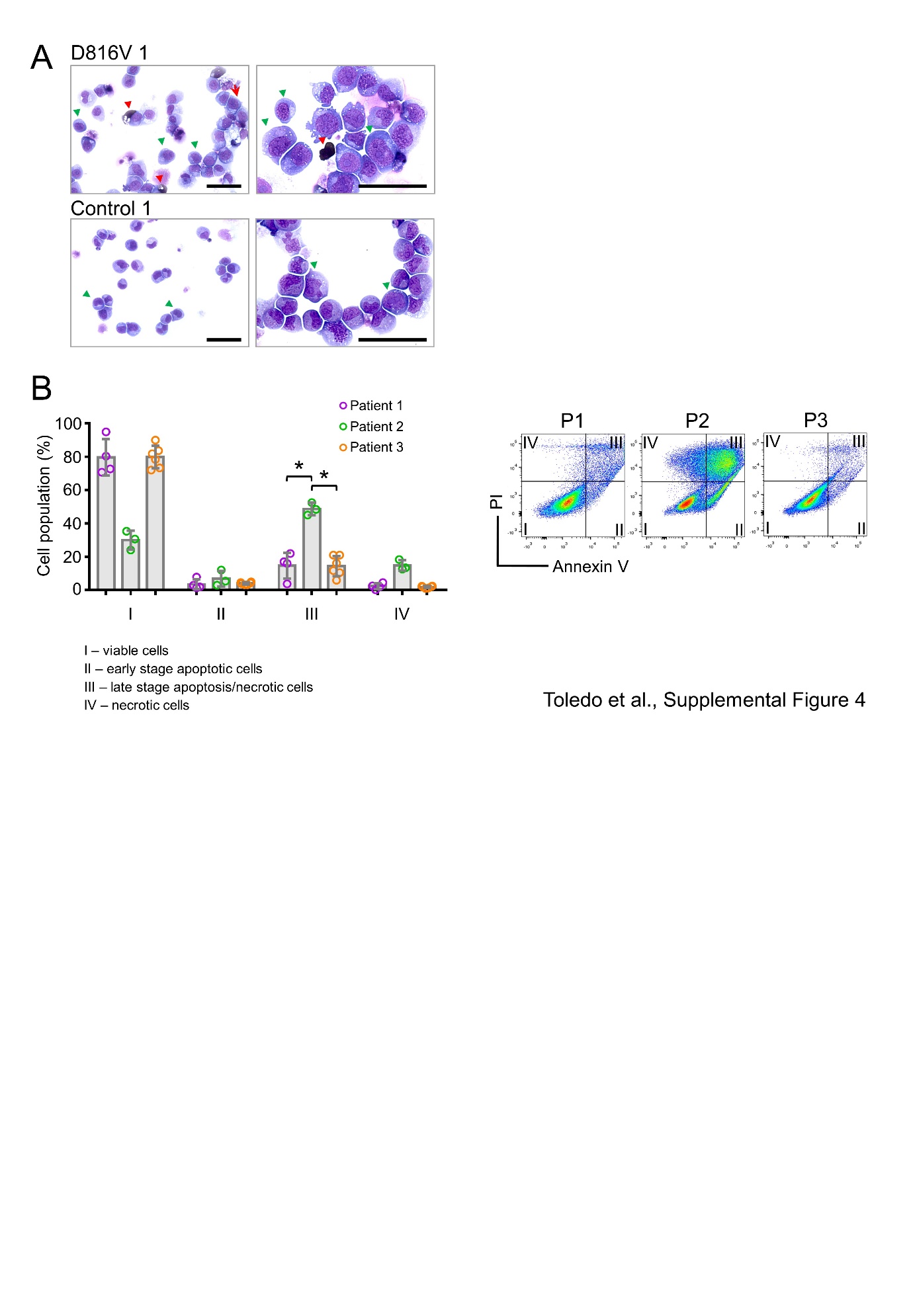

**Supplemental Figure 5**

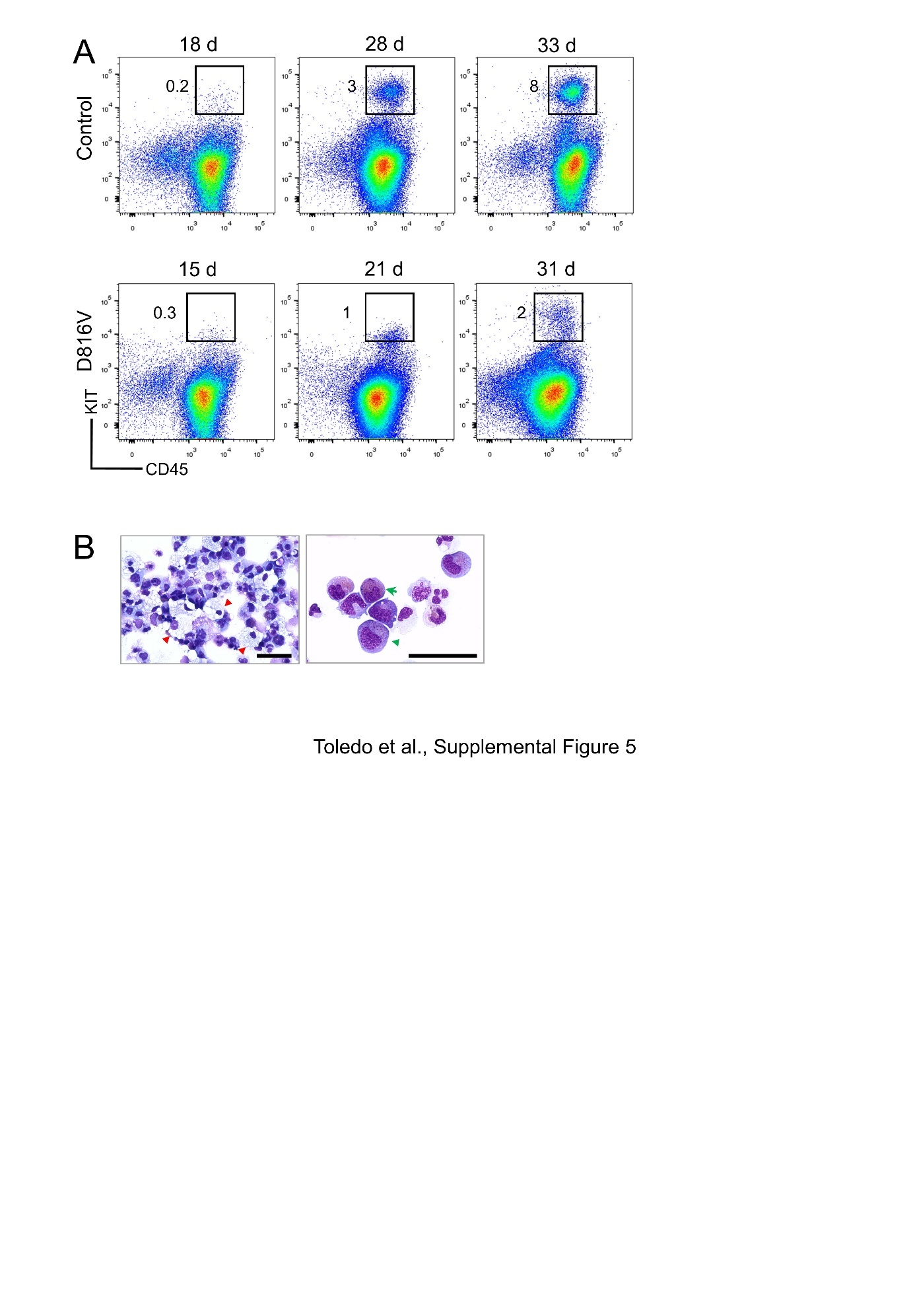

**Supplemental Figure 6**

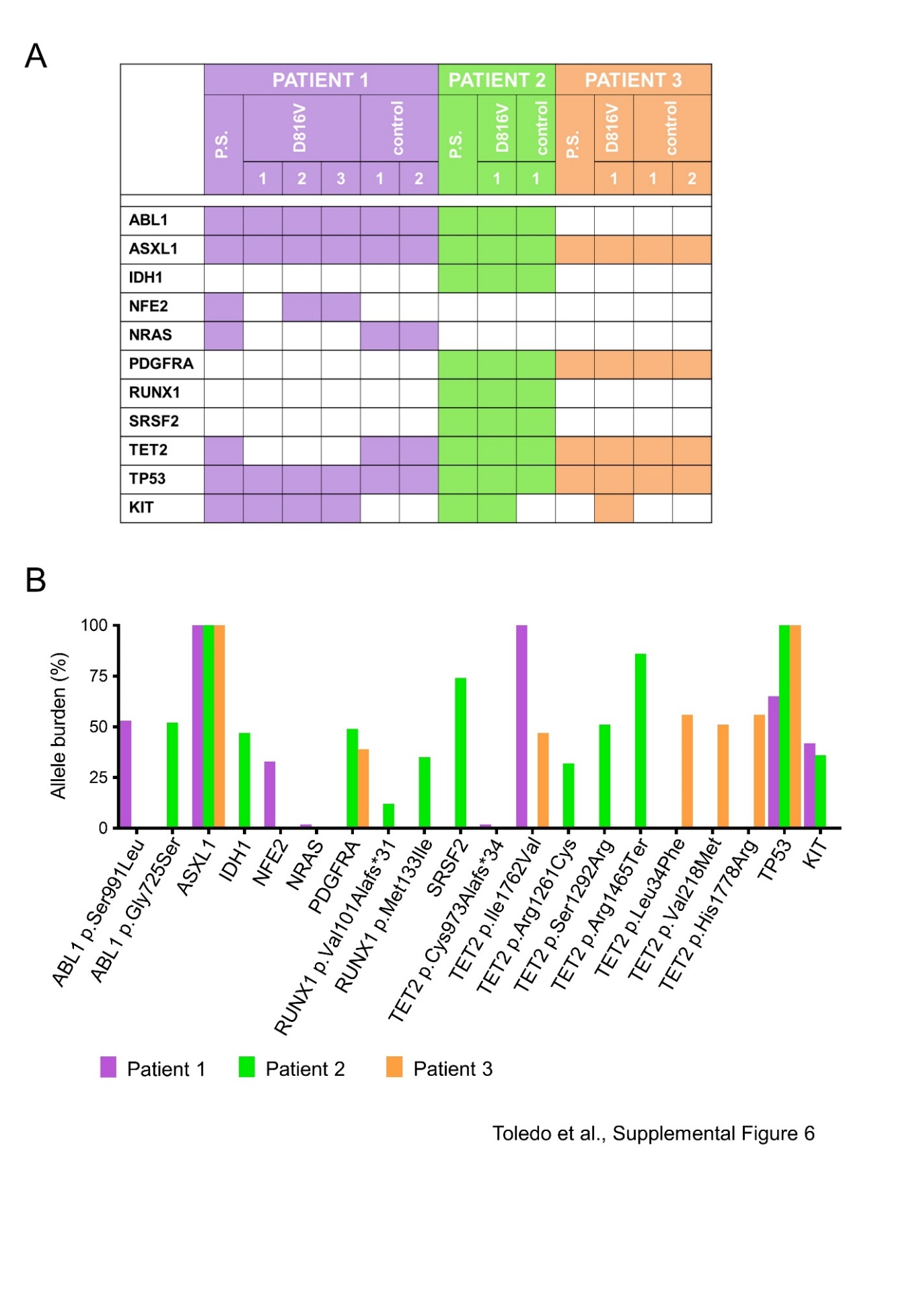

**Supplemental Figure 7**

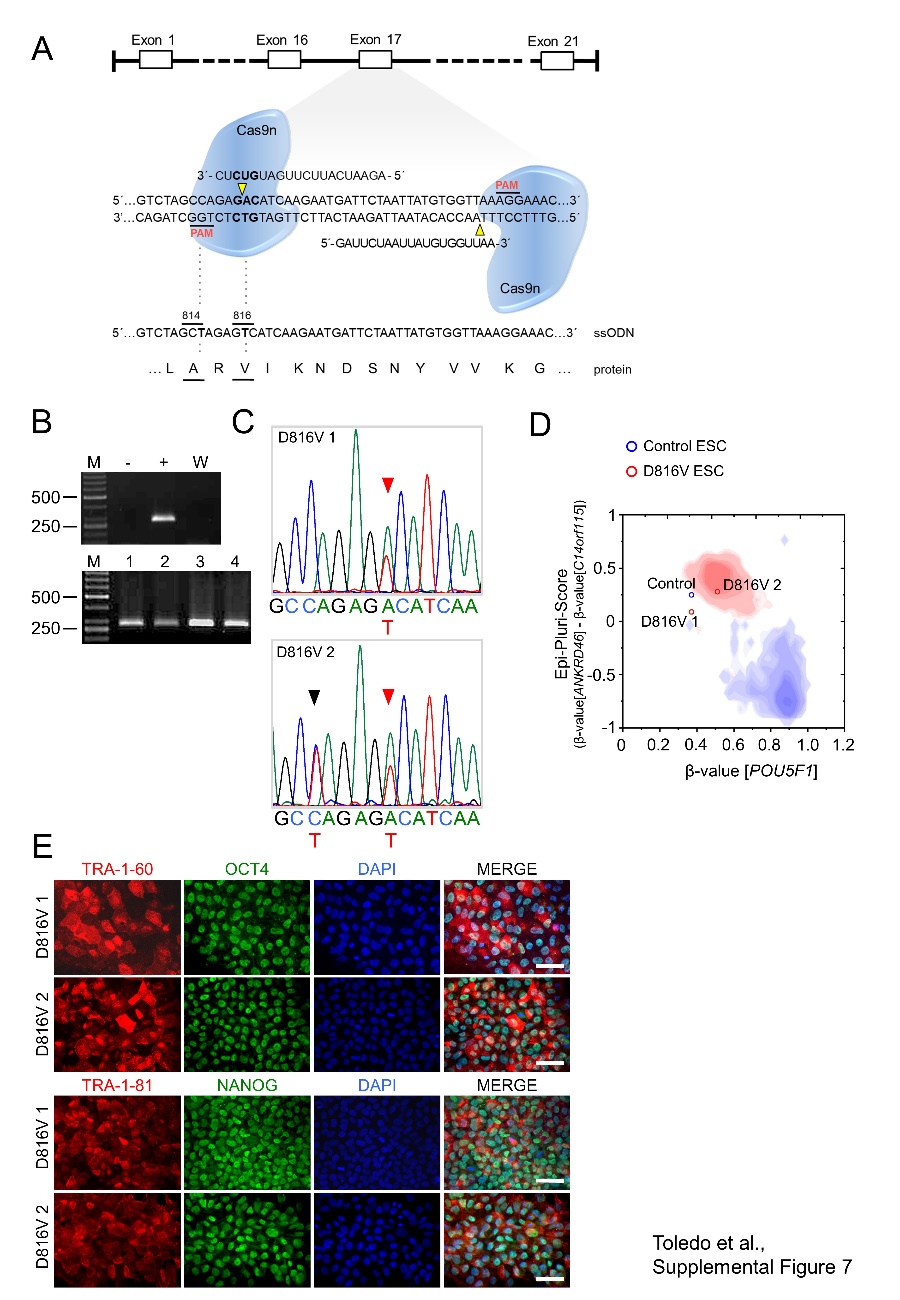

**Supplemental Figure 8**

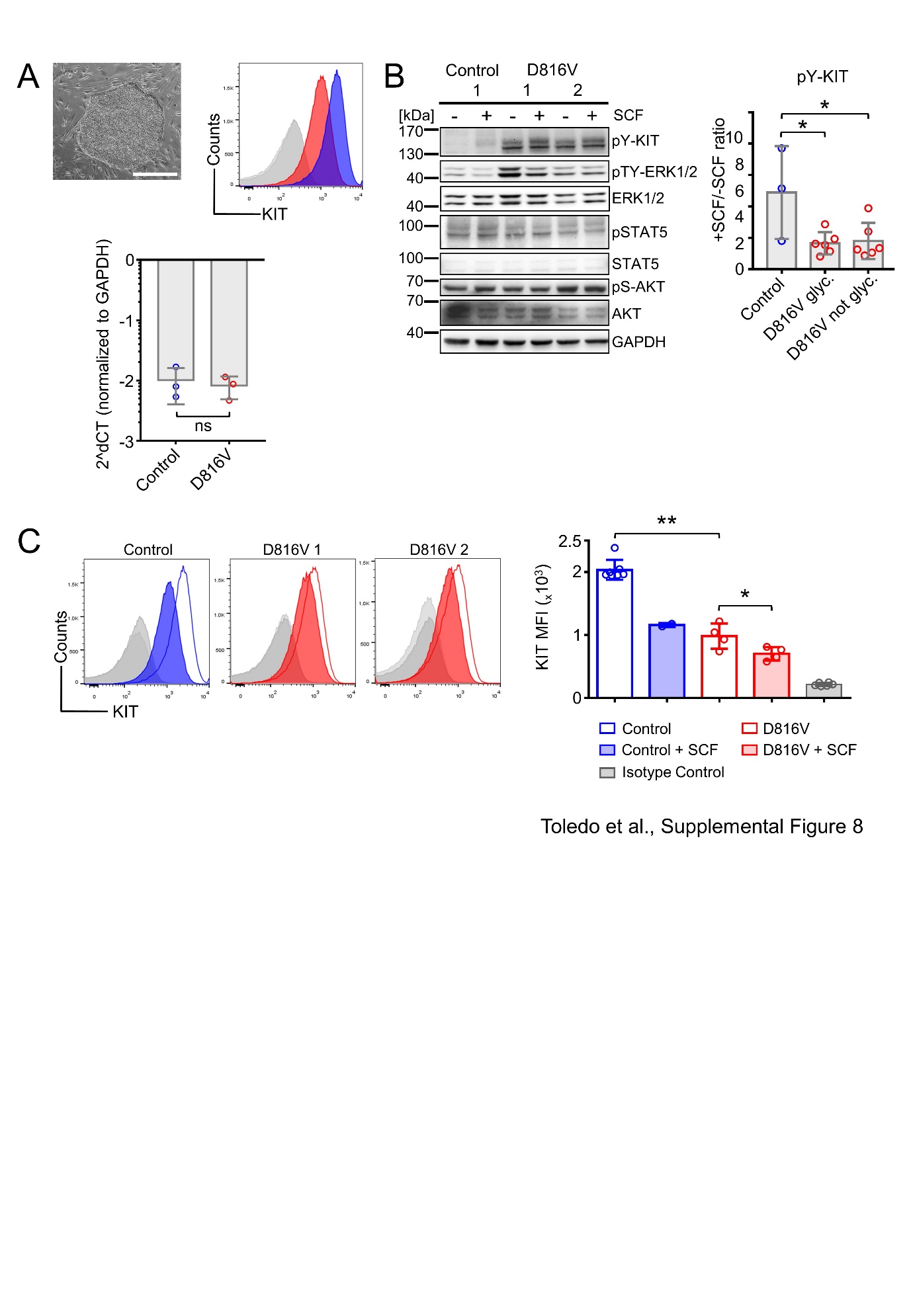

**Supplemental Figure 9**
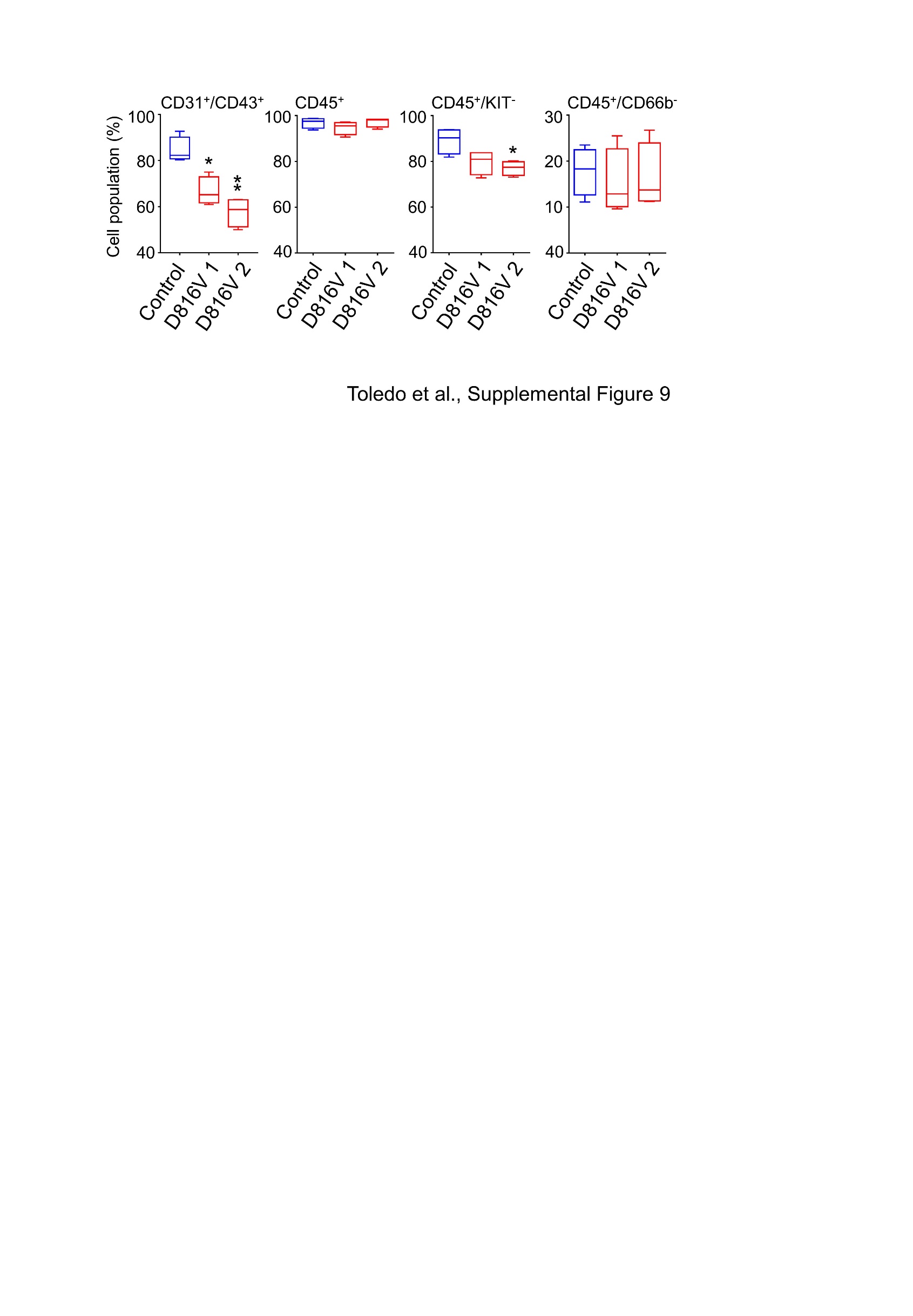

**Supplemental Figure 10**

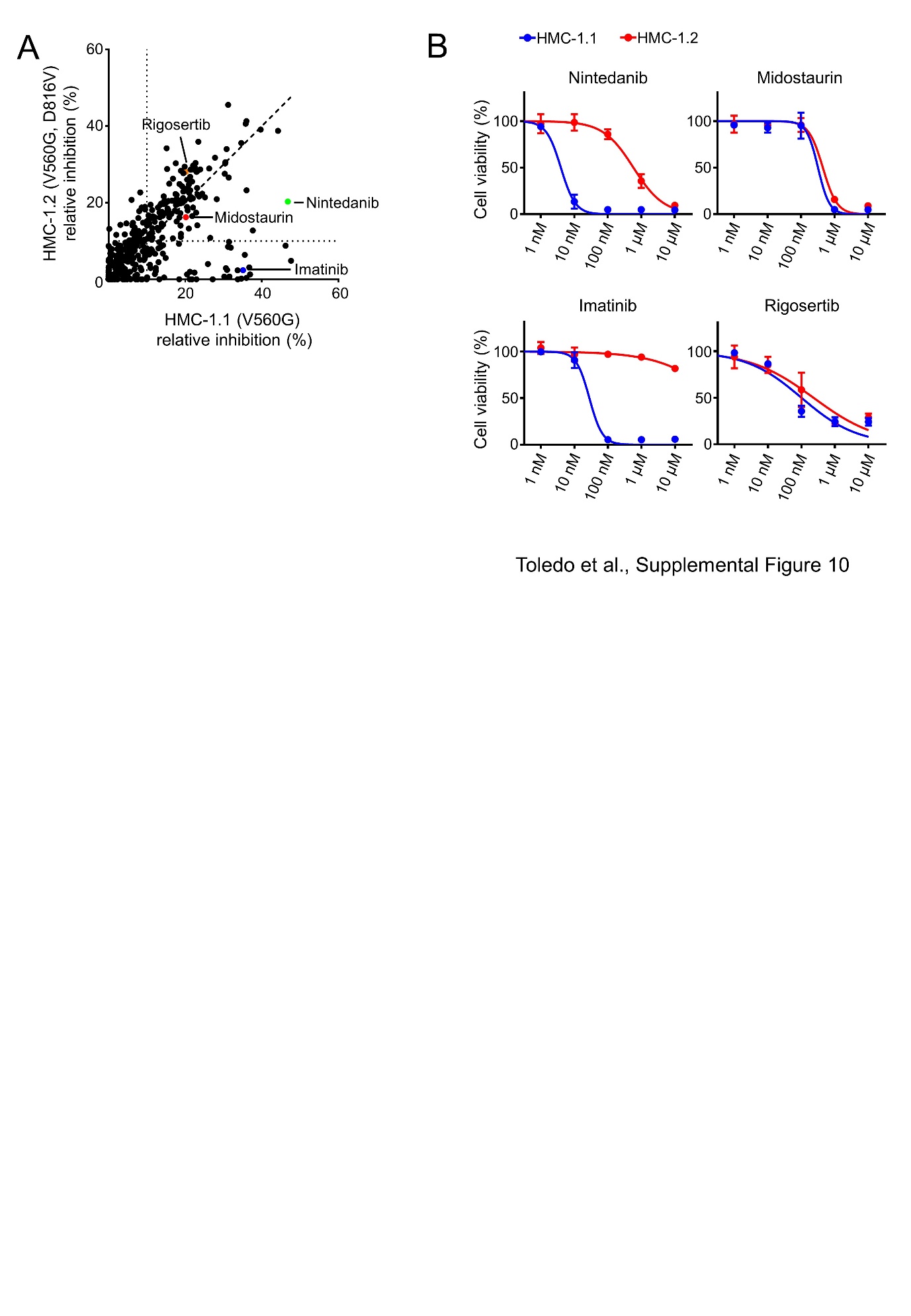

**Supplemental Figure 11**

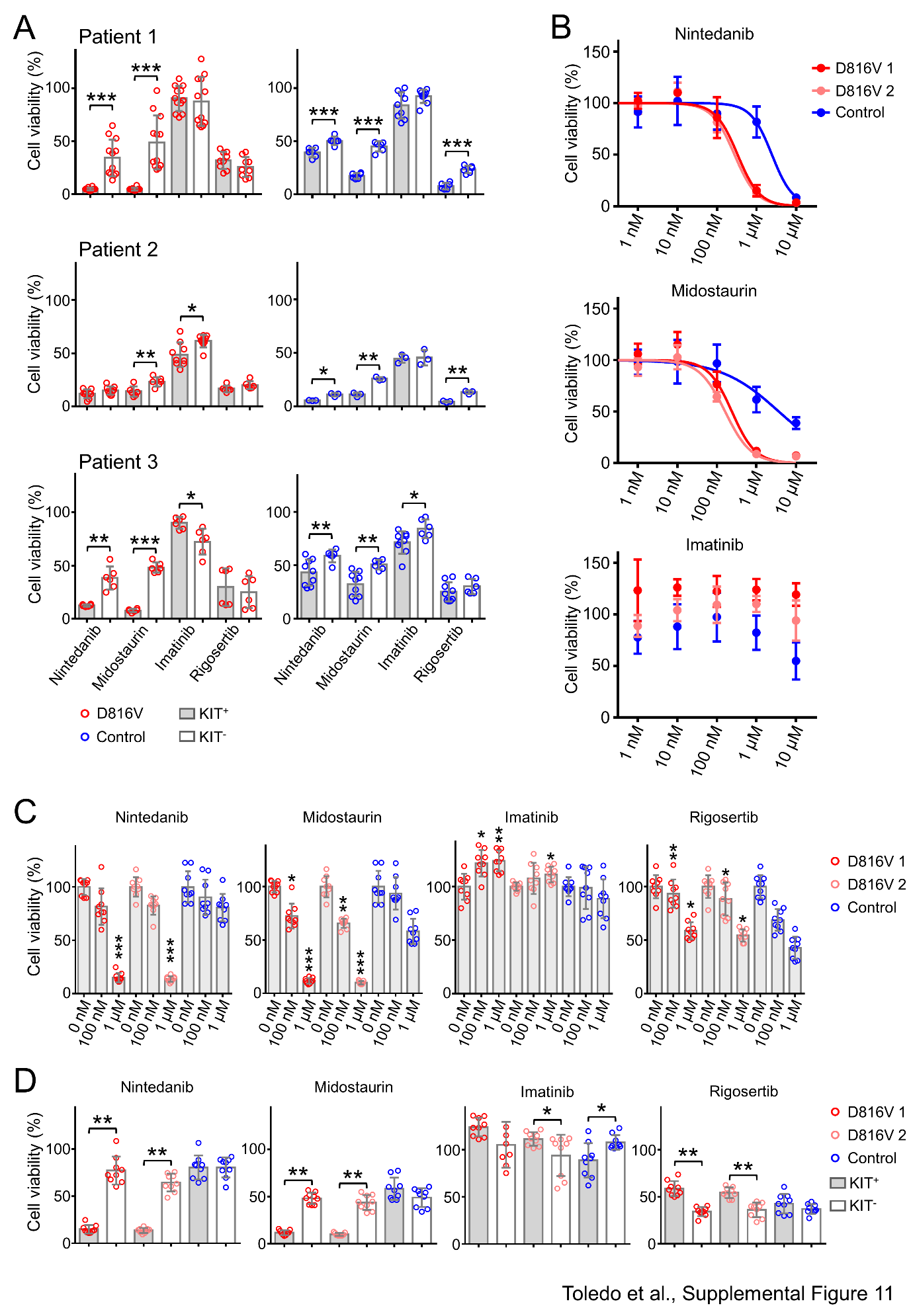

**Supplemental Figure 12**

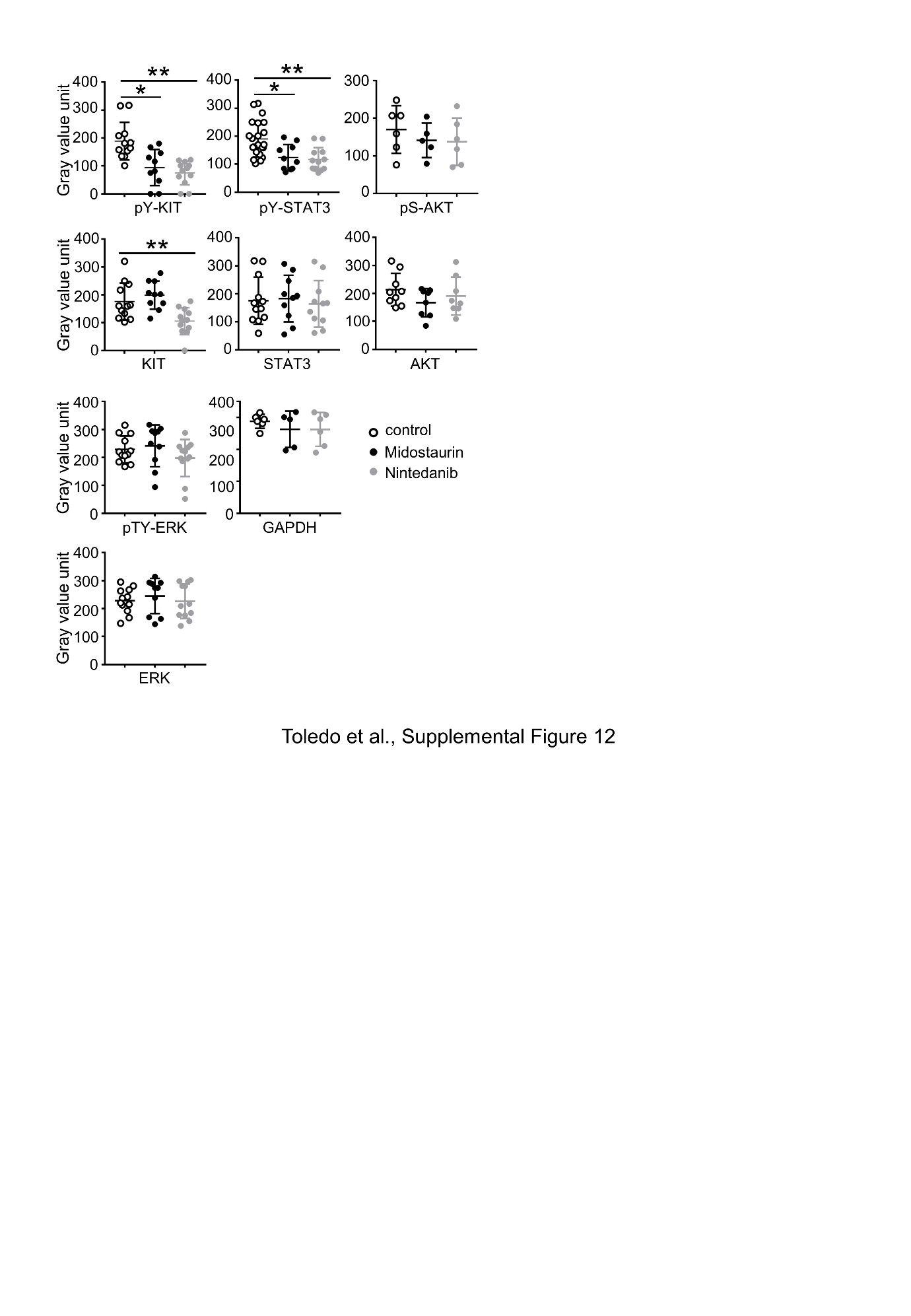

**Supplemental Figure 13**

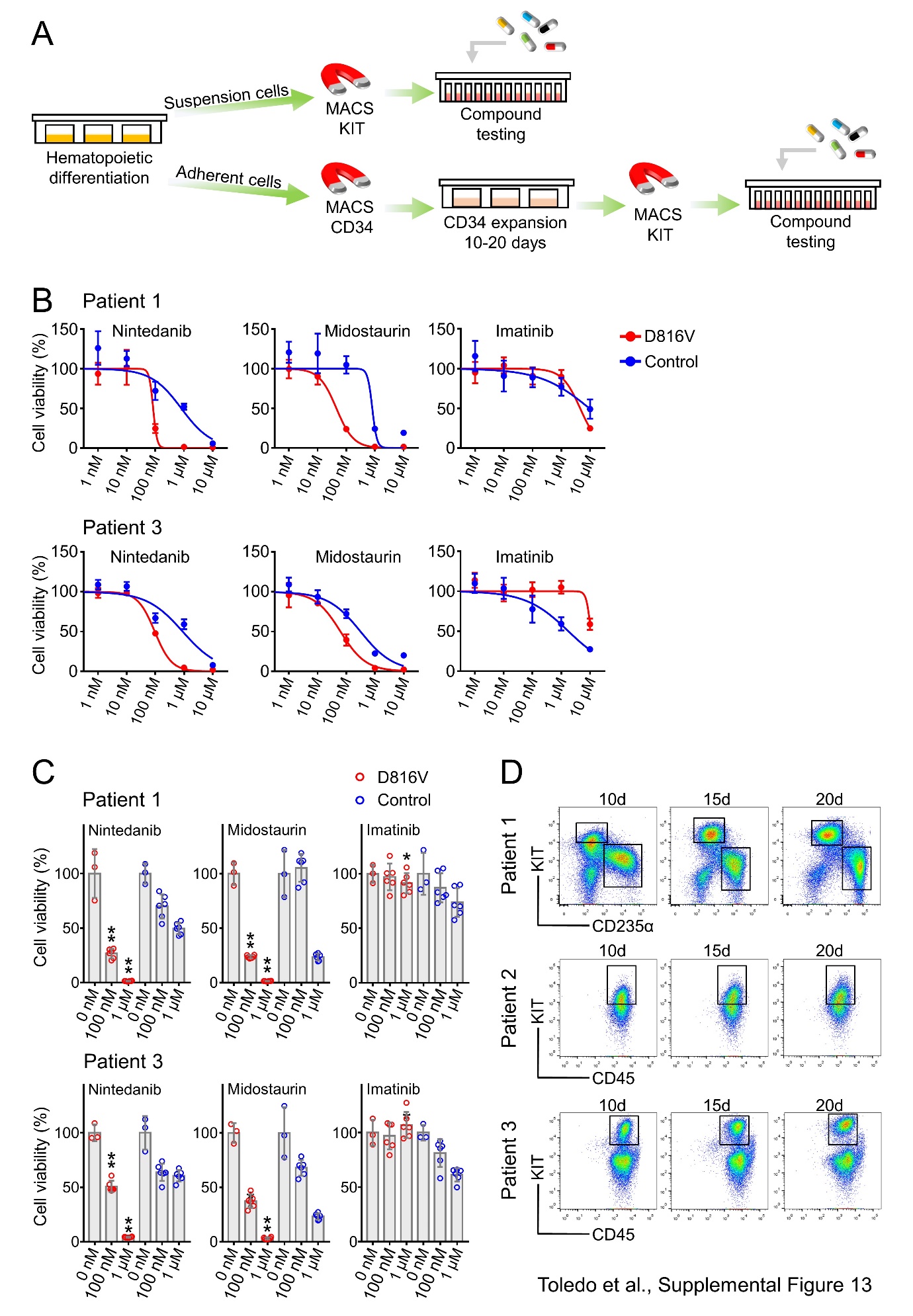

**Supplemental Figure 14**

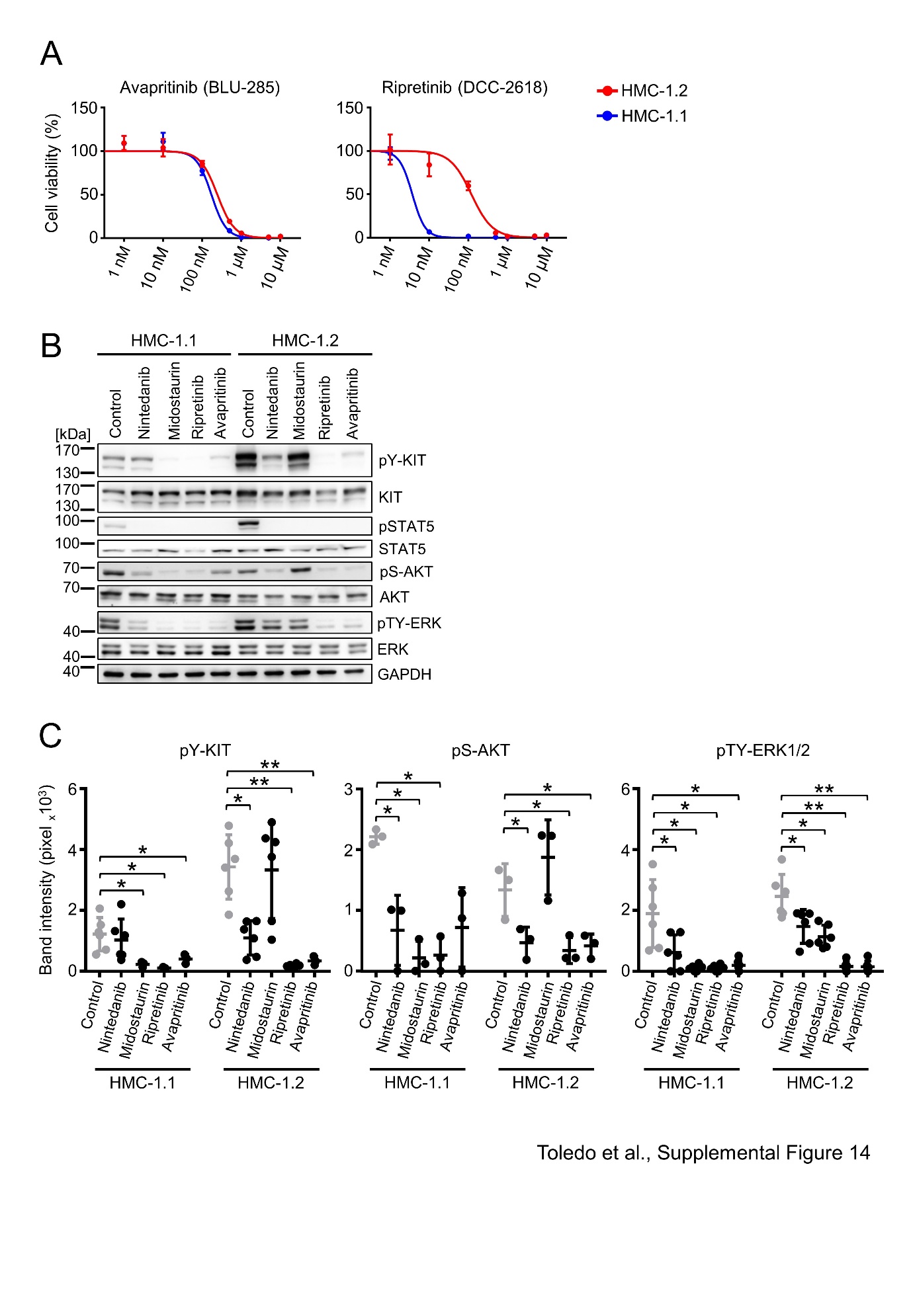

**Supplemental Figure 15**

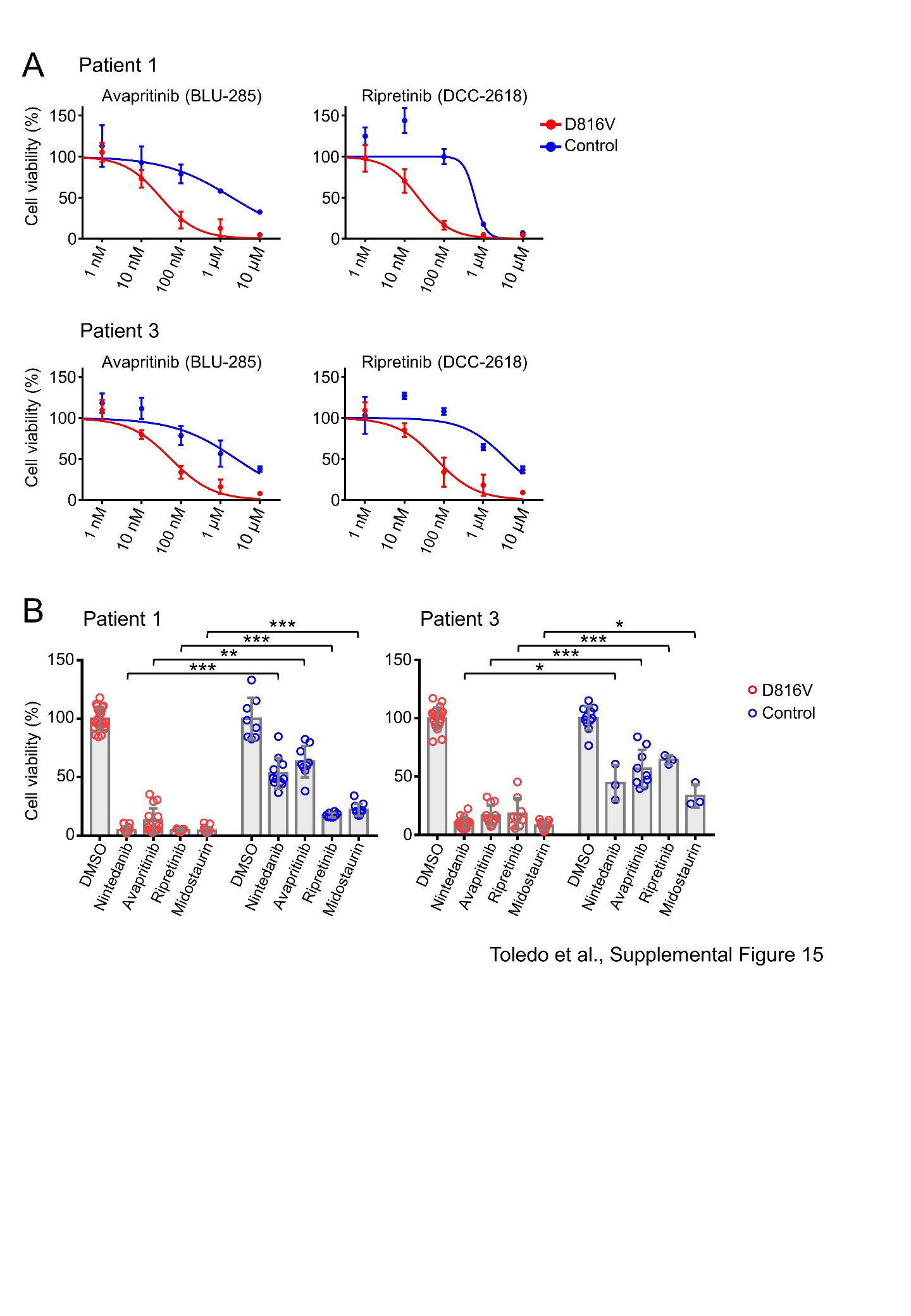

**Supplemental Figure 16**

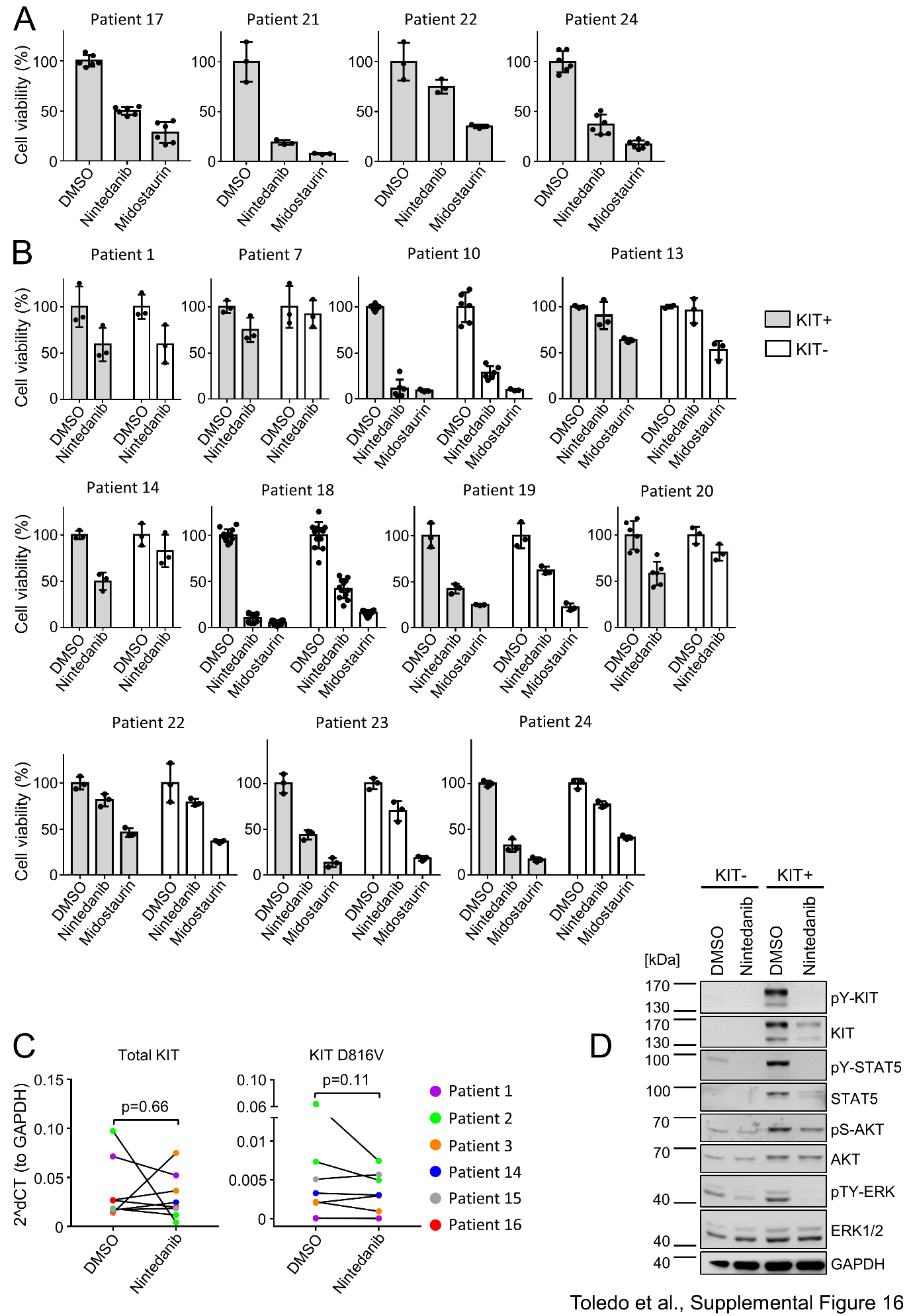

**Supplemental Figure 17**

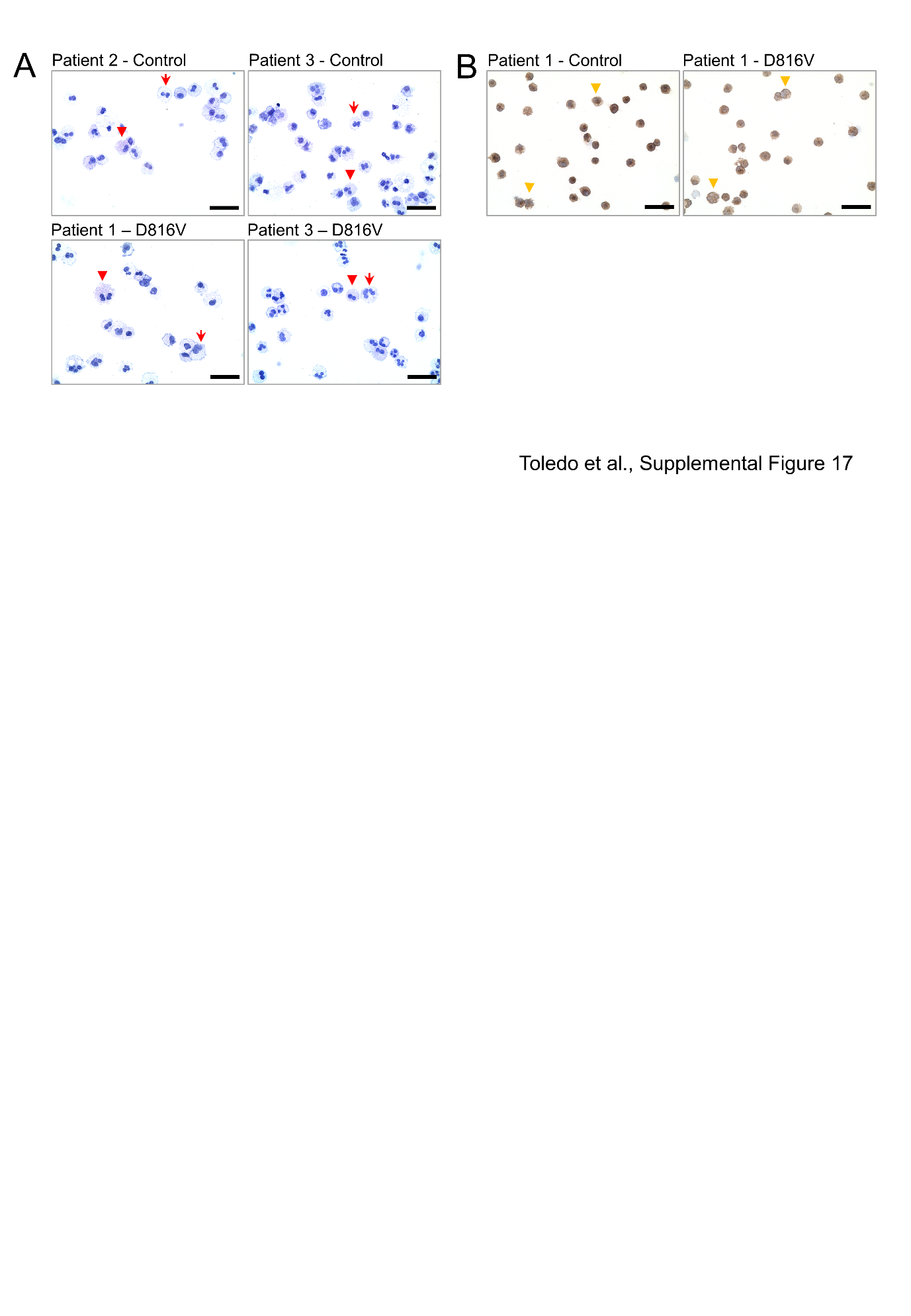

**Supplemental Figure 18**

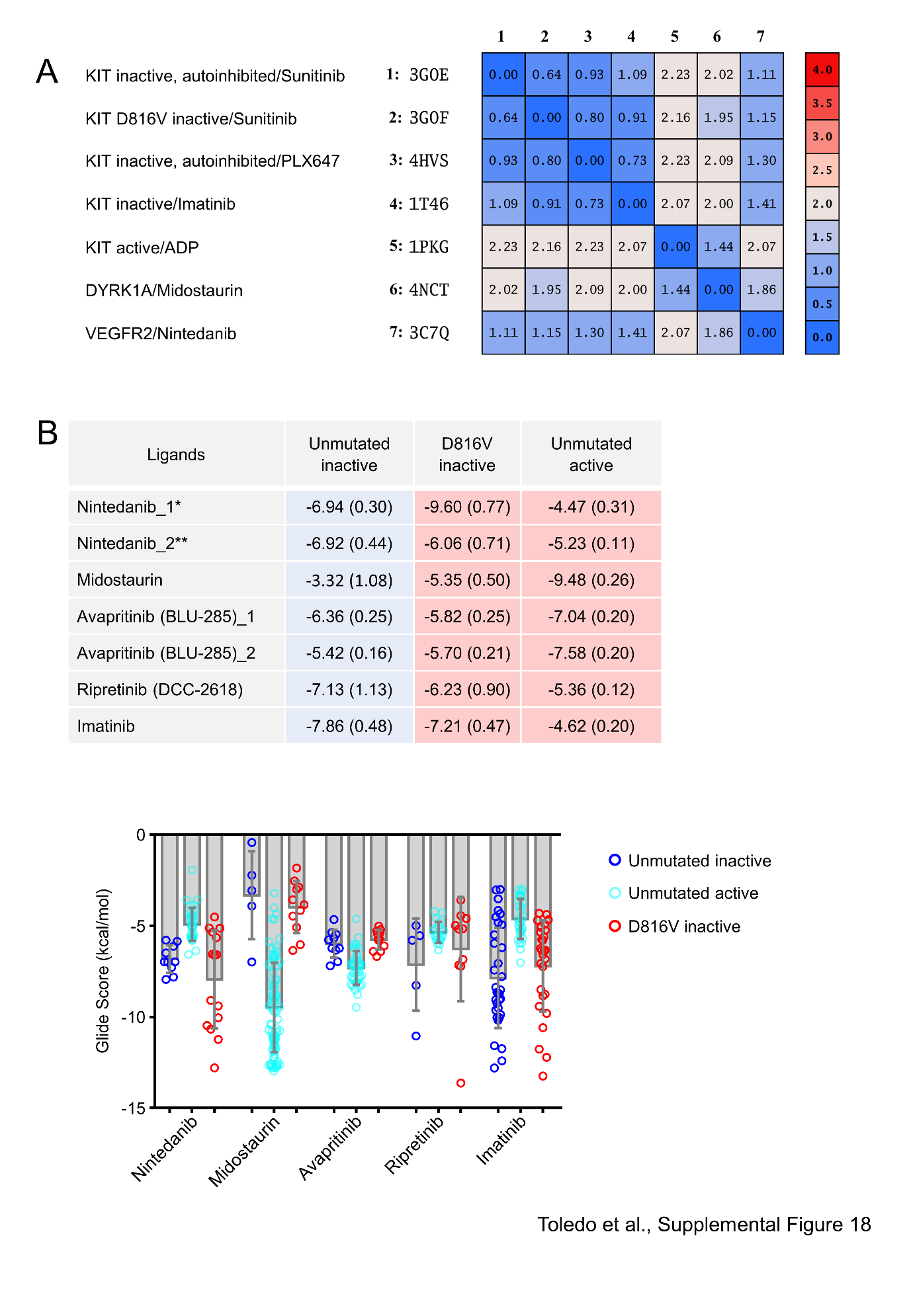

**Supplemental Figure 19**

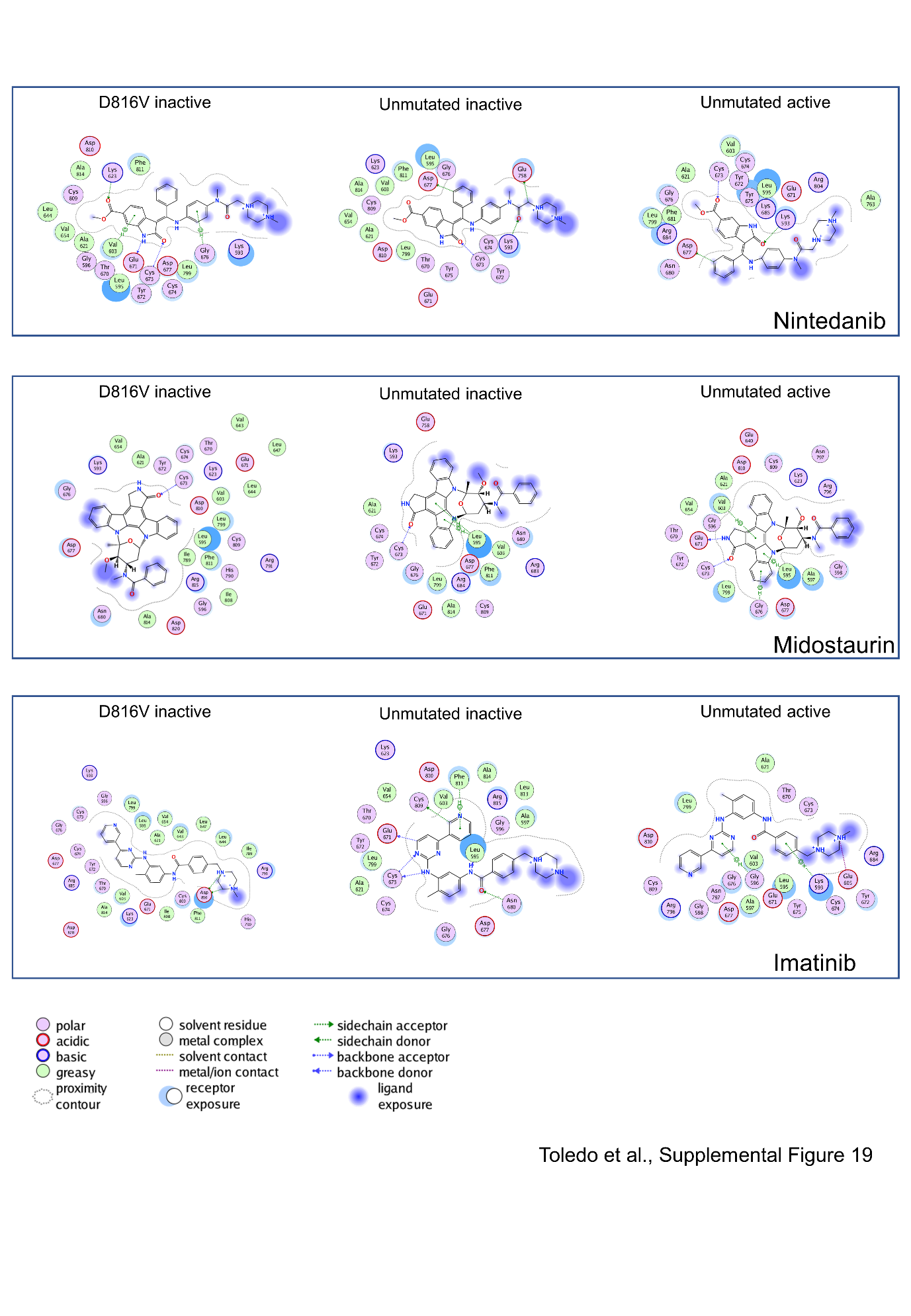

**Supplemental Figure 20**

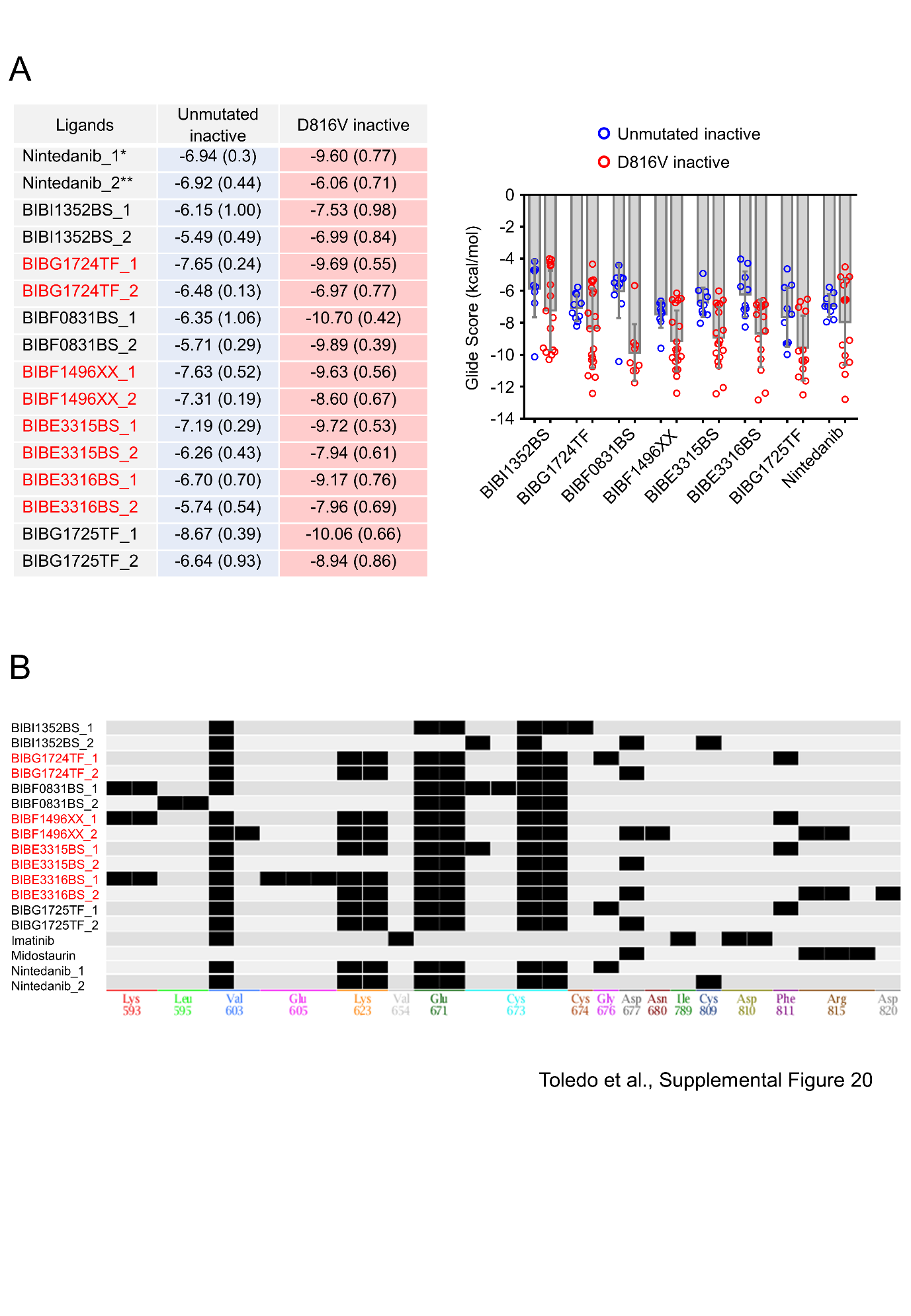
